# The practical impact of numerical variability on structural MRI measures of Parkinson’s disease

**DOI:** 10.64898/2026.01.09.698203

**Authors:** Yohan Chatelain, Andrzej Sokołowski, Madeleine Sharp, Jean-Baptiste Poline, Tristan Glatard

## Abstract

Numerical variability is rarely quantified in neuroimaging despite many measures relying on subtle morphometric differences across individuals. We instrumented FreeSurfer 7.3.1, a widely used neuroimaging pipeline, to simulate numerical differences across computational environments, and used it to measure numerical variability in MRI analyses of Parkinson’s disease patients and controls. In multiple cortical and subcortical regions, numerical variation reached nearly one-third of the population variability, altering statistical conclusions about group differences and clinical associations. To assess the impact of numerical noise in existing studies, we developed a practical tool that estimates the Numerical-Population Variability Ratio (NPVR) in a study, and propagates the resulting numerical variability to common statistics and associated p-values. By applying this framework to thirteen previously published studies reporting MRI measures in Parkinson’s disease, we quantified the probability of numerically induced false positives and false negatives in the literature, highlighting a substantial impact of numerical variability on MRI measures of Parkinson’s disease with an average significance-flip probability of 5% for cross-sectional studies and 10% for longitudinal studies. These results underscore the importance of systematically evaluating numerical stability in neuroimaging and provide a practical framework to do so.

## 1 Introduction

While Magnetic Resonance Imaging (MRI) remains a promising source of biomarkers for a variety of brain-related conditions, the reproducibility of MRI-derived measures across analytical conditions has been challenged in multiple contexts over the past few years. In structural MRI, cortical surface analyses have been shown to be substantially affected by software, parcellation, and quality control [1]. In functional MRI, different research teams analyzing the data have found only moderately consistent results [2]. In diffusion MRI, variability among white matter bundle segmentation protocols was found to be comparable to variability across subjects [3]. The reliability of MRI-derived measures critically depends on a better understanding and characterization of the impacts of analytical variability, including data selection, analytical decisions, tool selection, computational infrastructure, and numerical state [4, 5].

Among these sources of analytical variability, numerical variability has been shown to have a measurable impact on MRI analyses [6, 7] but remains understudied, mainly due to the practical challenges of quantifying its effects. Numerical variability arises from rounding and truncation errors associated with the use of limited-precision numerical formats, such as the widespread IEEE-754 standard for floating-point arithmetic [8]. Numerical errors manifest slightly differently across computational platforms (hardware, operating systems, or library versions) and these differences sometimes accumulate and amplify across computational analyses, eventually leading to measurable differences in final outputs [9, 10, 11, 12, 13, 14]. Such issues occur particularly within high-dimensional optimization processes such as linear and non-linear image registration, or the training of deep learning models [15].

The implications of numerical variability for clinical measures remain largely unknown. Previous studies measured its impact on image preprocessing but did not consider downstream statistical analyses. The first aim of this study was to quantify the impact of numerical variability in structural MRI analyses of Parkinson’s disease (PD), where robust MRI measures of the disease have yet to be identified. We conducted typical cross-sectional and longitudinal analyses of structural MRI data of PD participants, measuring numerical variability through an experimental stochastic arithmetic approach.

Building on these observations, we developed an analytical framework and associated tools to rapidly assess the numerical quality of structural MRI analyses reported in the published literature, opening the possibility to conduct large-scale impact evaluations of numerical variability. By making numerical variability evaluation accessible, our framework and tool enhance transparency, support peer review, and promote more reliable statistical inference in neuroimaging. Applying this framework to the PD literature, we obtained the first estimates of the numerical quality of MRI analyses in Parkinson’s disease studies, highlighting a widespread impact of numerical variability on MRI measures of PD.

## 2 Results

### 2.1 Numerical variability alters statistical inference in MRI measures of PD

We assessed the impact of numerical variability on conclusions drawn from MRI analyses of Parkinson’s disease, focusing on two common analyses: (1) volumetric group differences between PD subjects and Healthy Controls (HC), and (2) partial correlations between regional volumes and motor evaluation scores measured with the MDS-Unified Parkinson’s Disease Rating Scale part 3 (UPDRS-III). For both, we conducted a cross-sectional analysis at baseline and a longitudinal analysis across two time points.

Analyses included the 112 PD-non-MCI and 89 HC participants from the PPMI dataset [16] who met inclusion criteria after quality control (Methods, §4.1; Table 2).

We processed all images for both time points using FreeSurfer 7.3.1 instrumented with Monte Carlo Arithmetic (MCA) [17] to introduce machine-level numerical noise and quantify numerical variability across repeated runs, yielding 26 valid perturbed realizations per participant after quality control (Methods, §4.3; Supplementary Table S4). For all analyses, the unperturbed (IEEE-754) result fell within the range of numerically perturbed results, supporting the validity of the perturbation approach (Supplementary Note S4).

For both group comparisons and correlation analyses, statistical outcomes varied substantially across the 26 Monte Carlo Arithmetic (MCA) repetitions (Figures 1 and 2). For subcortical volumes (14 regions; Figure 1), p-values crossed the 0.05 threshold as a result of numerical perturbation in 27% of all (regions, analysis) pairs, indicating frequent inconsistencies across MCA repetitions. For cortical thickness (68 regions; Figure 2), 21% of (regions, analysis) pairs were similarly unstable. These rates varied across metric types: cortical area exhibited the highest longitudinal variability (53% for ANCOVA), followed by cortical volume (53% for partial correlation), subcortical volume (36% for partial correlation), and cortical thickness (25% for partial correlation), consistent with a metric-specific dependence on numerical precision (see Supplementary Table S5 and Supplementary Note S3 for region-level details). Test statistics (r-values and F-values) also exhibited substantial variability across repetitions (see Supplementary Note S4). The greater variance of longitudinal compared with cross-sectional analyses was confirmed statistically by a one-sided Ansari-Bradley test contrasting the two (Supplementary Note S3). This variability is consistent with the low numerical precision measured in FreeSurfer results: cross-sectional analyses revealed fewer than 2 significant digits for most subcortical and cortical regions (see Supplementary Note S2). These fluctuations demonstrate that numerical noise alone can alter downstream statistical inference in structural MRI analyses of PD.

**Figure 1:**
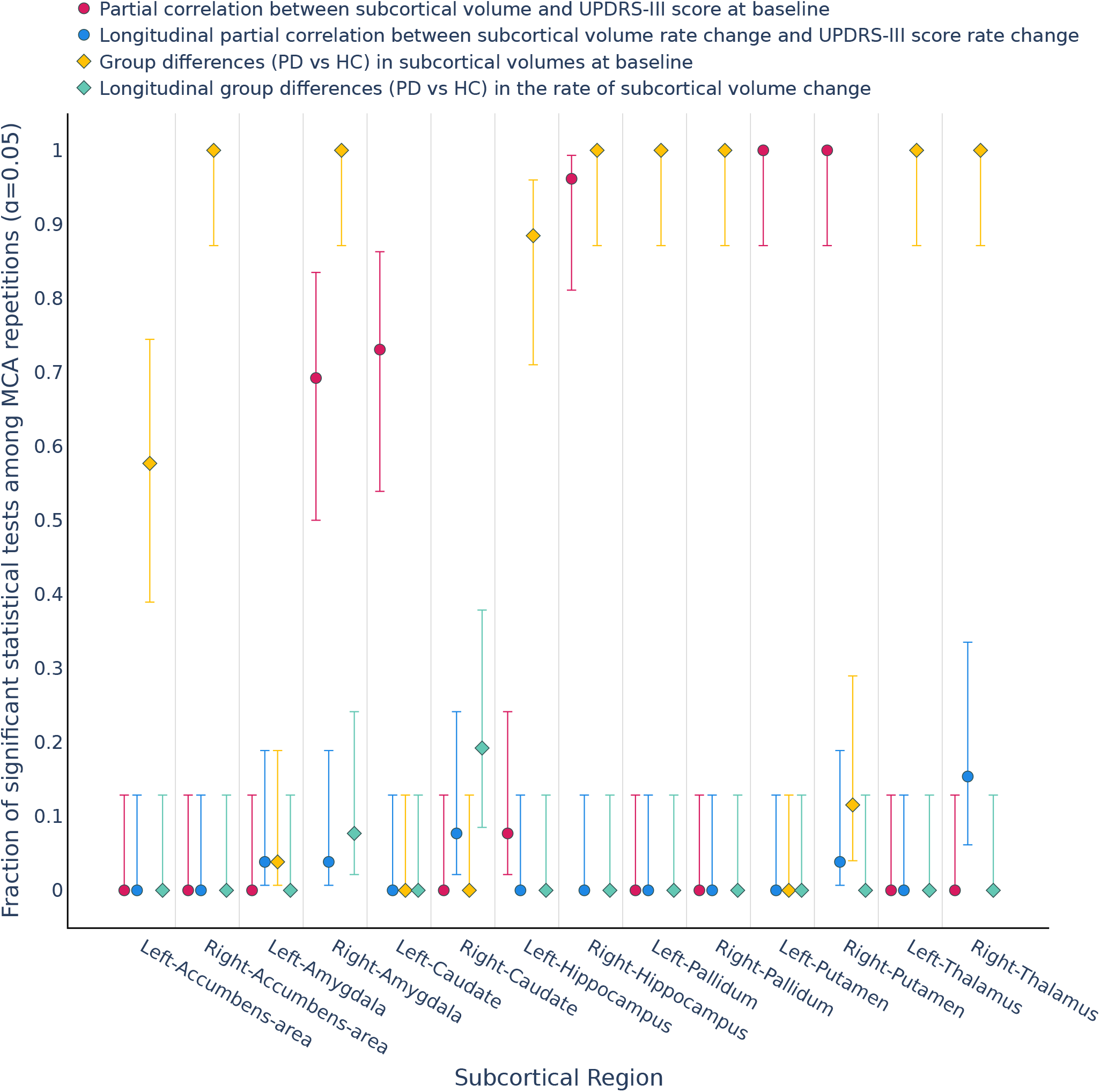
Proportion of significant tests (*p <* 0.05, uncorrected) for subcortical volumes across 26 numerical perturbations. Confidence intervals were computed using the Wilson score interval at 95% confidence level.

**Figure 2:**
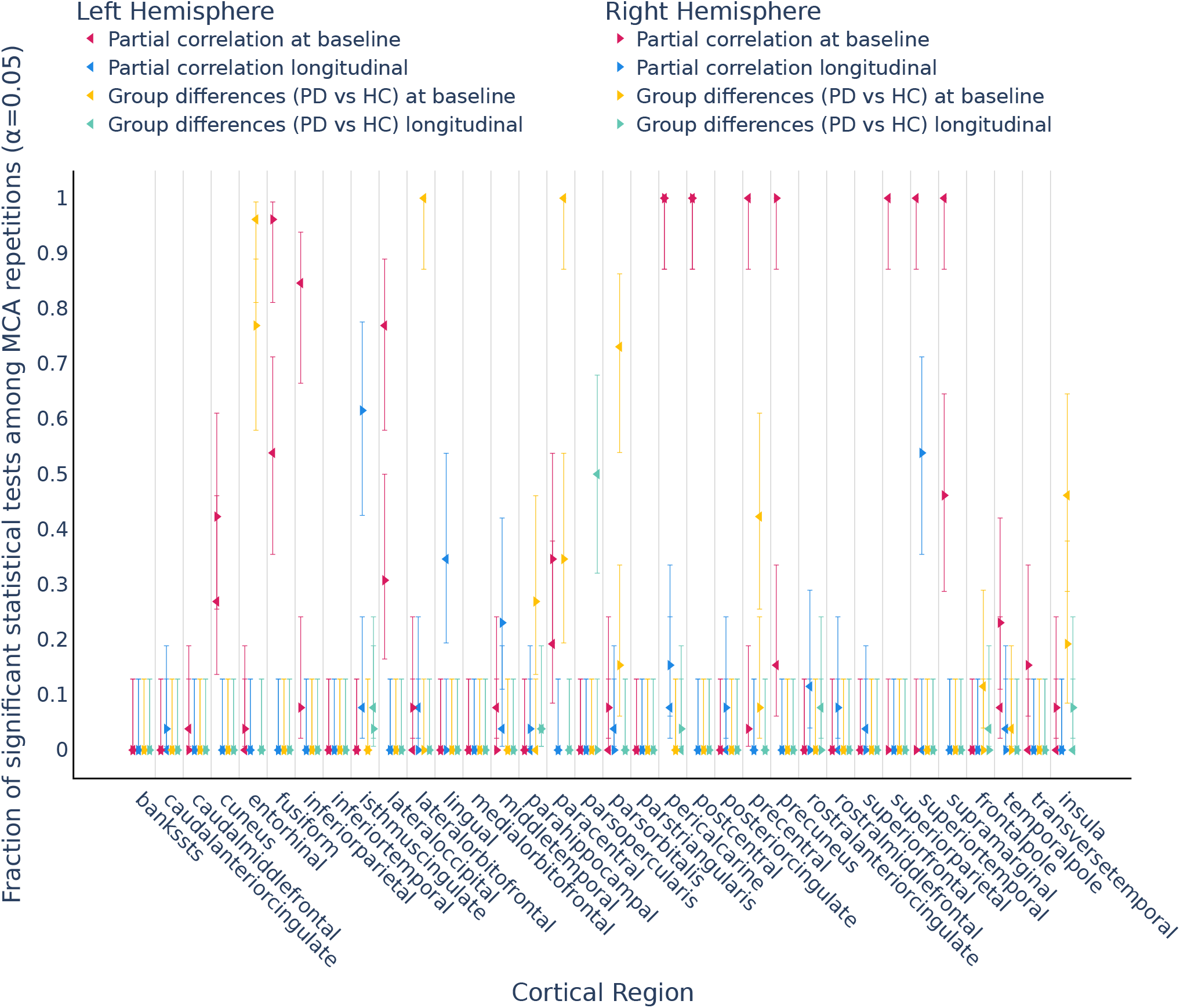
Proportion of significant tests (*p <* 0.05, uncorrected) for cortical thickness across 26 numerical perturbations. Confidence intervals were computed using the Wilson score interval at 95% confidence level.

### 2.2 A practical model to quantify the impact of numerical variability

While the previous results demonstrate that numerical variability can alter statistical inference in MRI analyses of PD, routinely conducting computationally expensive Monte Carlo evaluations is impractical for most studies. To address this issue, we developed a closed-form analytical model that captures numerical variability using only standard summary statistics. The core of this framework is the Numerical-Population Variability Ratio (*ν*_npv_), defined as the ratio between numerical variability (*σ*_num_; Eq. 1) and population variability (*σ*_pop_; Eq. 2):

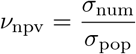

The *ν*_npv_ metric standardizes the quantification of numerical variability, enabling direct comparisons across different brain regions, software pipelines, and cohorts. Propagating this ratio through standard statistical estimators (Methods, §4.3.2) yields closed-form approximations for the numerical variability in common statistics (Table 1), numerically validated in Supplementary Note S5. This framework establishes a direct link between a pipeline’s numerical variability (*ν*_npv_), the study’s sample size (*n*), and the resulting reliability of p-values and effect sizes. Crucially, because these formulas rely solely on summary statistics, they allow for the retrospective quality control of existing literature without requiring access to original raw data or costly re-computation.

**Table 1:**
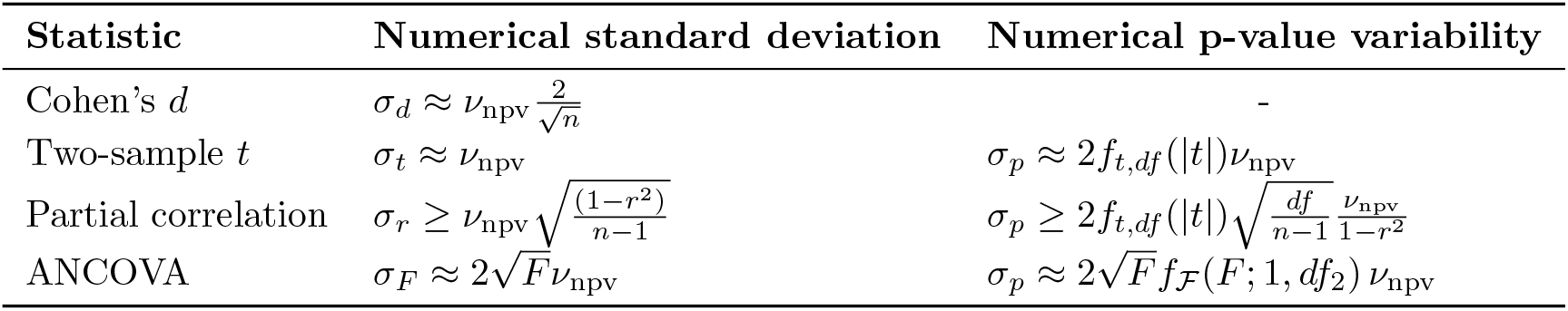
First-order numerical variability of common statistical tests under Monte Carlo Arithmetic perturbations. Cohen’s d formula assumes large and equal group sizes. *f*_*t,df*_ and *f*_*F*_ (*F* ; 1, *df*_2_) denote the probability density functions of the Student’s *t*-distribution with *df* degrees of freedom and the ℱ-distribution with (1, *df*_2_) degrees of freedom, respectively. The *p*-value approximation for the partial correlation uses *t* = *r*(*df /*(1 − *r*^2^))^1/2^.

**Table 2:** Participant characteristics. Values represent mean ± standard deviation. PD = Parkinson’s disease; MCI = mild cognitive impairment; UPDRS = Unified Parkinson’s Disease Rating Scale. The PD-non-MCI longitudinal subset corresponds to participants with available follow-up MRI and disease severity scores available.

| Cohort | HC | PD-non-MCI |
| --- | --- | --- |
| $n$ | 89 | 112 |
| Age (years) | $60.7 \pm 9.7$ | $60.6 \pm 8.9$ |
| Age range | 30.6 — 79.8 | 39.2 — 78.3 |
| Gender (male,%) | 47 (52.8%) | 74 (66.1%) |
| Education (years) | $16.7 \pm 3.4$ | $16.0 \pm 3.1$ |
| UPDRS-III OFF baseline | — | $23.3 \pm 10.0$ |
| UPDRS-III OFF follow-up | — | $25.6 \pm 11.2$ |
| Inter-visit interval (years) | $1.4 \pm 0.5$ | $1.4 \pm 0.6$ |

We applied this model to quantify the stability of the analyses reported in the previous section (Figure 3; Supplementary Figure S6). In cross-sectional baseline analyses, the average *ν*_npv_ was 0.191 for the PD group and 0.176 for HC. A permutation test found no difference surviving Bonferroni correction across the four metrics (area *p* = 0.734, thickness *p* = 0.033, volume *p* = 0.371, subcortical volume *p* = 0.646; Bonferroni-corrected threshold *α* = 0.05*/*8 = 0.00625 over the eight metric × analysis tests; Supplementary Table S6). Applying the Cohen’s *d* variability formula (Table 1) to these *ν*_npv_ values reveals that suppressing numerical variability to a negligible level (*σ*_*d*_ ≤ 0.01) in cross-sectional studies would require approximately 1,340 participants.

**Figure 3:**
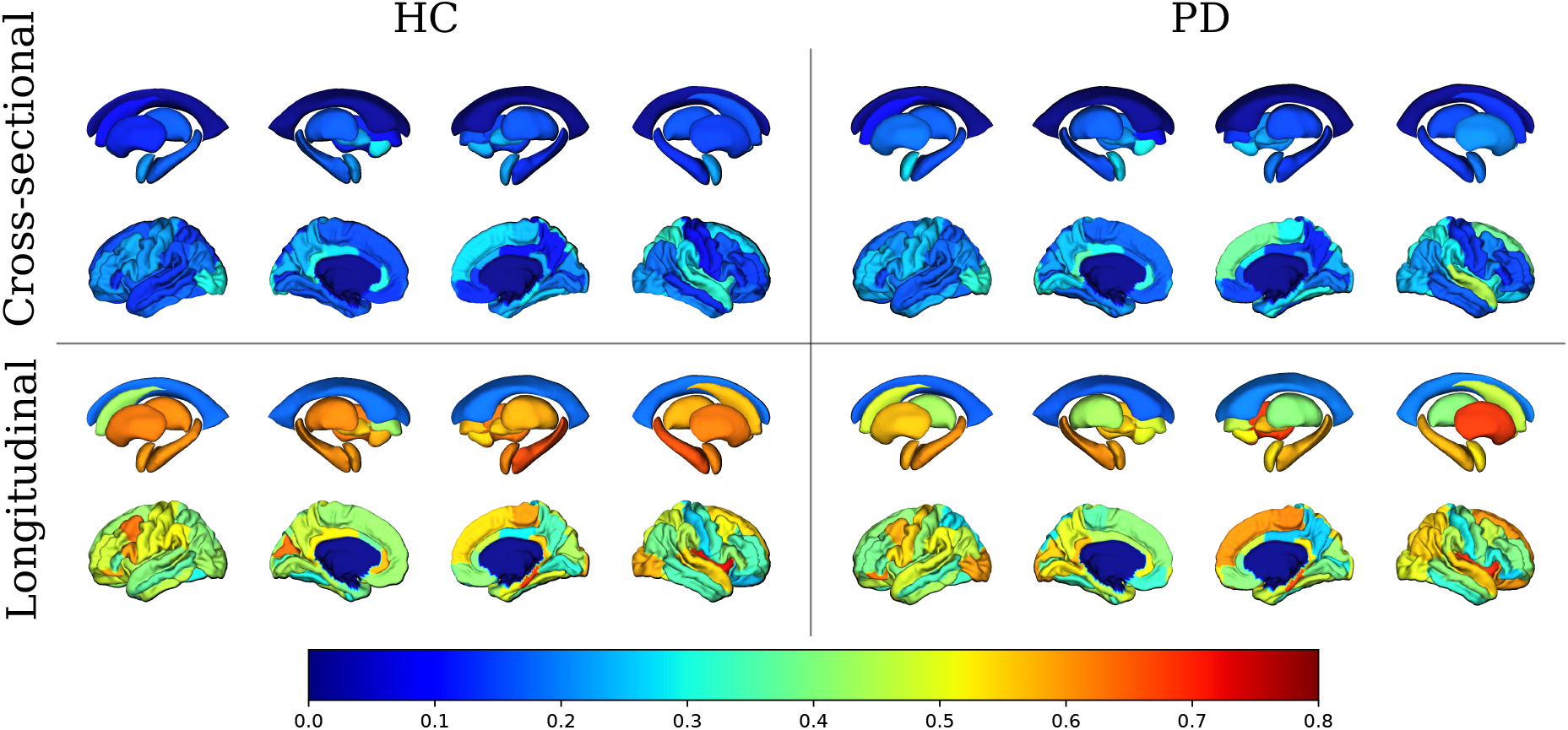
Numerical-Population Variability Ratio (*ν*_npv_) for subcortical volumes (top row in each panel) and cortical thickness (bottom row in each panel) in healthy controls (HC) and Parkinson’s disease (PD). Higher *ν*_npv_ values indicate higher numerical variability relative to inter-subject variability.

In contrast, longitudinal analyses exhibited substantially higher variability, with average *ν*_npv_ values of 0.561 for PD and 0.549 for HC; again, no difference survived Bonferroni correction across the four metrics (area *p* = 0.482, thickness *p* = 0.919, volume *p* = 0.370, subcortical volume *p* = 0.489; Supplementary Table S6). This amplification is likely driven by catastrophic cancellation that occurs when subtracting nearly identical values between time points. Consequently, achieving the same level of numerical variability (*σ*_*d*_ ≤ 0.01) would require a sample size exceeding 12,000 participants, highlighting the critical impact of numerical noise on reliability.

To facilitate the use of this framework, we developed an open-source, interactive web interface available at yohanchatelain.github.io/brain_render (Supplementary Figure S7). This tool allows researchers to upload standard summary statistics to instantly estimate the numerical variability floor of their findings. The interface includes a 3D visualization engine that maps numerical variability onto cortical and subcortical atlases, allowing for interactive exploration of pipeline stability. To aid interpretation, the tool automatically flags regions where the reported effect size is indistinguishable from the estimated numerical noise, thereby identifying potentially spurious findings.

### 2.3 Widespread impact of numerical variability on published findings

To assess the broader implications of numerical variability on MRI measures of Parkinson’s disease, we applied numerical variability propagation (Table 1) to re-evaluate findings from thirteen previously published articles reporting MRI measures of Parkinson’s disease, spanning a range of study designs and neuroscientific questions (selection criteria and study characteristics in Methods, §4.5 and Table 3), namely [18, 19, 20, 21, 22, 23, 24, 25, 26, 27, 28, 29, 30].

**Table 3:** Characteristics of the 13 retrospectively reviewed studies. *N* reports each study’s own analyzed sample sizes, as stated in that publication. Metrics lists the measures from which we extracted *p*-values for the numerical-variability re-analysis, which for some studies is a subset of all structural measures reported in the original publication. CT = cortical thickness; CV = cortical volume; CA = cortical area; SCV = subcortical volume.

| Study | Design | *N* | Metric(s) | Neuroscientific focus |
| --- | --- | --- | --- | --- |
| [18] | Cross-sectional | 43 PD | CT, CV, SCV | Psychiatric comorbidity (depression) |
| [19] | Cross-sectional | 36 HC, 92 PD | CT, CV | Cognitive correlates |
| [20] | Cross-sectional | 45 HC, 93 PD | CA, SCV | Cognitive heterogeneity/subtypes |
| [21] | Longitudinal | 18 HC, 32 PD | SCV | Mild cognitive impairment/decline |
| [22] | Both | 70 HC, 76 PD | CT, SCV | Disease-stage atrophy, clinical correlates |
| [23] | Cross-sectional | 56 HC, 90 PD | SCV | Motor subtypes |
| [24] | Cross-sectional | 90 PD | SCV, CT/CV | Cognitive impairment |
| [25] | Cross-sectional | 24 HC, 36 PD | CT, SCV | Psychiatric comorbidity (impulse control) |
| [26] | Cross-sectional | 28 HC, 28 PD | CT, CV, CA, SCV | Sleep disturbance |
| [27] | Both | 106 HC, 209 PD | SCV | Methodological (FreeSurfer version) |
| [28] | Longitudinal | 25 PD, 18 HC | SCV | Replication |
| [29] | Cross-sectional | 30 HC, 80 PD | CT, SCV | Disease staging/severity |
| [30] | Both | 34 PD, 32 HC | SCV | Longitudinal shape/volume progression |

For each *p*-value reported as significant in the original articles, we estimated the probability of a numerically induced significance flip using the Beta-distribution model described in Methods (§4.4), parametrized by the reported *p*-value (mean) and the propagated numerical variability (standard deviation). Numerical validation of this approach is provided in Supplementary Note S6.

Figure 4 shows the estimated probability of numerically induced significance flips as a function of the distance between reported *p*-values and the significance threshold (*p* − *α*), pooled across all statistical tests and the 13 reviewed papers (707 extracted results: 198 reported as significant and 509 as non-significant).

**Figure 4:**
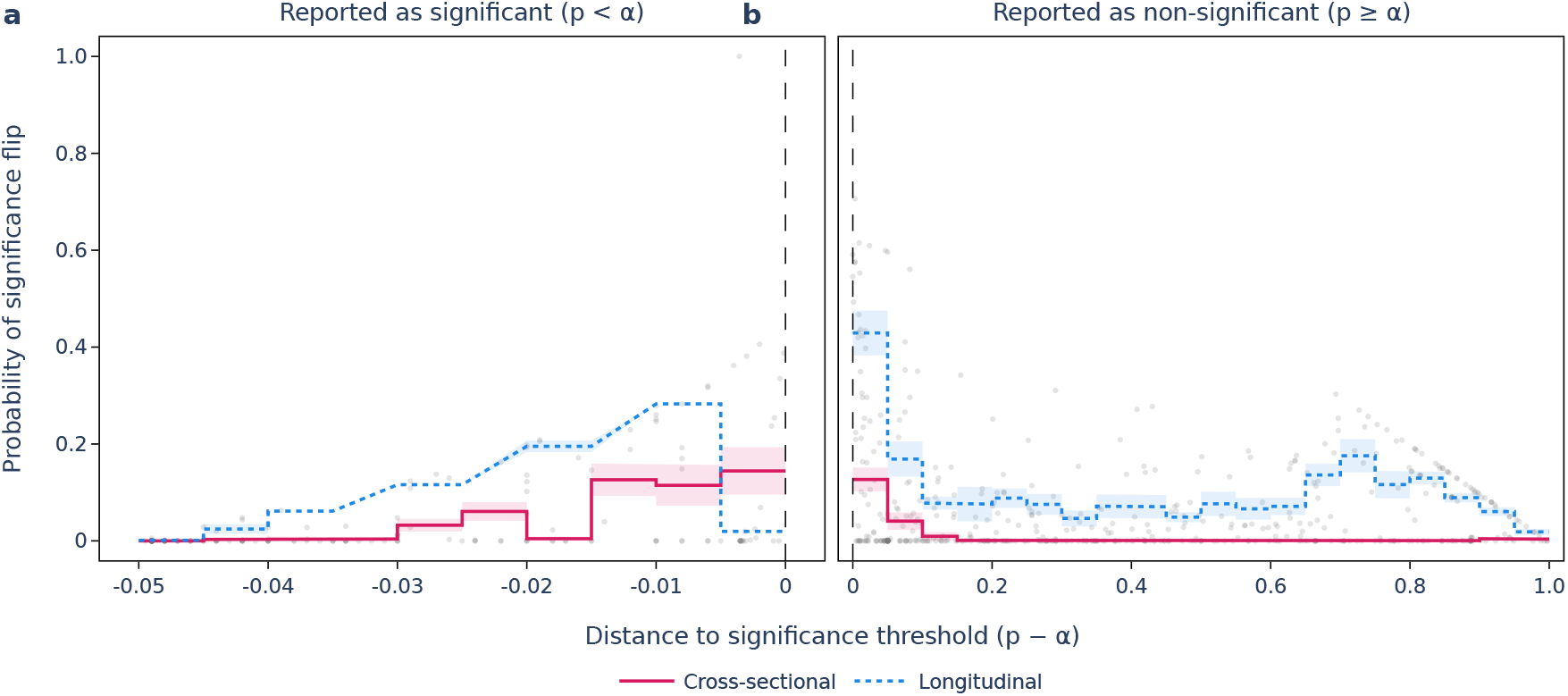
Probability of numerically-induced significance flips as a function of the *p*-value distance to significance threshold, pooled across statistics (T-values, F-values, and correlation coefficients; see Supplementary Note S6 for per-statistic breakdown). Each point represents a significant or non-significant result reported in one of the 13 papers that we reviewed. Negative *p* − *α* values correspond to results reported as significant (*p < α*), whereas positive *p* − *α* values correspond to results reported as non-significant (*p > α*). For results reported as significant, the y-axis gives the probability of flipping to non-significant under numerical variability (false positive risk); for results reported as non-significant, it gives the probability of flipping to significant (false negative risk). Lines indicate the mean flip probability within bins of size 0.005 for false positive risk and 0.05 for false negative risk. Standard error envelopes are shown in shaded areas when more than three points are available in a bin.

Significance flip probability peaks at the significance threshold (*p* − *α*=0), confirming that findings near this threshold are most likely to change significance status under numerical variability. By contrast, results farther from the threshold exhibit lower significance flip probabilities. High significance flip rates persist near the threshold across sample sizes, demonstrating that even well-powered studies can remain numerically fragile. This pattern is consistent with the theoretical relationship between numerical variability, sample size, and statistical variability (Table 1). Across all 707 extracted results, flip significance probability was only weakly associated with sample size (Pearson correlation *r* = − 0.118, *p* = 0.002). This overall association was driven by correlation-based tests, for which flip probability decreased significantly with sample size (*n* = 196, *r* = − 0.417, *p <* 10^−8^), consistent with the theoretical dependence of the sampling variance of the correlation coefficient on *n* (Table 1); for *T* - and *F* -based tests no significant association was found (*n* = 307, *r* = − 0.095, *p* = 0.096 and *n* = 204, *r* = − 0.093, *p* = 0.187, respectively). Significance flip probability therefore reflects the joint influence of distance to the significance threshold, sample size, and the Numerical-Population Variability Ratio. A breakdown by statistical test type (T-values, F-values, and correlation coefficients) is provided in Supplementary Note S6, where the pattern is consistent across test types: findings reported near the significance boundary carry substantially higher flip risk regardless of the statistic used. Longitudinal analyses exhibit a higher risk of significance flips than cross-sectional analyses (0.05 probability of significance flip for cross-sectional analyses on average, compared to 0.1 for longitudinal analyses). Moreover, longitudinal results reported as non-significant carry a high risk of significance flip even for p-values far from the significance threshold consistently with the data presented in Figure 3, suggesting a high risk of false negatives in longitudinal pipelines. These results reveal a widespread impact of numerical variability on reported MRI measures of Parkinson’s disease, affecting findings across multiple test types and study designs, and underscore the need for systematic numerical robustness assessments in neuroimaging studies.

## 3 Discussion

In this study, we demonstrated that numerical variability arising from floating-point computation substantially affects structural MRI analyses of Parkinson’s disease, accounting for 18% of population variability in cross-sectional analyses and 55% in longitudinal analyses on average, with up to 80% in some regions. By systematically perturbing the FreeSurfer pipeline using Monte Carlo Arithmetic, we showed that numerical noise alone accounted for a non-negligible fraction of population variability in both cortical and subcortical measures. This variability was sufficient to alter downstream statistical inference, leading to frequent significance flips in group comparisons and clinical correlations, even when pipeline configurations and computational environments were held constant. These findings highlight how numerical noise contributes to reproducibility challenges reported in neuroimaging studies.

The Numerical-Population Variability Ratio (NPVR) introduced in this work formalizes the relationship between numerical variability and population variability. By analytically propagating NPVR through common statistical estimators, we established explicit relationships between numerical variability, sample size, and the variability in effect sizes and p-values. This framework shows that numerical variability does not vanish with increasing sample size in the same manner as sampling error, and that results near conventional significance thresholds are sensitive to numerical perturbations. Crucially, while applied here to numerical variability, this formalism is generalizable: the NPVR could model the propagation of other independent error sources, such as test-retest variability or instrumentation noise, offering a unified perspective on non-sampling errors.

Importantly, NPVR enables practical numerical quality evaluations without requiring access to raw imaging data or computationally expensive reprocessing. Applying this framework to previously published Parkinson’s disease MRI studies revealed that a fraction of reported significant findings are susceptible to numerically-induced significance flips, highlighting that numerical variability is not only a technical artifact but a substantive factor that can directly influence scientific conclusions. These thirteen studies spanned a range of designs (eight cross-sectional, two longitudinal, and three combining both) and neuroscientific questions, including cognitive correlates and impairment, psychiatric and behavioral comorbidities, motor subtypes, disease staging and progression, and sleep disturbance (Methods, §4.5; Table 3), indicating that numerically induced fragility is not confined to a narrow category of neuroimaging findings but is broadly distributed across the PD structural-MRI literature. By making this assessment feasible from summary statistics alone, NPVR provides a scalable approach for retrospective evaluation of the existing literature. In this context, NPVR can be used by researchers and reviewers to estimate the potential contribution of numerical noise to observed false positive rates. Numerical validation (see Supplementary Note S5) demonstrated that these estimates tend to minimize the risk of underestimating the impact of numerical variability.

Although our empirical analyses focused on FreeSurfer 7.3.1, the observed variability is not unique to this tool. Prior work [14] has documented comparable numerical sensitivity in other widely used neuroimaging pipelines, including FMRIB Software Library [31] (FSL) and Advanced Normalization Tools [32] (ANTs), while suggesting lower sensitivity in Statistical Parametric Mapping [33] (SPM), potentially due to differences in optimization strategies and regularization. These findings indicate that numerical variability is a property of computational pipelines rather than a specific implementation artifact. The magnitude of the variability then depends on algorithmic design choices, numerical precision, and optimization dynamics. Torabi et al. [34] compared methodological variability to biological variability in dynamic functional connectivity assessment methods, reporting a ratio of methodological to biological variability of 0.94 across seven different methods. This ratio is comparable to the worst case NPVR values observed in our study.

The increasing adoption of deep learning-based components in neuroimaging pipelines does not eliminate numerical variability but instead shifts its focus. The latest FreeSurfer release (v8) now integrates deep learning models such as SynthSeg [35], SynthStrip [36], and SynthMorph [37] to replace classical segmentation, skull-stripping and registration steps. While inference in trained models is relatively stable [38], training introduces additional sources of numerical variability comparable to non-deep learning methods [15]. Quantifying and controlling numerical variability across both classical and deep learning approaches remains therefore critical.

We considered several factors that may impact the generalizability of our findings. The Parkinson’s disease cohort analyzed here is relatively homogeneous in age and phenotype, which could reduce population variance and inflate NPVR estimates. However, we observed no significant differences in numerical variability between patients and healthy controls, supporting the interpretation that numerical variability is primarily a property of the computational pipeline rather than the clinical population. Extending NPVR measurements across diverse datasets, disease contexts, and software packages will be important to build a comprehensive map of computational reliability in neuroimaging, though our results already suggest that numerical variability is sufficiently large to warrant routine consideration.

Our results also highlight that the impact of numerical variability differs across MRI metrics. Cortical area and cortical volume showed the highest rates of significance flipping, reaching up to 53% of regions in longitudinal analyses, while cortical thickness and subcortical volume were comparatively more stable (Supplementary Table S5). These metric-specific differences should inform the interpretation of neuroimaging findings and motivate targeted investigation of numerical stability in analyses relying on the most sensitive metrics.

Based on our findings, we offer the following practical recommendations for neuroimaging researchers. First, numerical stability should be systematically evaluated using the NPVR framework or the associated web tool (yohanchatelain.github.io/brain_render), particularly for longitudinal analyses where numerical amplification is pronounced. Second, pipeline development for longitudinal applications should explicitly prioritize numerical robustness. Third, these numerically focused practices should complement broader recommendations for robust neuroimaging analyses: statistical inference should move beyond binary *p*-value thresholds by reporting effect sizes, confidence intervals, and both significant and non-significant results [39]; results should be validated across multiple neuroimaging pipelines because numerical sensitivity varies across software [1, 14]; and multiverse analyses should be used to characterize how analytical decisions affect conclusions [40]. Fourth, a public gallery comparing numerical variability and population variability across neuroimaging tools and pipelines, modeled on existing galleries of effect sizes such as BrainEffeX [41], would help researchers situate the numerical variability of a given tool relative to the population variability typically observed for the same measure, and compare this relationship across tools.

More broadly, this work highlights that numerical variability should be treated as a serious component of uncertainty in neuroimaging, alongside biological variability and statistical sampling error. Floating-point rounding and truncation are only one contributor; algorithmic choices, preprocessing decisions, and data handling practices all interact with numerical precision to shape final results. Extending this quantification to the deep-learning training stage is equally important, given the field’s central role in modern neuroimaging, and would support more robust and interpretable models. Systematic quantification of these effects is essential for the development of numerically robust software and for the reliable translation of neuroimaging biomarkers into clinical and personalized-medicine settings.

In conclusion, NPVR provides a principled, interpretable, and scalable framework to expose hidden numerical variability in neuroimaging analyses. By enabling routine evaluation of numerical variability, this approach strengthens transparency, supports more reliable inference, and offers a concrete path toward improving reproducibility in computational neuroscience.

## 4 Methods

We quantified the impact of numerical variability on structural MRI findings in Parkinson’s disease (PD) using (1) stochastic perturbation experiments and (2) analytical variability propagation. Empirically, we processed a longitudinal PPMI cohort using FreeSurfer 7.3.1 instrumented with stochastic numerical noise to isolate run-to-run variability. Analytically, we derived closed-form approximations linking this pipeline variability (*ν*_npv_) to the variability in effect sizes and test statistics, enabling retrospective quality control using only summary statistics.

The experimental workflow proceeded in four stages: (1) curation of a longitudinal dataset with two visits per participant; (2) repeated processing under Monte Carlo Arithmetic (MCA) perturbations via Fuzzy-libm; (3) rigorous quality control to distinguish numerical artifacts from technical failures; and (4) statistical evaluation of inference variability across repetitions.

### 4.1 Participants

Structural MRI data were obtained from the Parkinson’s Progression Markers Initiative (PPMI; www.ppmi-info.org). The study included 201 participants: 112 individuals diagnosed with Parkinson’s disease without mild cognitive impairment (PD-non-MCI) and 89 healthy controls (HC). All participants had two usable T1-weighted MRI scans acquired approximately 1.4 ± 0.5 years apart (0.9–2.0 years). Patients with mild cognitive impairment were excluded to minimize confounding effects of cognitive decline.

Inclusion criteria were: (i) diagnosis of idiopathic Parkinson’s disease (PD-non-MCI) or healthy control status; (ii) availability of two high-quality T1-weighted scans at distinct visits; and (iii) absence of other neurological or psychiatric conditions. PD severity was quantified using the Unified Parkinson’s Disease Rating Scale part III (UPDRS-III) in the OFF medication state at both baseline and follow-up visits.

All procedures were approved by the research ethics boards of participating PPMI sites, and written informed consent was obtained from all participants in accordance with the Declaration of Helsinki and was exempt from review by Concordia University’s Research Ethics Unit. The PD and HC groups did not differ significantly in age (*t* = − 0.035, *p* = 0.972), education (*t* = − 1.479, *p* = 0.141; two-sample *t*-tests), or sex distribution (*χ*^2^ = 3.108, *p* = 0.078; chi-square test; Table 2).

### 4.2 Image acquisition and preprocessing

High-resolution T1-weighted MRI scans were obtained from the Parkinson’s Progression Markers Initiative (PPMI). Data were acquired using standardized 3D MPRAGE protocols (TR = 2.3 s, TE = 2.98 ms, TI = 0.9 s, voxel size = 1 mm isotropic) across multiple sites. While the protocol was standardized, we accounted for minor acquisition variations inherent to the PPMI multisite design.

Structural images were processed using FreeSurfer 7.3.1 instrumented with Fuzzy-libm, a modified mathematical library that injects stochastic perturbations into floating-point operations to probe numerical stability (Section 4.3). To estimate numerical variability, each scan was processed 34 times. We filtered the resulting outputs to exclude both technical failures (e.g., incomplete execution and FreeSurfer failures) and quality control (QC) failures.

For QC, we visually inspected nine representative slices (3 along each axis) per run to identify gross artifacts, such as missing brain tissue, blurring, or biologically implausible segmentation. Supplementary Table S4 details the exclusion rates. To ensure statistical consistency across the cohort, we randomly subsampled the remaining runs to retain exactly 26 valid realisations per participant.

### 4.3 Numerical variability assessment

To quantify the numerical variability in the FreeSurfer pipeline, we employed Monte Carlo Arithmetic (MCA) [17]. MCA is a stochastic technique that simulates the propagation of rounding errors by introducing controlled random perturbations into floating-point operations. We utilized the Random Rounding (RR) mode, where the result of an arithmetic operation is perturbed by a zero-mean random noise scaled to the magnitude of the least significant bit. For any operation producing an exact result *x*, the perturbed result 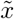 is modeled as 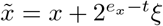 where *e*_*x*_ = ⌊log_2_ |*x*|⌋ is the exponent of *x, t* is the *virtual precision* parameter, and *ξ* is a random variable drawn uniformly from the interval 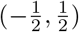.

While comprehensive MCA instrumentation provides a rigorous bound on numerical error, it incurs a prohibitive computational cost (typically 100 × to 1000 × slowdown), rendering it intractable for largescale neuroimaging cohorts. To address this, we used *Fuzzy-libm* [9], a lightweight implementation that restricts MCA instrumentation to elementary mathematical library functions (e.g., `exp`, `log`, `sin`, `cos`). By targeting these elementary functions, which are sources of divergence across operating systems [7], Fuzzy-libm significantly reduces computational overhead while effectively capturing the numerical variability relevant to cross-platform reproducibility [9, 12, 11]. This machine-level noise is representative of the numerical differences introduced by typical variations in hardware (e.g., CPU models), software libraries (e.g., operating-system updates), and parallelization (e.g., sequential vs. multi-threaded execution), and its validity as a model of realistic numerical variability in neuroimaging pipelines has been demonstrated for operating-system [9] and hardware [12] variations.

Fuzzy-libm is deployed via a Docker container and uses the `LD_PRELOAD` mechanism to dynamically intercept calls to the standard system math library and redirect them to the instrumented version. The library is compiled using Verificarlo [42], an LLVM-based compiler that injects the MCA logic at compile time. Virtual precision parameters are set to match standard hardware precision (*t* = 53 bits for double precision and *t* = 24 bits for single precision), ensuring that the simulated variability remains representative of realistic machine-level precision errors.

#### 4.3.1 Numerical-Population Variability Ratio (*ν*_npv_)

To quantify computational stability relative to population variation, we introduce the Numerical-Population Variability Ratio (*ν*_npv_). For each brain region, *ν*_npv_ measures the ratio of numerical variability arising from computational processes to natural inter-subject variation:

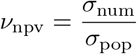

where *σ*_num_ represents numerical variability (measurement precision across MCA repetitions for individual subjects) and *σ*_pop_ represents population variability (inter-subject differences within each repetition). For each region of interest, measurements from *k* MCA repetitions across *n* subject-visit pairs form a data matrix ℳ_*k×n*_ with entries 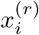, where *i* = 1, …, *n* indexes subject-visits and *r* = 1, …, *k* indexes repetitions. Let

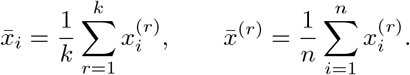

##### Numerical variability (within-subject, across repetitions)

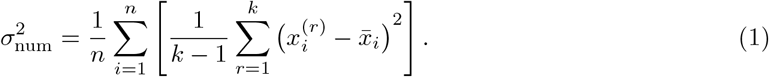

##### Population variability (within-repetition, across subjects)

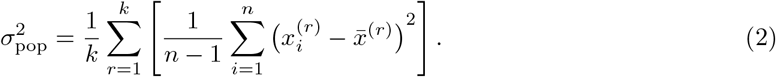

where 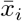 and 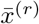 denote column and row means, respectively. Higher *ν*_npv_ values indicate regions where numerical variability approaches population variation.

#### 4.3.2 Relationship between *ν*_npv_ and downstream statistical test variability

To establish a quantitative link between a method’s computational reproducibility and the reliability of group-level statistical inferences, we derived analytical expressions connecting numerical variability to the resulting variability of commonly used statistical tests (Cohen’s *d, t*-tests, partial correlation, and ANCOVA). Our goal is to characterize how numerical noise propagates through the analytical pipeline to produce variability in the reported statistics.

For each MCA repetition *r*, we denote by 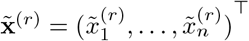 the vector of perturbed measurements across *n* subjects. Each perturbed observation is modeled as the sum of the subject’s biological value and a numerical error term:

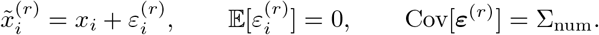

Here, **x** = (*x*_1_, …, *x*_*n*_)^⊤^ represents the fixed, biological measurements, while ***ε***^(*r*)^ captures the random numerical perturbations introduced during computation.

##### Assumptions

To isolate the contribution of numerical variability, we make three simplifying assumptions:

1. **Numerical error model**. The numerical perturbations are modeled as independent, zero-mean Gaussian random variables: 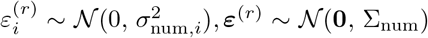, where, under homoscedasticity, 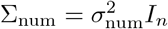.
2. **Population variability**. The between-subject variance 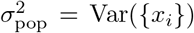 dominates the numerical noise, i.e. *σ*_num,*i*_ ≪ *σ*_pop_. This implies that the pooled empirical standard deviation of the observed data can be approximated by the biological one, *s*_*p*_ ≈ *σ*_pop_.
3. **Null-hypothesis scenario**. We condition on the observed biological measurements **x** and quantify variability only from numerical perturbations ***ε***^(*r*)^. This isolates numerical variability effects from sampling variation; the resulting expressions remain accurate near the null and for small effect sizes. In this setting, the biological values **x** = (*x*_1_, …, *x*_*n*_)^⊤^ are treated as fixed, and all randomness arises from numerical perturbations ***ε***^(*r*)^.

Under these assumptions, each downstream statistic 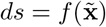 can be linearized around the baseline **x** as 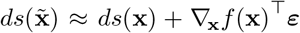, allowing the numerical variance to be expressed through the delta method as

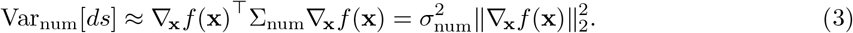

Table 1 summarizes the derived expressions for the numerical standard deviation of several common statistics. Proofs of partial derivations of sample statistics^1^ used in the following sections are provided in the Supplementary Note S1.

**Cohen’s** d Cohen’s effect size quantifies the standardized difference between two sample means. For two independent groups *G*_1_ and *G*_2_ with sample sizes *n*_1_ and *n*_2_ (*df* = *n*_1_ + *n*_2_ − 2), we define:

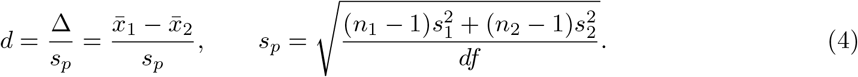

The variance of *d* across Monte Carlo Arithmetic (MCA) repetitions, conditional on the fixed dataset **x**, is given by:

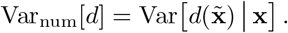

Applying the multivariate delta method (Eq. 3) around the baseline **x**_0_ = **x**, we obtain:

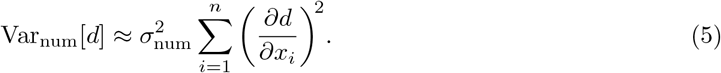

For an observation *x*_*i*_ ∈ *G*_*g*_ (*g* ∈ {1, 2}), the chain rule gives:

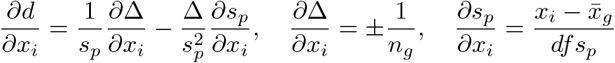

where the sign in *∂*Δ*/∂x*_*i*_ is positive for *g* = 1 and negative for *g* = 2. Substituting back into the expression for *∂d/∂x*_*i*_, we have:

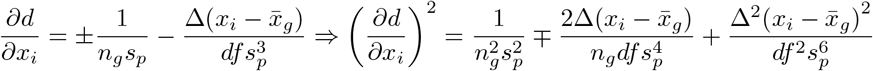

and summing over all *i* in group *G*_*g*_:

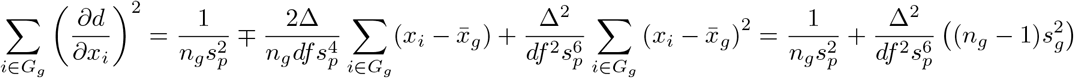

since 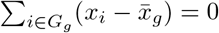, so finally summing over both groups:

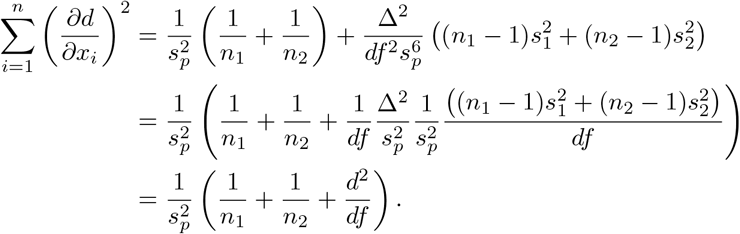

Finally, assuming *s*_*p*_ ≈ *σ*_pop_, the population (between-subject) variance, Eq. (5) becomes:

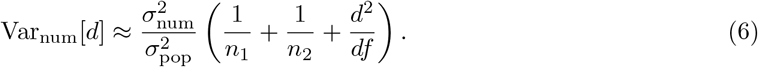

For balanced groups (*n*_1_ = *n*_2_ = *n/*2) and large *n*, the *d*^2^*/df* term is negligible:

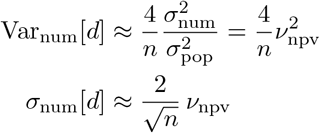

where *ν*_npv_ = *σ*_num_*/σ*_pop_ is the numerical-population variability ratio. This expression quantitatively links the numerical variability captured by *ν*_npv_ to the dispersion of Cohen’s *d* effect size, providing a practical measure of the stability of statistical inferences under finite-precision arithmetic.

##### Two-sample t-test statistic

The pooled two-sample *t* statistic quantifies the standardized difference between two group means:

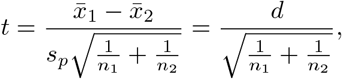

where *d* is Cohen’s *d* defined in Eq. (4). Defining 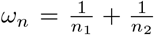, the *t* statistic can be expressed as 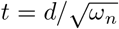. From Eq. (6), the variance of *d* due to numerical perturbations propagates to the variance of *t* as:

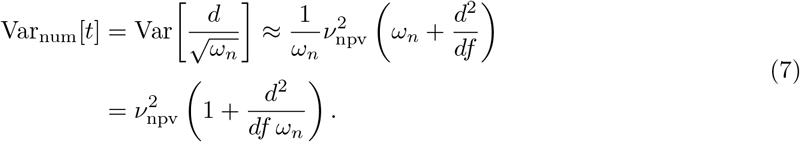

To analyze the correction term, consider

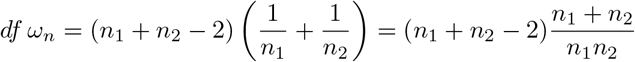

When 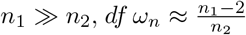 symmetrically, when 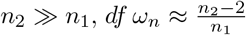. In both cases, 1*/*(*df ω*_*n*_) → 0, so the correction term 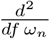 vanishes for unbalanced groups. When the groups are balanced (*n*_1_ = *n*_2_ = *n/*2), 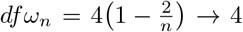 as *n* → ∞, so that 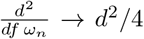. Substituting these results into Eq. (7) gives:

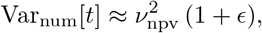

where *ϵ* tends to 0 for strongly unbalanced groups and to *d*^2^*/*4 for large balanced samples. For small effect sizes (*d*^2^ ≪ 4) and unbalanced groups, the correction term *ϵ* is negligible, yielding the simplified expression:

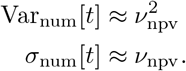

The variability of the corresponding *p*-values can then be derived. Let *X* be the random variable with E_num_[*X*] = *t*_0_ and 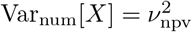 . Let *f*_*t*_, *F*_*t*_ be the probability density and cumulative distribution functions of the Student *t*-distribution with *df* degrees of freedom. Applying the delta method to the two-sided *p*-value *p*(*X*) = 2 (1 − *F*_*t*_(|*X*|)) gives:

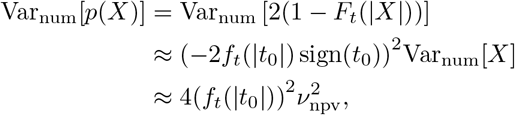

which gives the standard deviation of the numerical variability in the *p*-value:

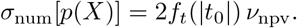

##### ANCOVA group effect

Analysis of covariance (ANCOVA) evaluates group differences using a general linear model:

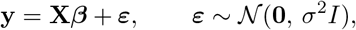

where **y** is the vector of measurements across subjects, and **X** includes an intercept, diagnostic group (PD vs. HC), and covariates (e.g., age and sex). The adjusted group difference is expressed as the one-degree-of-freedom contrast *c*^⊤^***β***, with contrast vector *c* = [0, 1, 0, 0]^⊤^. Let 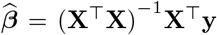 denote the ordinary least-squares (OLS) estimator and 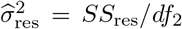 the residual mean square, where *df*_2_ = *n* − rank(**X**). The sum of squares associated with the group effect is:

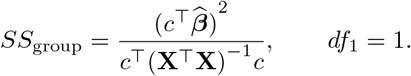

The corresponding ANCOVA *F* statistic is given by:

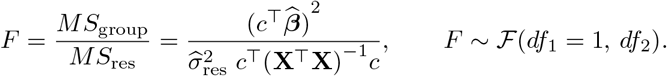

Significance is evaluated using the upper-tail of the central ℱ distribution under the null hypothesis:

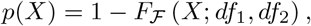

with *X* ∼ ℱ(*df*_1_, *df*_2_) and *F*_ℱ_ the cumulative distribution function of the *F* distribution with (*df*_1_, *df*_2_) degrees of freedom. For *df*_1_ = 1, the ANCOVA *F* statistic is equivalent to the two-sample *t*-test through *F* = *t*^2^ (see [43, p. 403]). Then the variability in the *F* statistic due to numerical noise follows directly from the variability of *t*:

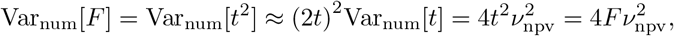

yielding

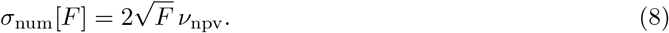

The variability in the corresponding *p*-values can be obtained by the delta method. Let *X* be a random variable with E_num_[*X*] = *F*_0_ and 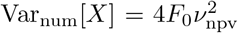 with *f*_ℱ_, *F*_ℱ_ the probability density and cumulative distribution functions. Applying the delta-method to the upper-tail *p*-value is *p*(*X*) = 1 − *F*_ℱ_ (*X*) yields:

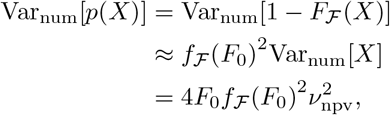

so that

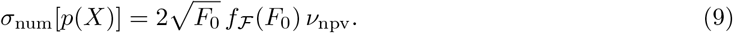

Equations (8) and (9) show that numerical imprecision introduces a variance in the estimated *F* statistic and its corresponding *p*-value that scales linearly with the numerical-population variability ratio *ν*_npv_, and proportionally to 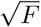 for the group effect magnitude.

##### Partial correlation

Partial correlation measures the association between two variables (*x, y*) while controlling for the influence of one or more additional variables *z*. In our analysis, this corresponds to quantifying the relationship between regional brain measurements and UPDRS-III motor scores, controlling for age and sex. The sample partial correlation is defined as:

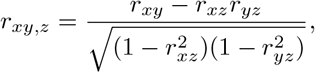

where *r*_*xy*_ denotes the Pearson correlation between variables *x* and *y*,

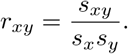

To simplify notation, we set *a* = *r*_*xy*_, *b* = *r*_*xz*_, and *c* = *r*_*yz*_ so that

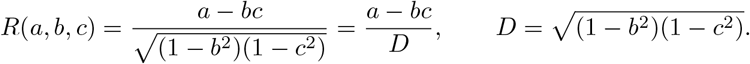

Applying the delta method (Eq. 3) to the partial correlation, we have:

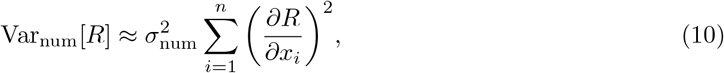

Assuming only *x* is affected by numerical perturbations while *y* and *z* are fixed, the gradient with respect to each observation *x*_*i*_ is:

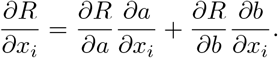

The first-order partial derivatives are:

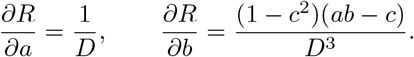

and the derivatives of the correlations with respect to *x*_*i*_ are (see Supplementary Eq. S6)

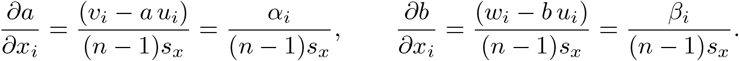

where 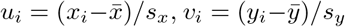, and 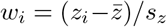 are standardized and centered observations of *x, y*, and *z* respectively. Then *∂R/∂x*_*i*_ becomes:

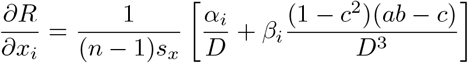

thus (*∂R/∂x*_*i*_)^2^ is:

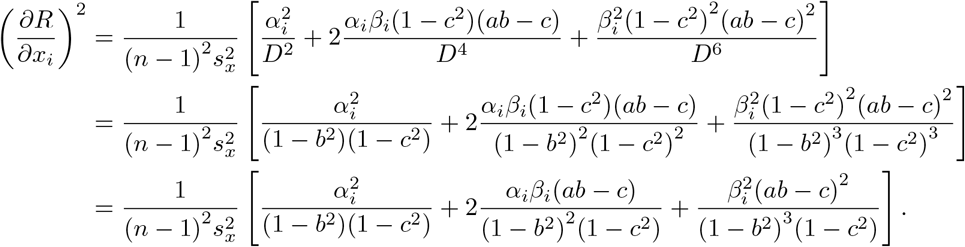

Using the correlation identities 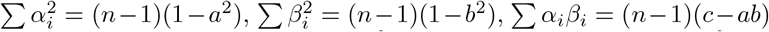 (see Supplementary Eq. S8), and that (1 − *b*^2^)(1 *c*^2^) (*a bc*)^2^ = (1 − *a*^2^)(1 *b*^2^) (*c ab*)^2^, we sum over all *i* to obtain:

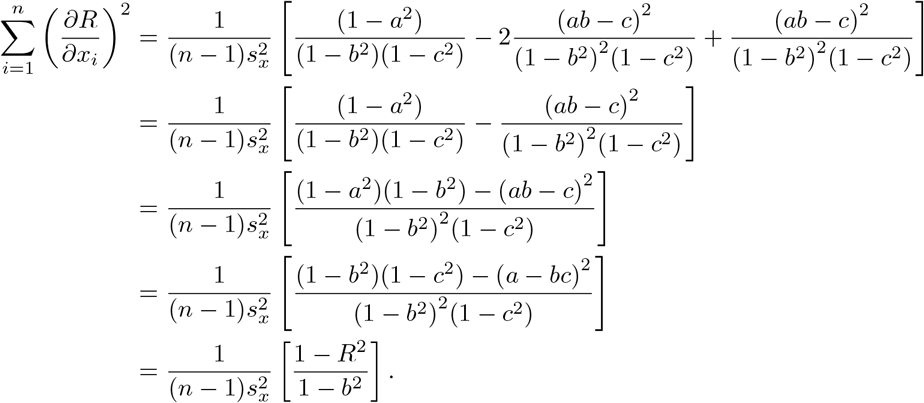

Substituting back into Eq. (10) with 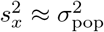 gives:

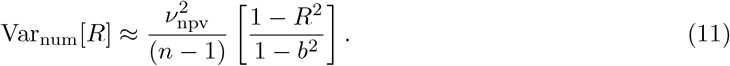

Since *b* is rarely reported in practice, we further simplify this expression by deriving a lower bound on Var_num_[*R*]. Since 0 ≤ *b*^2^ ≤ 1, then 1*/*(1 − *b*^2^) ≥ 1. Therefore:

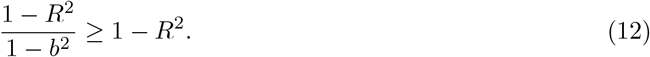

So, substituting into Eq. (11) gives the lower bound:

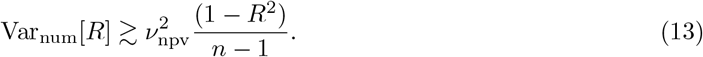

Taking the square root yields the standard deviation:

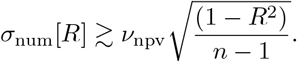

The two-sided significance of a partial correlation is computed from the *t*-statistic

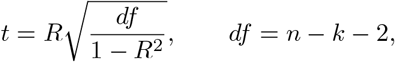

where *k* is the number of controlling variables. Let *X* be the random variable with E_num_[*X*] = *R*_0_, 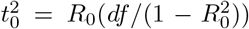 with Var_num_[*X*] bounded by Equation (13). Let *f*_*t*_, *F*_*t*_ be the probability density and cumulative distribution functions of the Student *t*-distribution with *df* degrees of freedom. Applying the delta method to the two-sided *p*-value *p*(*X*) = 2 (1 − *F*_*t*_(|*X*|)) gives:

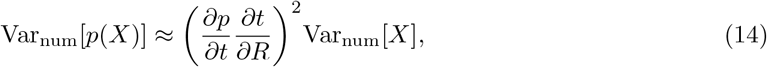

with the partial derivatives given by:

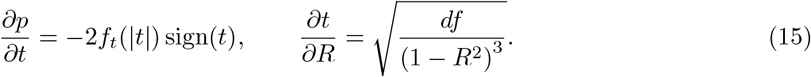

Combining equations (14) and (15) yields

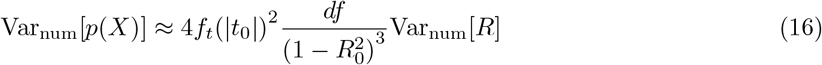

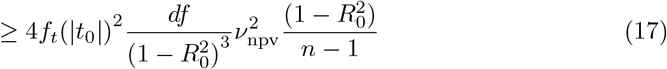

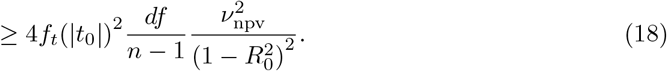

Using Equation (13) to bound Var_num_[*R*] leads to:

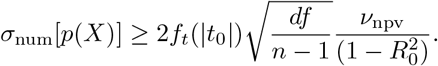

### 4.4 Probability of significance flips induced by numerical variability

To quantify the probability of significance flips findings arising from numerical variability, we model the computed *p*-value as a random variable following a Beta distribution, *p* ∼ Beta(*a, b*), which is suitable for modeling probabilities and proportions bounded on [0, 1]. The distribution is parameterized by shape parameters (*a, b*) determined from a target mean *µ*_*p*_ and variance 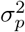 estimated using the variability propagation formulae reported in Table 1. The parameters are given by:

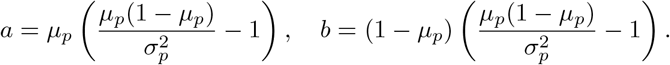

These shape parameters are positive, i.e. the Beta distribution is well defined only when 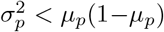.

This formulation defines a distribution of plausible *p*-values centered around the nominal value *p*_0_ = *µ*_*p*_, with dispersion determined by numerical variability. Importantly, this model enables direct estimation of error rates caused by numerical perturbations relative to a fixed significance threshold *α*. Two distinct regimes are considered, depending on whether the nominal *p*-value *p*_0_ lies above or below *α* (Fig. 5):

**Figure 5:**
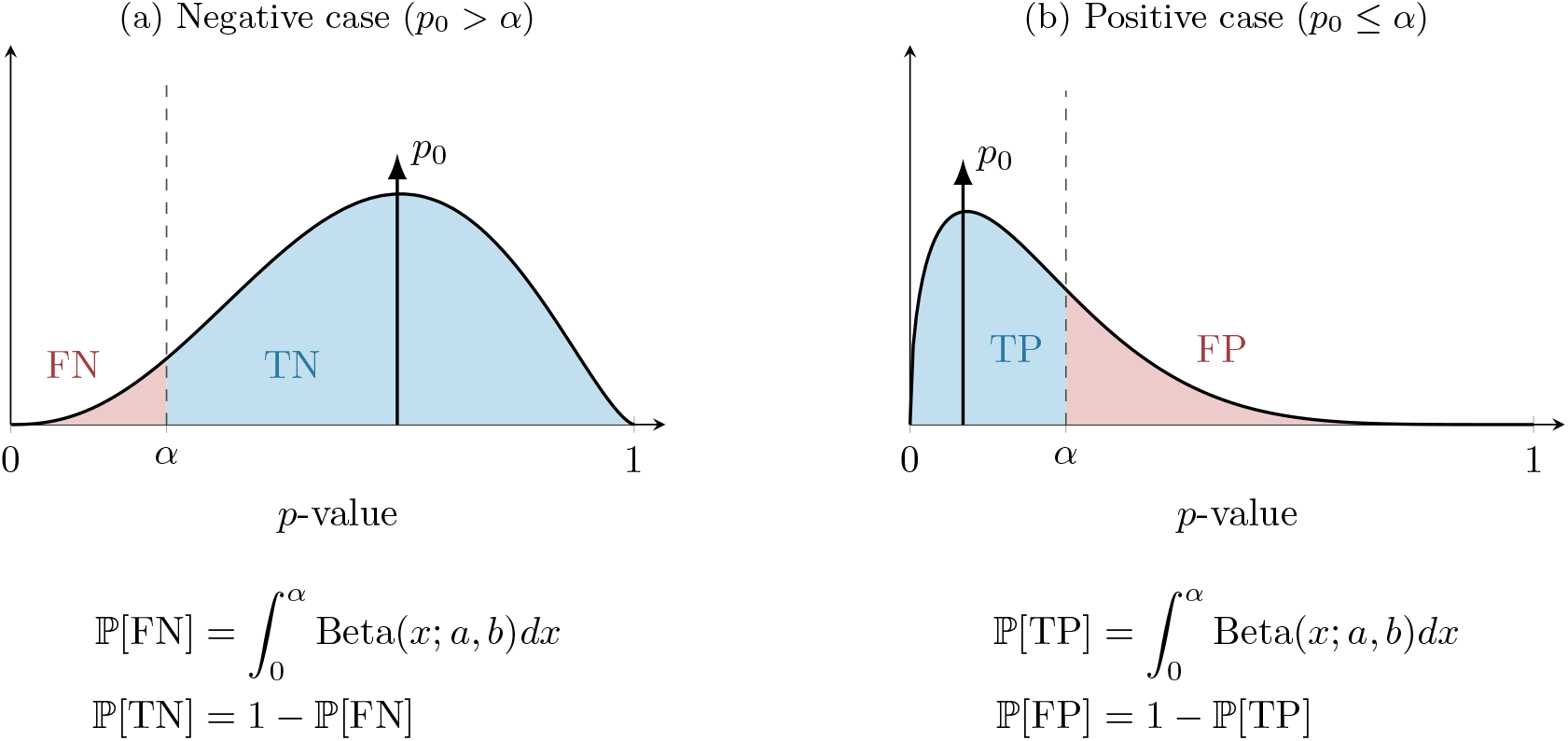
Modeling inference stability using Beta distributions. The panels illustrate the probability of significance flipping due to numerical variability. The negative case (*p*_0_ *> α*, Fig. 5a), where the tail of the distribution crossing *α* represents the probability of a false negative finding (FN). The positive case (*p*_0_ ≤ *α*, Fig. 5b), where the tail extending beyond *α* represents the probability of a false positive finding (FP).

- Negative Case (*p*_0_ *> α*): The primary finding is non-significant, i.e., a true negative (TN). Here, we calculate the probability that numerical noise shifts the *p*-value below *α*, resulting in a false negative (FN).
- Positive Case (*p*_0_ ≤ *α*): The primary finding is significant, i.e., a true positive (TP). We calculate the probability that numerical noise shifts the *p*-value above *α*, resulting in a false positive (FP).

The probabilities for these outcomes are obtained by computing the cumulative distribution function (CDF) of the Beta distribution up to the threshold *α*, as illustrated in Figure 5. This probabilistic modeling provides an explicit link between numerical variability and classical Type I and Type II error rates, allowing us to isolate the contribution of numerical variability to spurious statistical significance. Numerical validation of this model is presented in Supplementary Note S6.

### 4.5 Overview of evaluated studies

We compiled a set of previously published studies reporting FreeSurfer-derived structural MRI findings in Parkinson’s disease, to illustrate the impact of numerical variability on reported outcomes across a range of study designs and research questions. Eligible studies had to report, for at least one cortical or subcortical measure, a *p*-value together with the sample size and statistical test used, so that numerical variability could be estimated from the formulas in Table 1 without requiring access to the original raw data. Thirteen studies met these criteria and were retained [18, 19, 20, 21, 22, 23, 24, 25, 26, 27, 28, 29, 30].

Table 3 summarizes the design, sample size, FreeSurfer-derived metric(s), and neuroscientific focus of each study. The set spans eight purely cross-sectional studies, two purely longitudinal studies, and three studies contributing both cross-sectional and longitudinal analyses, addressing questions of cognitive correlates and impairment (4 studies), psychiatric and behavioral comorbidities (2), motor subtypes (1), disease staging, atrophy, and progression (3), sleep disturbance (1), and methodological replication (2).

For each study, we extracted every *p*-value reported for a FreeSurfer-derived structural measure, together with the corresponding test statistic, sample size, and test type, and propagated numerical variability through the significance-flip model described in Section 4.4.

## 5 Data Availability

The data that support the findings of this study are available from the Parkinson’s Progression Markers Initiative (PPMI) database (www.ppmi-info.org/access-data-specimens/download-data), but restrictions apply to the availability of these data, which were used under license for the current study, and so are not publicly available. Data are available from the authors upon reasonable request and with permission of the PPMI.

## 6 Code Availability

All MCA instrumentation scripts, FreeSurfer build instructions, and analysis notebooks are available at github.com/yohanchatelain/livingpark-numerical-variability. The exact commit hashes are archived on Zenodo (10.5281/zenodo.21522109) to ensure reproducibility.

## 7 Acknowledgements

The analyses were conducted on the Virtual Imaging Platform [44], which utilizes resources provided by the biomed Virtual Organization within the European Grid Infrastructure (EGI). We extend our gratitude to Sorina Pop from CREATIS, Lyon, France, for her support. This work was funded by the Michael J. Fox Foundation for Parkinson’s Research (MJFF-021134; www.michaeljfox.org). The funders had no role in study design, data collection and analysis, decision to publish, or preparation of the manuscript.

## Declarations

### Author Contributions

Y.C. and T.G. conceived and designed the study. Y.C. implemented the MCA instrumentation, conducted the analyses, and developed the web tool. A.S. contributed to data curation and analysis. M.S. contributed to clinical interpretation of the results. J.-B.P. contributed to the conceptualization of the study and the statistical analyses. T.G., M.S., and J.-B.P. acquired funding and supervised the project. Y.C. drafted the manuscript. All authors contributed to reviewing and editing the manuscript and approved the final version.

### Competing Interests

The authors declare no competing interests.

---

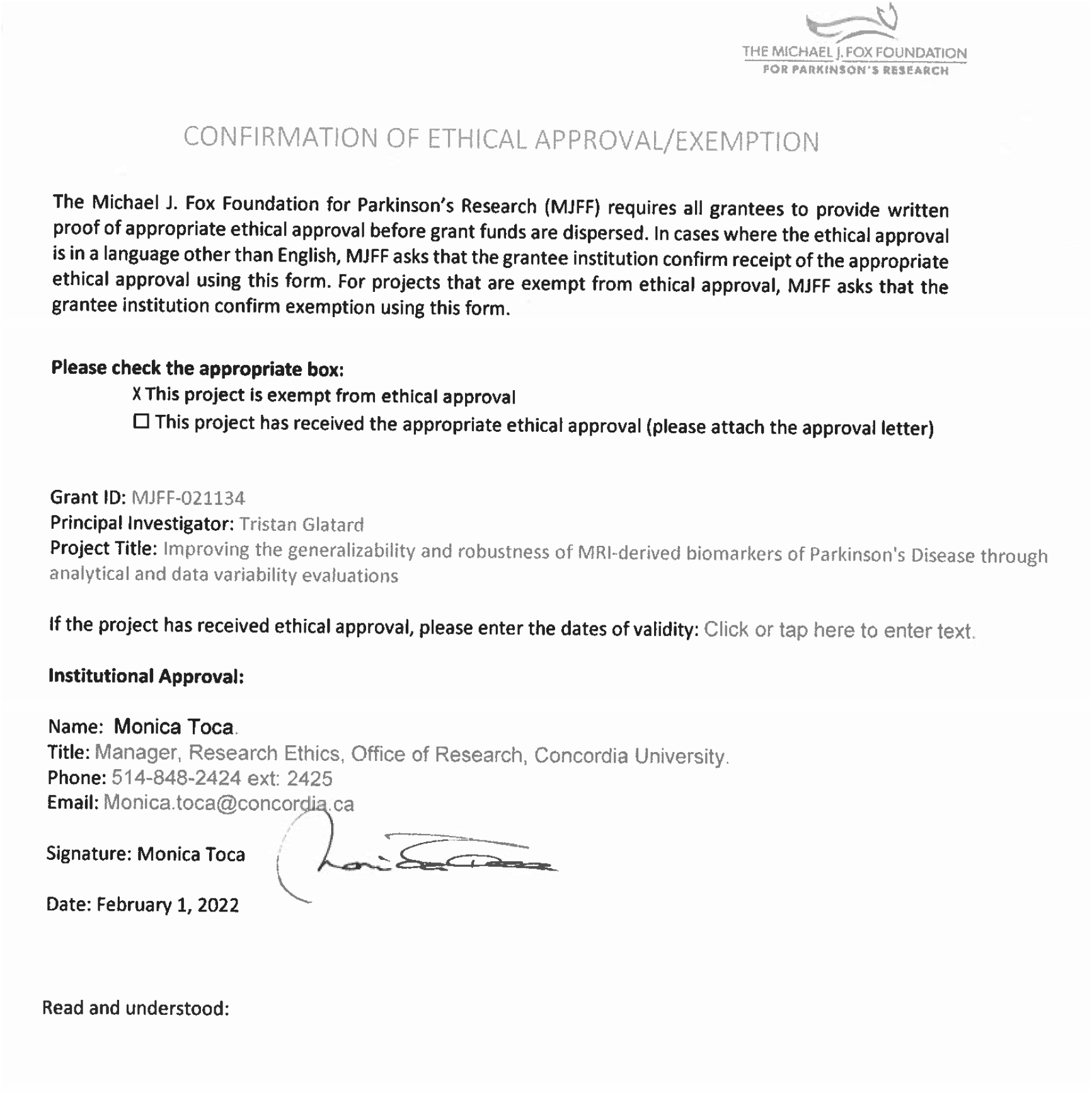

## Supplementary Information

This Supplementary Information provides the derivations, additional analyses, and validation experiments supporting the main text. It is organised as follows:

- **Section S1** derives the partial derivatives of common sample statistics that underlie the closed-form variability-propagation formulas of Table 1 (Methods, “Relationship between NPVR and downstream statistical test variability”).
- **Section S2** quantifies cross-sectional numerical variability of FreeSurfer measures (significant digits and spatial-overlap Dice coefficients), with region-by-region precision tables supporting the low-precision claim in Results, Section 2.1.
- **Section S3** reports the region-level significance-flip frequencies, the HC-vs-PD permutation test, and the per-region Ansari-Bradley comparisons showing that longitudinal processing amplifies variance (Results, Sections 2.1–2.3).
- **Section S4** shows the distributions of test-statistic coefficients across MCA repetitions, including the unperturbed (IEEE-754) reference used to check the validity of the perturbation approach (Results, Section 2.1).
- **Section S5** numerically validates the variability-propagation formulas of Table 1 against sampled MCA estimates.
- **Section S6** numerically validates the Beta-distribution significance-flip model used in the retrospective analysis (Results, Section 2.3).

### S1 Partial derivatives of sample statistics

These derivations underpin the delta-method variability-propagation formulas of Table 1 (Methods, “Relationship between NPVR and downstream statistical test variability”). We derive below the partial derivatives of common sample statistics for a dataset *x* = {*x*_1_, *x*_2_, …, *x*_*n*_} with respect to an individual observation *x*_*i*_, where *n* denotes the sample size. The Kronecker delta *δ*_*ij*_ equals 1 when *i* = *j* and 0 otherwise.

#### Sample Mean

The partial derivative of the sample mean with respect to *x*_*i*_ is constant:

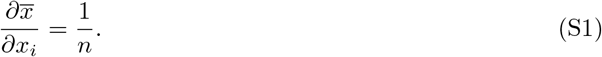

*Proof*. The sample mean is defined as

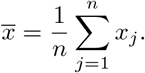

Taking the partial derivative with respect to *x*_*i*_ gives

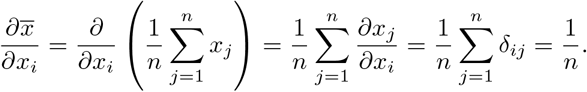

#### Sample Variance

The partial derivative of the sample variance with respect to *x*_*i*_ is

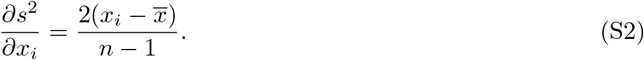

*Proof*. The sample variance is defined as

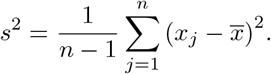

Differentiating with respect to *x*_*i*_ yields

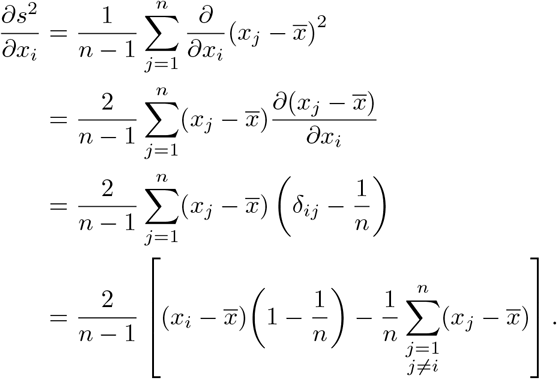

Since 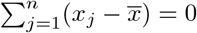 then 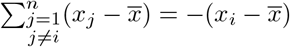 so the second term simplifies, giving

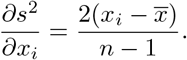

#### Sample Standard Deviation

The partial derivative of the sample standard deviation with respect to *x*_*i*_ is

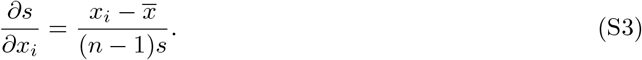

*Proof*. Given that 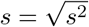, the derivative follows directly from the chain rule:

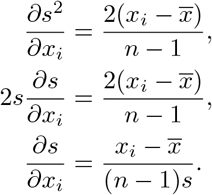

#### Pooled Standard Deviation

The pooled standard deviation is a weighted average of the variances of two groups |*G*_1_| = *n*_1_ and |*G*_2_| = *n*_2_ with *df* = *n*_1_ + *n*_2_ − 2. Its partial derivative with respect to *x*_*i*_, *i* ∈ *G*_*g*_ is given by

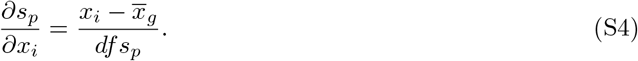

*Proof*. Let *x* be partitioned into two groups *G*_1_ and *G*_2_ with sizes *n*_1_ and *n*_2_, 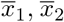 be the sample means and *s*_1_, *s*_2_ be the sample standard deviations of groups *G*_1_ and *G*_2_, respectively. Let *df* = *n*_1_ + *n*_2_ − 2. Then the pooled standard deviation *s*_*p*_ is defined as

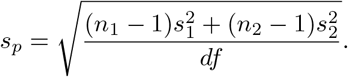

Differentiating *s*_*p*_ with respect to *x*_*i*_ in group *G*_*g*_ gives:

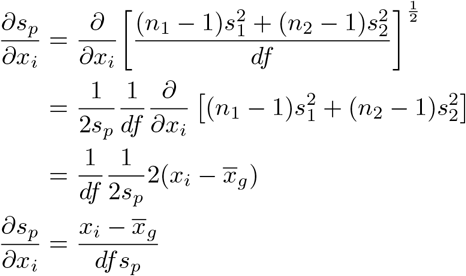

#### Sample Covariance

The partial derivative of the sample covariance with respect to *x*_*i*_ is

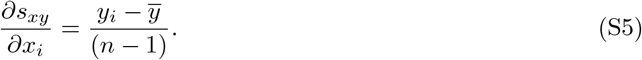

*Proof*. The sample covariance between two variables *x* and *y* is defined as

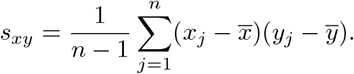

Taking the partial derivative with respect to *x*_*i*_ gives

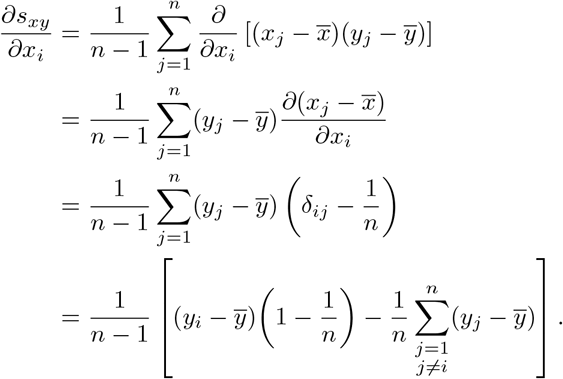

Since 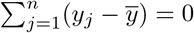 then 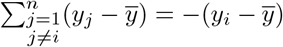 so the second term simplifies, giving

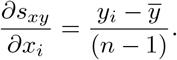

#### Pearson correlation coefficient

The partial derivative of *r*_*x,y*_ with respect to an individual observation *x*_*i*_ is given by

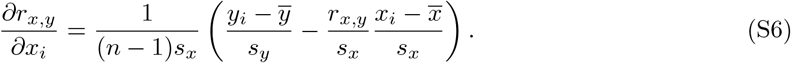

*Proof*. The Pearson correlation coefficient *r* between two variables *x* and *y* is defined as

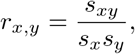

using the quotient rule, we differentiate *r*(*x, y*) with respect to *x*_*i*_:

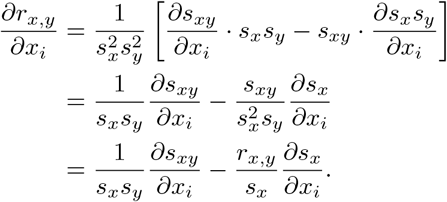

Substituting the partial derivatives of the sample covariance (Eq. S5) and standard deviation (Eq. S3) we obtain

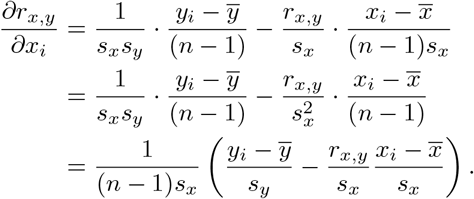

#### Correlation identities

Let 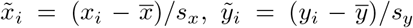 and 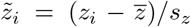 be the standardized variables. The following identities hold:

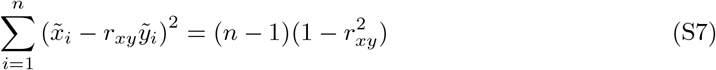

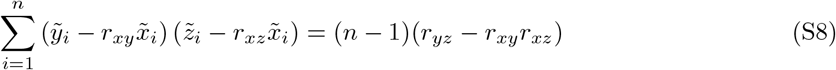

*Proof*. Note that 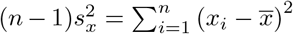 and 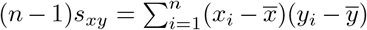 then by using the definitions of standardized variables, we have for the first identity:

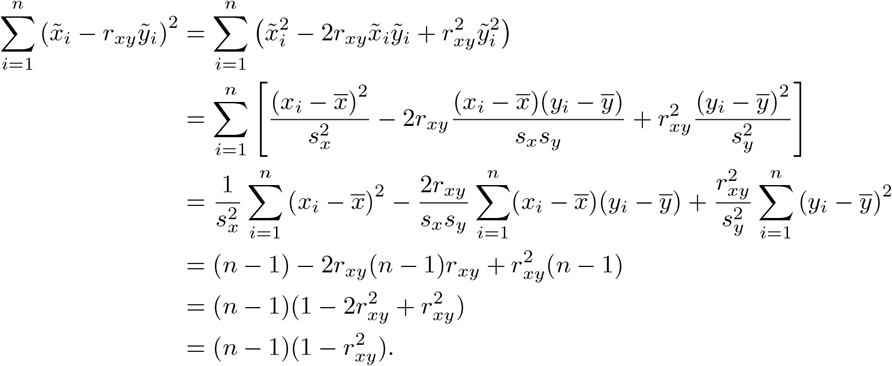

and for the second identity:

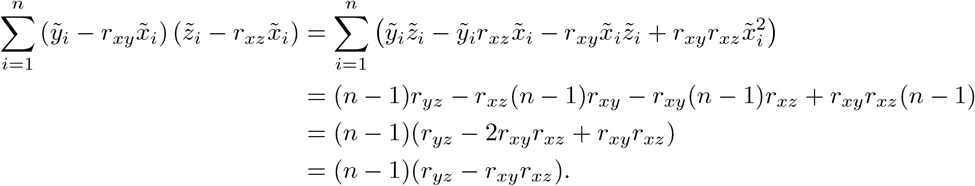

### S2 Cross-sectional numerical variability quantification

As a complementary component of our analysis, we quantified cross-sectional numerical variability for cortical and subcortical measures. Numerical variability was assessed using two metrics: (1) the number of significant digits [45] (*p*_*s*_ = 0.95, confidence 1 − *α*_*s*_ = 0.95), calculated using the `significantdigits` package^2^ (version 0.4.0); and (2) the extended Sørensen-Dice coefficient, which measures the spatial overlap of segmentation masks across *n* repetitions. The extended Dice coefficient is defined as:

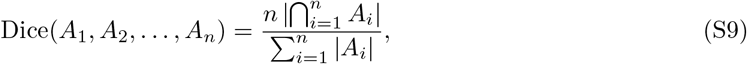

where *A*_*i*_ represents the segmentation for the *i*-th Monte Carlo observation.

FreeSurfer 7.3.1 demonstrated limited numerical precision across all anatomical measures. Cortical metrics showed averages of 1.62 ± 0.20 significant digits for thickness, 1.34 ± 0.24 for surface area, and 1.34 ± 0.23 for cortical volume (Figure S2). Subcortical volumes exhibited similar variability, with an average of 1.33 ± 0.22 significant digits (Figure S3). These results indicate that, on average, FreeSurfer measurements are precise to only one decimal place, with specific instances exhibiting complete loss of significant digits. While variability was observed across all metrics, cortical thickness displayed the highest relative precision (range: 1.22 − 1.94 digits), whereas surface area (0.83 − 1.73) and cortical volume (0.80 − 1.73) were less stable. Table S1 and Table S2 provide the region-by-region breakdown underlying these averages: within-subject significant digits and standard deviation for each cortical metric and hemisphere (lh/rh = left/right hemisphere); lower significant-digit values indicate regions where numerical variability more strongly erodes precision. Table S3 presents the corresponding region-level data for subcortical volumes.

To assess spatial stability, we evaluated the volumetric overlap using the extended Sørensen-Dice coefficient (Eq. S9). The analysis revealed substantial inter-subject and regional variability (Figure S1). Cortical regions showed marked variability, with Dice coefficients ranging from 0.00 to 0.91 (mean 0.73 ± 0.13), indicating that for some regions, numerical noise could lead to a complete lack of spatial overlap. Subcortical structures appeared comparatively more stable, ranging from 0.03 to 0.94 (mean 0.81 ± 0.09). Overall, subcortical segmentations demonstrated higher spatial robustness compared to cortical parcellations.

**Table S1:**
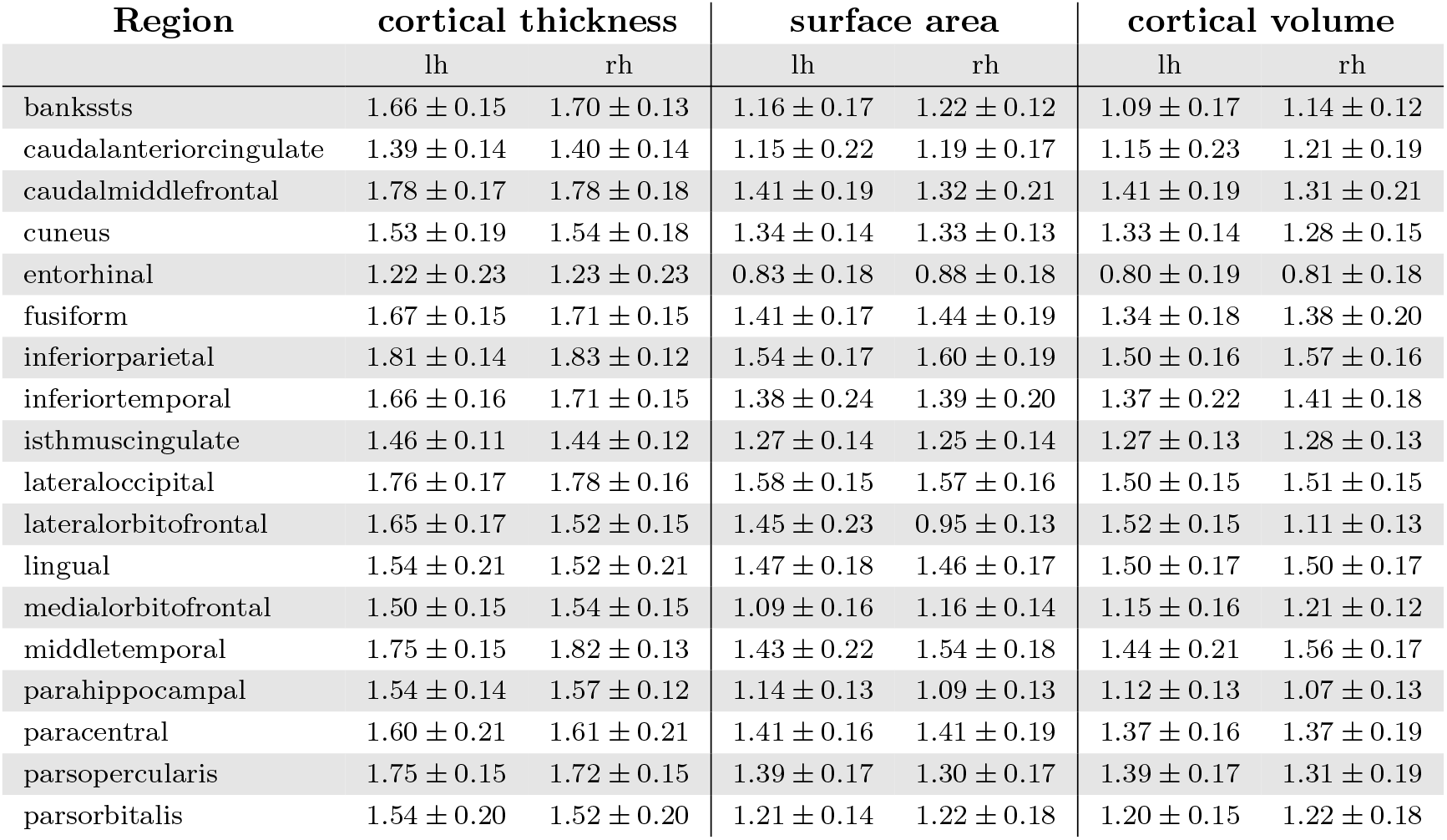
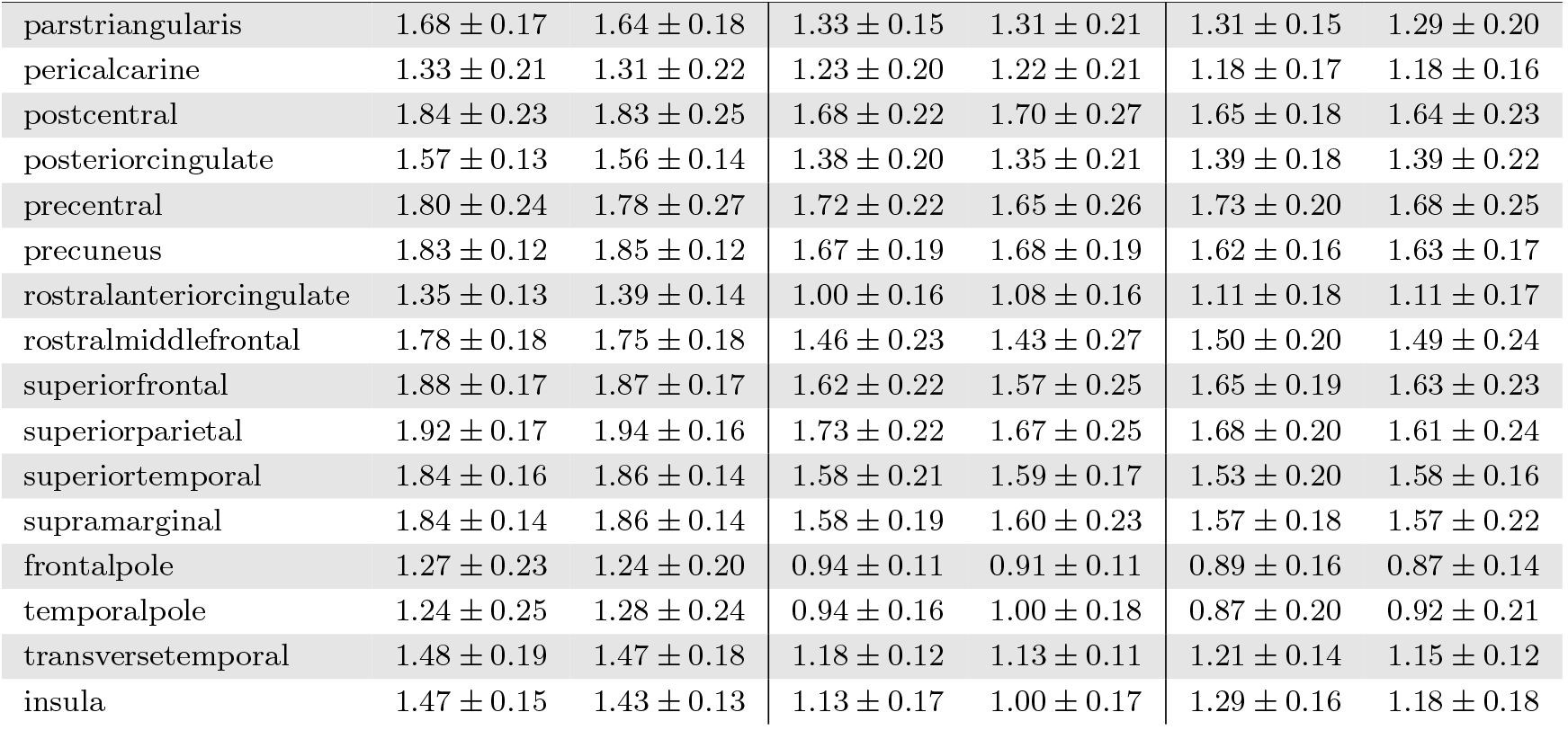
Within-subject significant digits averaged across all subjects.

| Region | cortical thickness |  | surface area |  | cortical volume |  |
| --- | --- | --- | --- | --- | --- | --- |
|  | lh | rh | lh | rh | lh | rh |
| bankssts | 1.66 ± 0.15 | 1.70 ± 0.13 | 1.16 ± 0.17 | 1.22 ± 0.12 | 1.09 ± 0.17 | 1.14 ± 0.12 |
| caudalanteriorcingulate | 1.39 ± 0.14 | 1.40 ± 0.14 | 1.15 ± 0.22 | 1.19 ± 0.17 | 1.15 ± 0.23 | 1.21 ± 0.19 |
| caudalmiddlefrontal | 1.78 ± 0.17 | 1.78 ± 0.18 | 1.41 ± 0.19 | 1.32 ± 0.21 | 1.41 ± 0.19 | 1.31 ± 0.21 |
| cuneus | 1.53 ± 0.19 | 1.54 ± 0.18 | 1.34 ± 0.14 | 1.33 ± 0.13 | 1.33 ± 0.14 | 1.28 ± 0.15 |
| entorhinal | 1.22 ± 0.23 | 1.23 ± 0.23 | 0.83 ± 0.18 | 0.88 ± 0.18 | 0.80 ± 0.19 | 0.81 ± 0.18 |
| fusiform | 1.67 ± 0.15 | 1.71 ± 0.15 | 1.41 ± 0.17 | 1.44 ± 0.19 | 1.34 ± 0.18 | 1.38 ± 0.20 |
| inferiorparietal | 1.81 ± 0.14 | 1.83 ± 0.12 | 1.54 ± 0.17 | 1.60 ± 0.19 | 1.50 ± 0.16 | 1.57 ± 0.16 |
| inferiortemporal | 1.66 ± 0.16 | 1.71 ± 0.15 | 1.38 ± 0.24 | 1.39 ± 0.20 | 1.37 ± 0.22 | 1.41 ± 0.18 |
| isthmuscingulate | 1.46 ± 0.11 | 1.44 ± 0.12 | 1.27 ± 0.14 | 1.25 ± 0.14 | 1.27 ± 0.13 | 1.28 ± 0.13 |
| lateraloccipital | 1.76 ± 0.17 | 1.78 ± 0.16 | 1.58 ± 0.15 | 1.57 ± 0.16 | 1.50 ± 0.15 | 1.51 ± 0.15 |
| lateralorbitofrontal | 1.65 ± 0.17 | 1.52 ± 0.15 | 1.45 ± 0.23 | 0.95 ± 0.13 | 1.52 ± 0.15 | 1.11 ± 0.13 |
| lingual | 1.54 ± 0.21 | 1.52 ± 0.21 | 1.47 ± 0.18 | 1.46 ± 0.17 | 1.50 ± 0.17 | 1.50 ± 0.17 |
| medialorbitofrontal | 1.50 ± 0.15 | 1.54 ± 0.15 | 1.09 ± 0.16 | 1.16 ± 0.14 | 1.15 ± 0.16 | 1.21 ± 0.12 |
| middletemporal | 1.75 ± 0.15 | 1.82 ± 0.13 | 1.43 ± 0.22 | 1.54 ± 0.18 | 1.44 ± 0.21 | 1.56 ± 0.17 |
| parahippocampal | 1.54 ± 0.14 | 1.57 ± 0.12 | 1.14 ± 0.13 | 1.09 ± 0.13 | 1.12 ± 0.13 | 1.07 ± 0.13 |
| paracentral | 1.60 ± 0.21 | 1.61 ± 0.21 | 1.41 ± 0.16 | 1.41 ± 0.19 | 1.37 ± 0.16 | 1.37 ± 0.19 |
| parsopercularis | 1.75 ± 0.15 | 1.72 ± 0.15 | 1.39 ± 0.17 | 1.30 ± 0.17 | 1.39 ± 0.17 | 1.31 ± 0.19 |
| parsorbitalis | 1.54 ± 0.20 | 1.52 ± 0.20 | 1.21 ± 0.14 | 1.22 ± 0.18 | 1.20 ± 0.15 | 1.22 ± 0.18 |
|  | lh | rh | lh | rh | lh | rh |
| parstriangularis | 1.68 ± 0.17 | 1.64 ± 0.18 | 1.33 ± 0.15 | 1.31 ± 0.21 | 1.31 ± 0.15 | 1.29 ± 0.20 |
| pericalcarine | 1.33 ± 0.21 | 1.31 ± 0.22 | 1.23 ± 0.20 | 1.22 ± 0.21 | 1.18 ± 0.17 | 1.18 ± 0.16 |
| postcentral | 1.84 ± 0.23 | 1.83 ± 0.25 | 1.68 ± 0.22 | 1.70 ± 0.27 | 1.65 ± 0.18 | 1.64 ± 0.23 |
| posteriorcingulate | 1.57 ± 0.13 | 1.56 ± 0.14 | 1.38 ± 0.20 | 1.35 ± 0.21 | 1.39 ± 0.18 | 1.39 ± 0.22 |
| precentral | 1.80 ± 0.24 | 1.78 ± 0.27 | 1.72 ± 0.22 | 1.65 ± 0.26 | 1.73 ± 0.20 | 1.68 ± 0.25 |
| precuneus | 1.83 ± 0.12 | 1.85 ± 0.12 | 1.67 ± 0.19 | 1.68 ± 0.19 | 1.62 ± 0.16 | 1.63 ± 0.17 |
| rostralanteriorcingulate | 1.35 ± 0.13 | 1.39 ± 0.14 | 1.00 ± 0.16 | 1.08 ± 0.16 | 1.11 ± 0.18 | 1.11 ± 0.17 |
| rostralmiddlefrontal | 1.78 ± 0.18 | 1.75 ± 0.18 | 1.46 ± 0.23 | 1.43 ± 0.27 | 1.50 ± 0.20 | 1.49 ± 0.24 |
| superiorfrontal | 1.88 ± 0.17 | 1.87 ± 0.17 | 1.62 ± 0.22 | 1.57 ± 0.25 | 1.65 ± 0.19 | 1.63 ± 0.23 |
| superiorparietal | 1.92 ± 0.17 | 1.94 ± 0.16 | 1.73 ± 0.22 | 1.67 ± 0.25 | 1.68 ± 0.20 | 1.61 ± 0.24 |
| superiortemporal | 1.84 ± 0.16 | 1.86 ± 0.14 | 1.58 ± 0.21 | 1.59 ± 0.17 | 1.53 ± 0.20 | 1.58 ± 0.16 |
| supramarginal | 1.84 ± 0.14 | 1.86 ± 0.14 | 1.58 ± 0.19 | 1.60 ± 0.23 | 1.57 ± 0.18 | 1.57 ± 0.22 |
| frontalpole | 1.27 ± 0.23 | 1.24 ± 0.20 | 0.94 ± 0.11 | 0.91 ± 0.11 | 0.89 ± 0.16 | 0.87 ± 0.14 |
| temporalpole | 1.24 ± 0.25 | 1.28 ± 0.24 | 0.94 ± 0.16 | 1.00 ± 0.18 | 0.87 ± 0.20 | 0.92 ± 0.21 |
| transversetemporal | 1.48 ± 0.19 | 1.47 ± 0.18 | 1.18 ± 0.12 | 1.13 ± 0.11 | 1.21 ± 0.14 | 1.15 ± 0.12 |
| insula | 1.47 ± 0.15 | 1.43 ± 0.13 | 1.13 ± 0.17 | 1.00 ± 0.17 | 1.29 ± 0.16 | 1.18 ± 0.18 |

**Figure S1:**
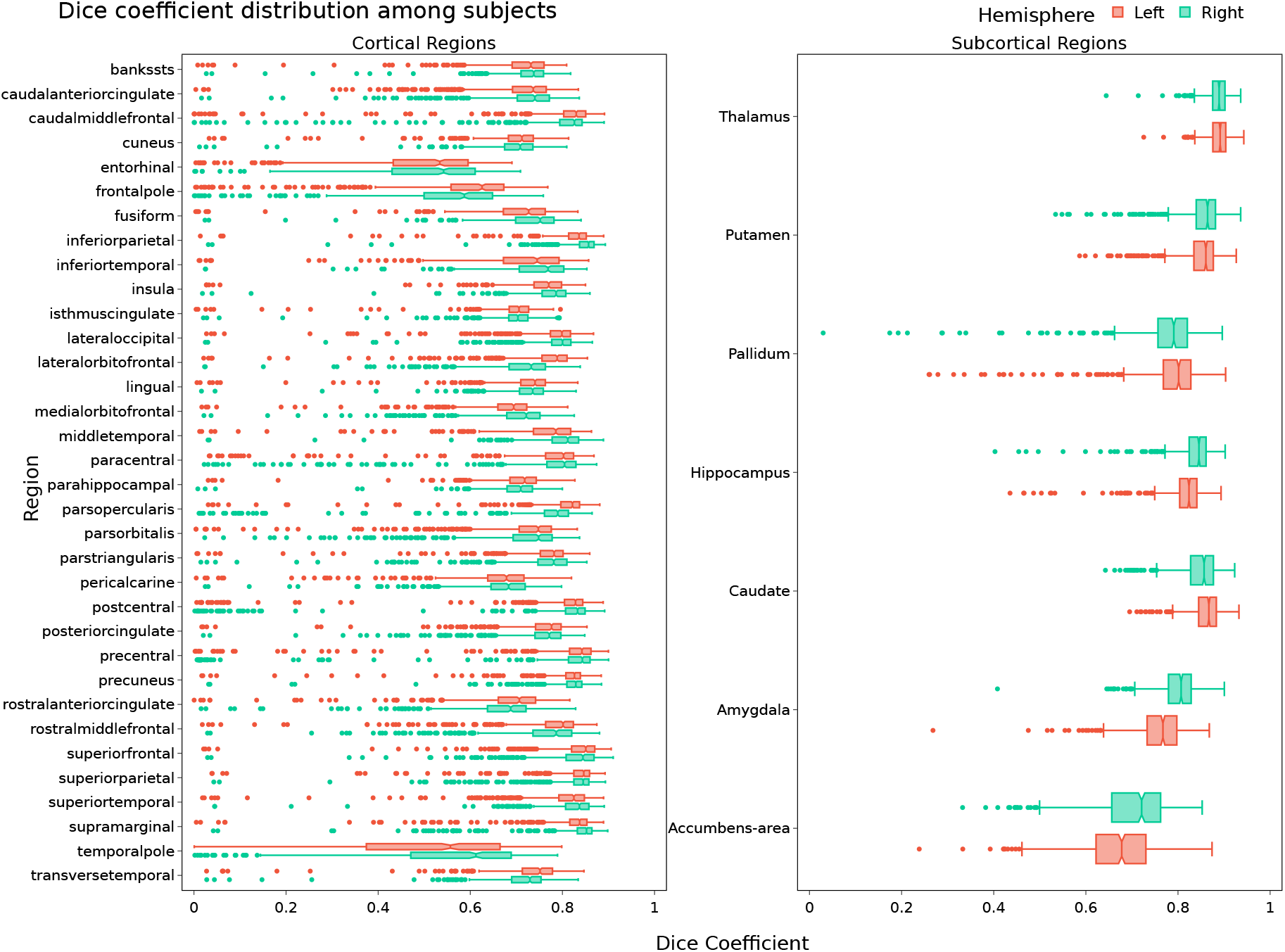
Numerical variability of segmentation. The distribution of extended Sørensen-Dice coefficients, highlighting the variability in spatial overlap for cortical and subcortical segmentations due to numerical variability. Lower values indicate higher variability.

**Table S2:**
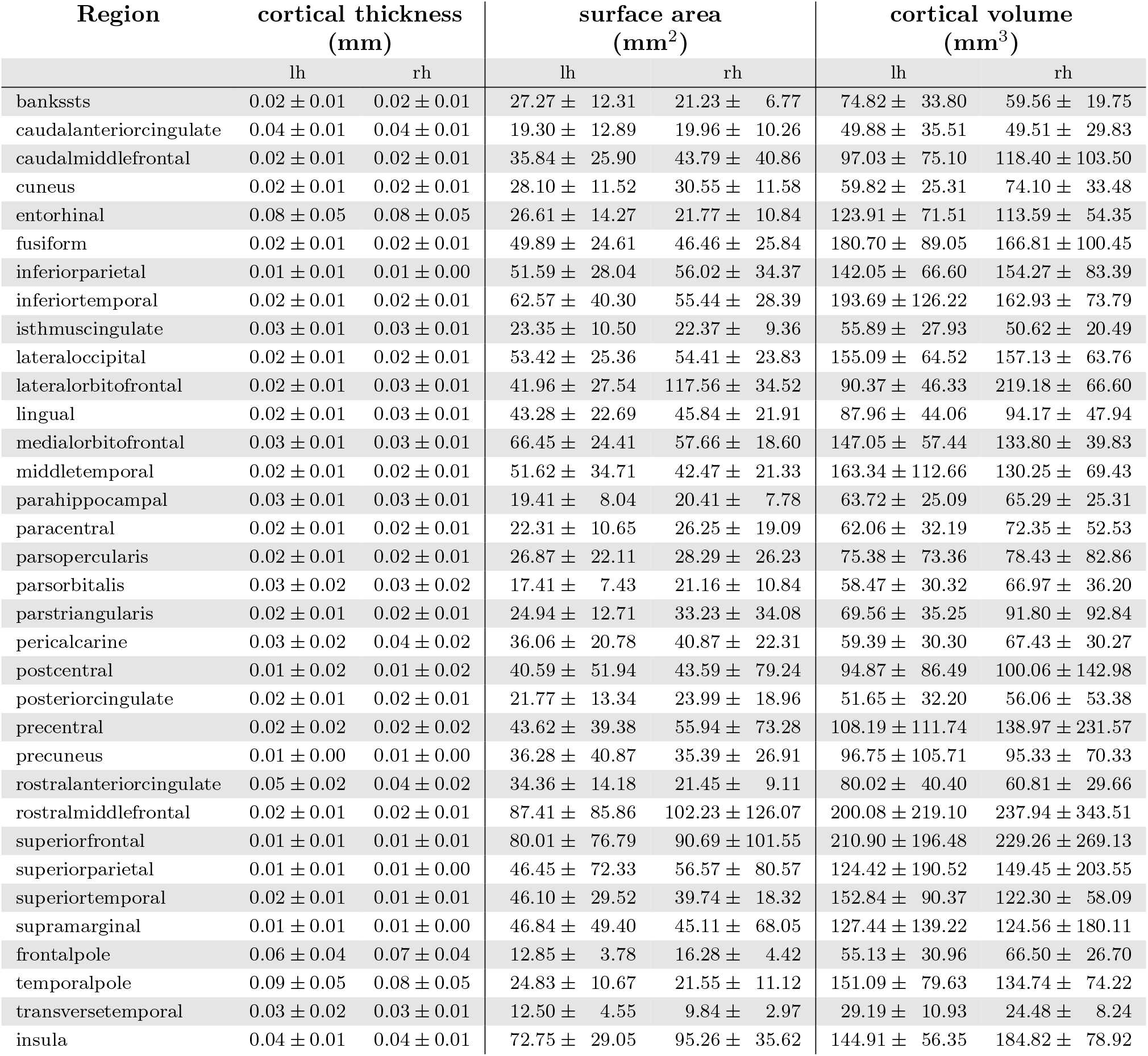
Within-subject standard-deviation average across all subjects for cortical metrics.

**Table S3:**
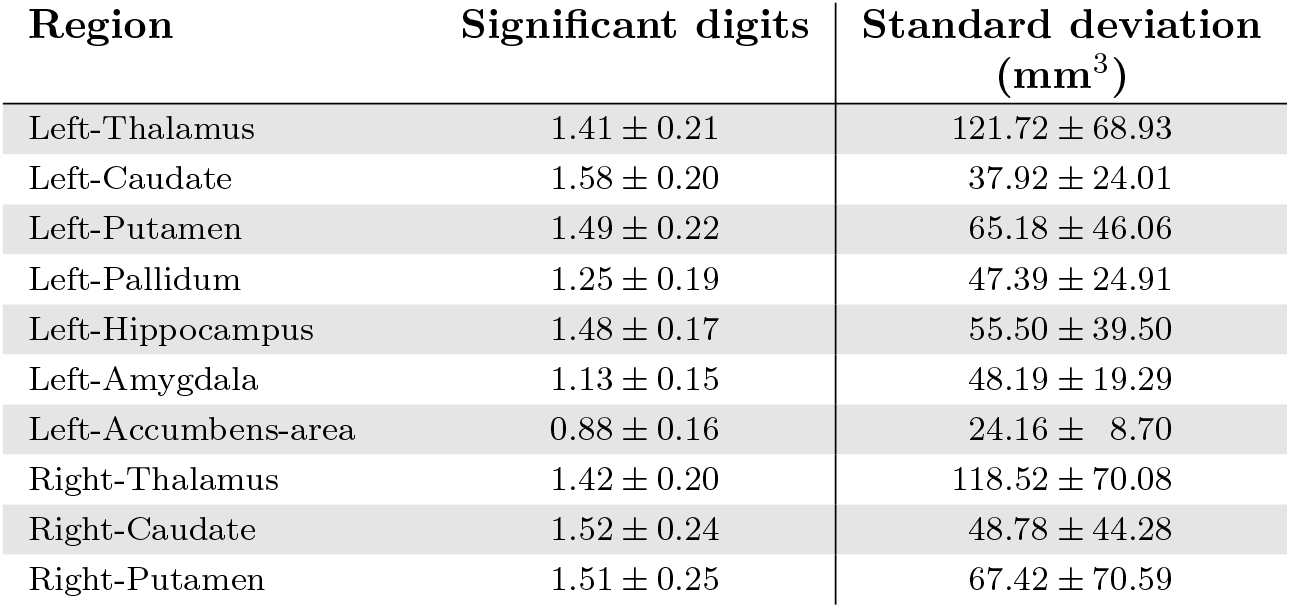
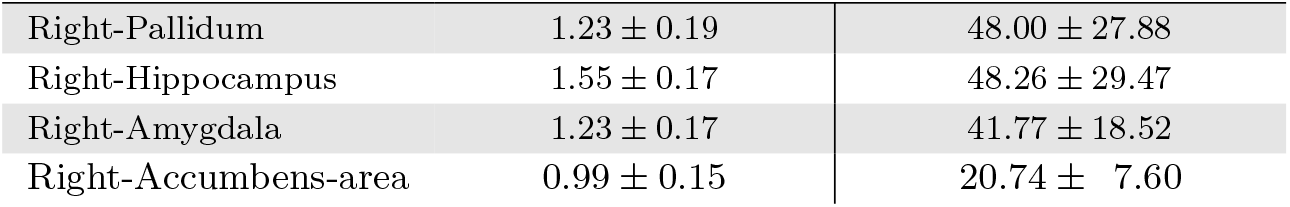
Within-subject significant digits averaged across all subjects for subcortical volumes.

| Region | Significant digits | Standard deviation<br>(mm<sup>3</sup>) |
| --- | --- | --- |
| Left-Thalamus | 1.41 ± 0.21 | 121.72 ± 68.93 |
| Left-Caudate | 1.58 ± 0.20 | 37.92 ± 24.01 |
| Left-Putamen | 1.49 ± 0.22 | 65.18 ± 46.06 |
| Left-Pallidum | 1.25 ± 0.19 | 47.39 ± 24.91 |
| Left-Hippocampus | 1.48 ± 0.17 | 55.50 ± 39.50 |
| Left-Amygdala | 1.13 ± 0.15 | 48.19 ± 19.29 |
| Left-Accumbens-area | 0.88 ± 0.16 | 24.16 ± 8.70 |
| Right-Thalamus | 1.42 ± 0.20 | 118.52 ± 70.08 |
| Right-Caudate | 1.52 ± 0.24 | 48.78 ± 44.28 |
| Right-Putamen | 1.51 ± 0.25 | 67.42 ± 70.59 |
| Right-Pallidum | 1.23 ± 0.19 | 48.00 ± 27.88 |
| Right-Hippocampus | 1.55 ± 0.17 | 48.26 ± 29.47 |
| Right-Amygdala | 1.23 ± 0.17 | 41.77 ± 18.52 |
| Right-Accumbens-area | 0.99 ± 0.15 | 20.74 ± 7.60 |

**Table S4:** Summary of executions failures and excluded subjects. To standardize the sample, we keep 26 repetitions per subject/visits pair. Subject/visit pairs with less than 26 repetitions were excluded (12 subjects).

| Stage | Number of rejected repetitions | Total number of repetitions |
| --- | --- | --- |
| Cluster failure | 1246 (5.80%) | 21488 |
| FreeSurfer failure | 68 (0.33%) | 21488 |
| QC failure | 319 (1.48%) | 21488 |
| Total | 1633 (7.60%) | 21488 |

### S3 Numerical variability of statistical inference: additional information

We evaluated the stability of statistical inference by quantifying the frequency of significance flipping across 26 MCA repetitions (Table S5). The analysis revealed that longitudinal processing introduces considerably more variability than cross-sectional analysis, particularly for cortical metrics. Notably, partial correlation analysis exhibited higher susceptibility to variability in cortical volume (53% of regions unstable) and area (38% unstable) compared to ANCOVA.

To investigate whether longitudinal processing amplifies numerical variability, we compared the dispersion of the test statistics (*F* -values for ANCOVA, correlation coefficients *r* for partial correlation) in longitudinal versus cross-sectional pipelines. We used a one-sided Ansari-Bradley test on mean-centered distributions to test the specific hypothesis that longitudinal processing has significantly higher numerical variance compared to the cross-sectional (Tables S7 and S8).

For subcortical structures (Table S7), partial correlation yielded significantly higher variance in the longitudinal setting for nearly all regions (13/14), whereas ANCOVA remained largely stable, with only the caudate showing significant variance inflation. This trend was mirrored in cortical regions (Table S8); partial correlation analysis consistently demonstrated significantly greater variance longitudinally across volume, thickness, and surface area. In contrast, ANCOVA showed significant variance differences in far fewer regions.

**Table S5:** Frequency of significance flipping across 26 MCA repetitions. Values report the number (and percentage) of unstable regions for each metric and analysis, for baseline and longitudinal settings.

| Metric | ANCOVA |  | Partial correlation |  |
| --- | --- | --- | --- | --- |
|  | Baseline | Longitudinal | Baseline | Longitudinal |
| Cortical Area (68 regions) | 18 (27%) | 36 (53%) | 4 (6%) | 26 (38%) |
| Cortical Thickness (68 regions) | 11 (16%) | 9 (13%) | 19 (28%) | 17 (25%) |
| Cortical Volume (68 regions) | 14 (20%) | 20 (29%) | 8 (12%) | 36 (53%) |
| Subcortical Volume (14 regions) | 4 (29%) | 2 (14%) | 4 (29%) | 5 (36%) |

**Table S6:**
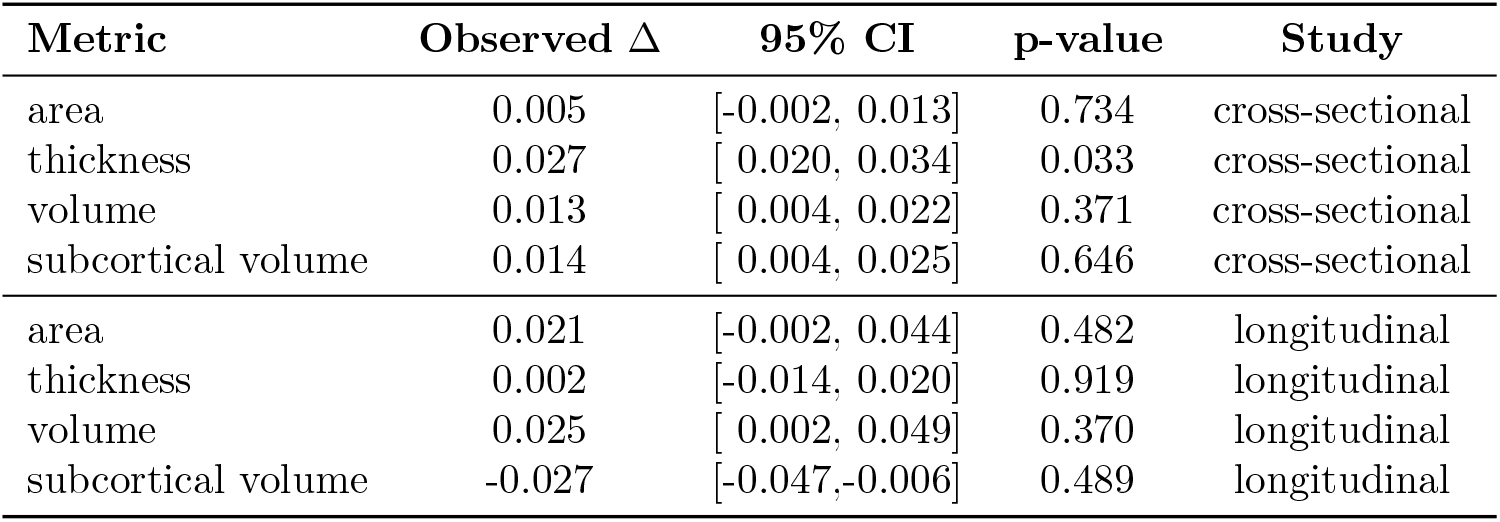
Permutation test comparison of *ν*_npv_ between HC and PD. No significance detected after Bonferroni correction (*α* = 0.05*/*8). Values are observed differences (PD − HC). 95% CIs from percentile bootstrap.

| Metric | Observed $\Delta$ | 95% CI | p-value | Study |
| --- | --- | --- | --- | --- |
| area | 0.005 | [-0.002, 0.013] | 0.734 | cross-sectional |
| thickness | 0.027 | [ 0.020, 0.034] | 0.033 | cross-sectional |
| volume | 0.013 | [ 0.004, 0.022] | 0.371 | cross-sectional |
| subcortical volume | 0.014 | [ 0.004, 0.025] | 0.646 | cross-sectional |
| area | 0.021 | [-0.002, 0.044] | 0.482 | longitudinal |
| thickness | 0.002 | [-0.014, 0.020] | 0.919 | longitudinal |
| volume | 0.025 | [ 0.002, 0.049] | 0.370 | longitudinal |
| subcortical volume | -0.027 | [-0.047,-0.006] | 0.489 | longitudinal |

**Table S7:** Longitudinal versus cross-sectional variance of test statistics in **subcortical** regions. One-sided Ansari-Bradley test comparing cross-sectional vs. longitudinal variances of the test statistics (*F* -values for ANCOVA, *r*-values for partial correlation). *W* = Ansari-Bradley statistic, *p* = FDR-adjusted p-value; an asterisk (∗*p*_FDR_ *<* 0.05) marks regions where the test statistic is significantly *more* variable longitudinally than cross-sectionally, i.e. where longitudinal processing inflates numerical variance. L/R = Left/Right.

| Region | ANCOVA |  | Partial Corr |  |
| --- | --- | --- | --- | --- |
|  | W | p | W | p |
| L-Thalamus | 248 | 1.00 | 465 | 6.1e-6* |
| L-Caudate | 501 | 3.1e-10* | 459 | 2.0e-5* |
| L-Putamen | 327 | 0.81 | 491 | 9.6e-9* |
| L-Pallidum | 264 | 1.00 | 449 | 1.1e-4* |
| L-Hippocampus | 314 | 0.91 | 428 | 2.2e-3* |
| L-Amygdala | 261 | 1.00 | 476 | 5.5e-7* |
| L-Accumbens | 265 | 1.00 | 442 | 3.3e-4* |
| R-Thalamus | 281 | 1.00 | 441 | 3.9e-4* |
| R-Caudate | 485 | 5.5e-8* | 478 | 3.4e-7* |
| R-Putamen | 294 | 0.98 | 489 | 1.7e-8* |
| R-Pallidum | 212 | 1.00 | 469 | 2.7e-6* |
| R-Hippocampus | 335 | 0.73 | 493 | 5.1e-9* |
| R-Amygdala | 316 | 0.90 | 408 | 1.9e-2* |
| R-Accumbens | 214 | 1.00 | 418 | 7.0e-3* |

**Table S8:**
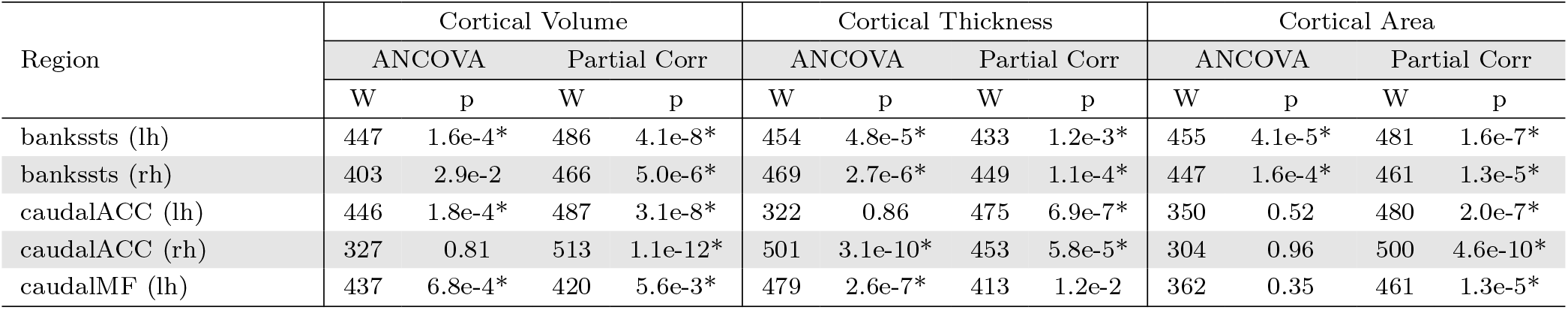
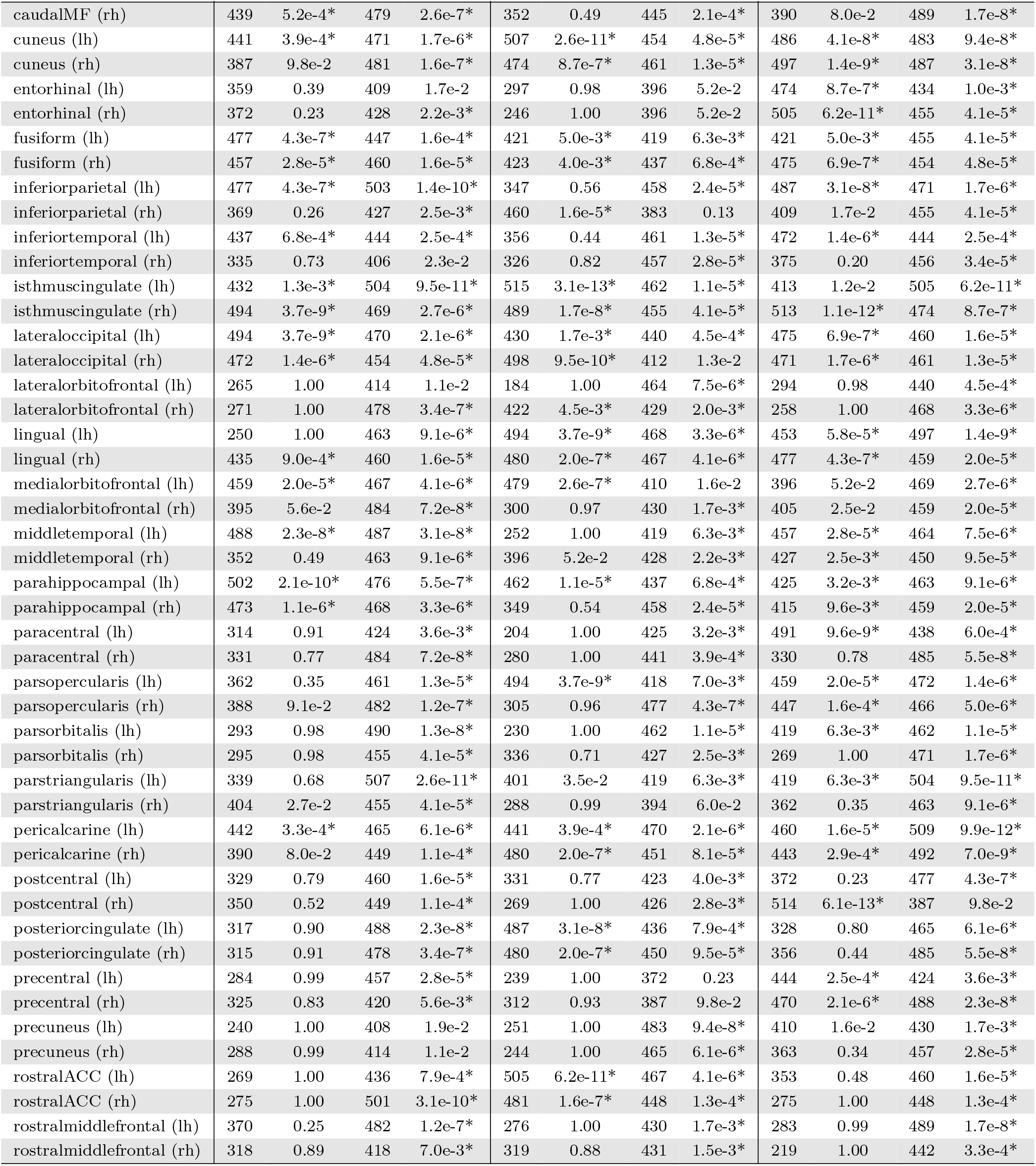
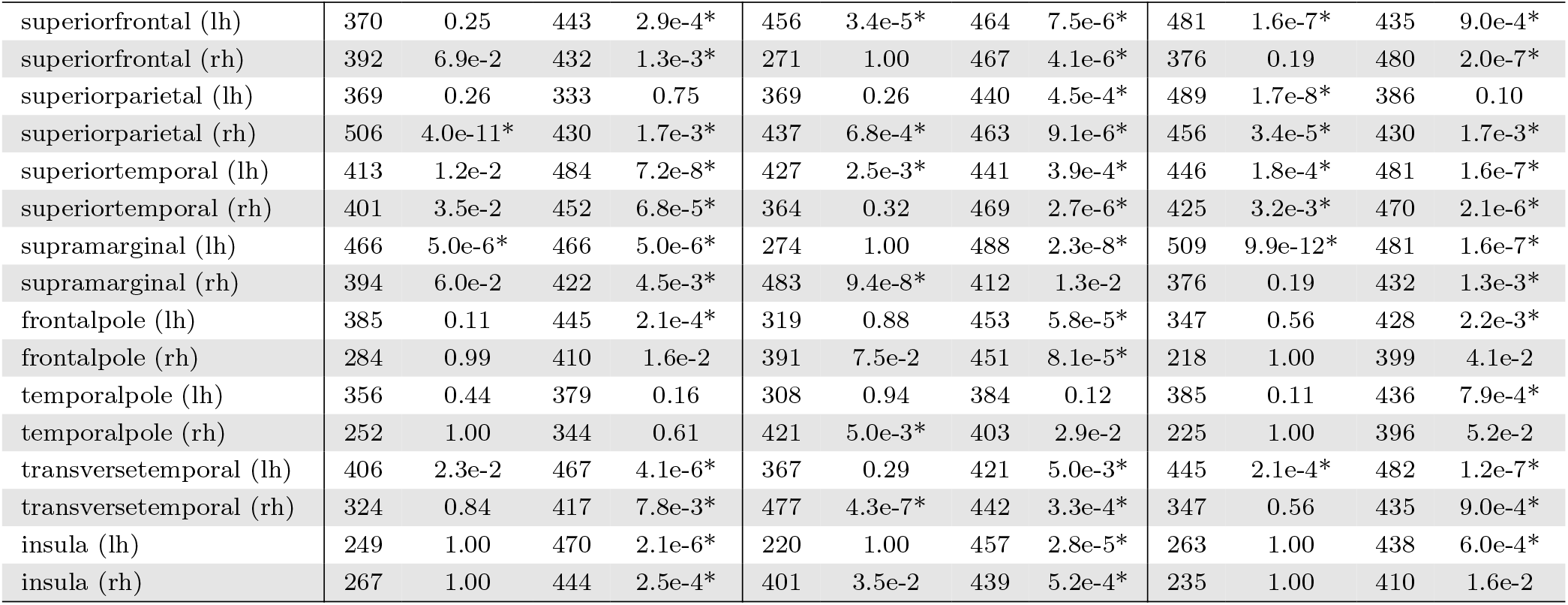
Longitudinal versus cross-sectional variance of test statistics in **cortical** regions (one-sided Ansari-Bradley test). *W* = Ansari-Bradley statistic, *p* = FDR-adjusted p-value; an asterisk (* *p <* 0.05) marks regions where the test statistic is significantly more variable longitudinally than cross-sectionally, i.e. where longitudinal processing inflates numerical variance. lh/rh = left/right hemisphere; ACC = anterior cingulate cortex, MF = middle frontal.

| Region | Cortical Volume |  |  |  | Cortical Thickness |  |  |  | Cortical Area |  |  |  |
| --- | --- | --- | --- | --- | --- | --- | --- | --- | --- | --- | --- | --- |
|  | ANCOVA |  | Partial Corr |  | ANCOVA |  | Partial Corr |  | ANCOVA |  | Partial Corr |  |
|  | W | p | W | p | W | p | W | p | W | p | W | p |
| bankssts (lh) | 447 | 1.6e-4* | 486 | 4.1e-8* | 454 | 4.8e-5* | 433 | 1.2e-3* | 455 | 4.1e-5* | 481 | 1.6e-7* |
| bankssts (rh) | 403 | 2.9e-2 | 466 | 5.0e-6* | 469 | 2.7e-6* | 449 | 1.1e-4* | 447 | 1.6e-4* | 461 | 1.3e-5* |
| caudalACC (lh) | 446 | 1.8e-4* | 487 | 3.1e-8* | 322 | 0.86 | 475 | 6.9e-7* | 350 | 0.52 | 480 | 2.0e-7* |
| caudalACC (rh) | 327 | 0.81 | 513 | 1.1e-12* | 501 | 3.1e-10* | 453 | 5.8e-5* | 304 | 0.96 | 500 | 4.6e-10* |
| caudalMF (lh) | 437 | 6.8e-4* | 420 | 5.6e-3* | 479 | 2.6e-7* | 413 | 1.2e-2 | 362 | 0.35 | 461 | 1.3e-5* |
|  | ANCOVA |  | Partial Corr |  | ANCOVA |  | Partial Corr |  | ANCOVA |  | Partial Corr |  |
|  | W | p | W | p | W | p | W | p | W | p | W | p |
| caudalMF (rh) | 439 | 5.2e-4* | 479 | 2.6e-7* | 352 | 0.49 | 445 | 2.1e-4* | 390 | 8.0e-2 | 489 | 1.7e-8* |
| cuneus (lh) | 441 | 3.9e-4* | 471 | 1.7e-6* | 507 | 2.6e-11* | 454 | 4.8e-5* | 486 | 4.1e-8* | 483 | 9.4e-8* |
| cuneus (rh) | 387 | 9.8e-2 | 481 | 1.6e-7* | 474 | 8.7e-7* | 461 | 1.3e-5* | 497 | 1.4e-9* | 487 | 3.1e-8* |
| entorhinal (lh) | 359 | 0.39 | 409 | 1.7e-2 | 297 | 0.98 | 396 | 5.2e-2 | 474 | 8.7e-7* | 434 | 1.0e-3* |
| entorhinal (rh) | 372 | 0.23 | 428 | 2.2e-3* | 246 | 1.00 | 396 | 5.2e-2 | 505 | 6.2e-11* | 455 | 4.1e-5* |
| fusiform (lh) | 477 | 4.3e-7* | 447 | 1.6e-4* | 421 | 5.0e-3* | 419 | 6.3e-3* | 421 | 5.0e-3* | 455 | 4.1e-5* |
| fusiform (rh) | 457 | 2.8e-5* | 460 | 1.6e-5* | 423 | 4.0e-3* | 437 | 6.8e-4* | 475 | 6.9e-7* | 454 | 4.8e-5* |
| inferiorparietal (lh) | 477 | 4.3e-7* | 503 | 1.4e-10* | 347 | 0.56 | 458 | 2.4e-5* | 487 | 3.1e-8* | 471 | 1.7e-6* |
| inferiorparietal (rh) | 369 | 0.26 | 427 | 2.5e-3* | 460 | 1.6e-5* | 383 | 0.13 | 409 | 1.7e-2 | 455 | 4.1e-5* |
| inferiortemporal (lh) | 437 | 6.8e-4* | 444 | 2.5e-4* | 356 | 0.44 | 461 | 1.3e-5* | 472 | 1.4e-6* | 444 | 2.5e-4* |
| inferiortemporal (rh) | 335 | 0.73 | 406 | 2.3e-2 | 326 | 0.82 | 457 | 2.8e-5* | 375 | 0.20 | 456 | 3.4e-5* |
| isthmuscingulate (lh) | 432 | 1.3e-3* | 504 | 9.5e-11* | 515 | 3.1e-13* | 462 | 1.1e-5* | 413 | 1.2e-2 | 505 | 6.2e-11* |
| isthmuscingulate (rh) | 494 | 3.7e-9* | 469 | 2.7e-6* | 489 | 1.7e-8* | 455 | 4.1e-5* | 513 | 1.1e-12* | 474 | 8.7e-7* |
| lateraloccipital (lh) | 494 | 3.7e-9* | 470 | 2.1e-6* | 430 | 1.7e-3* | 440 | 4.5e-4* | 475 | 6.9e-7* | 460 | 1.6e-5* |
| lateraloccipital (rh) | 472 | 1.4e-6* | 454 | 4.8e-5* | 498 | 9.5e-10* | 412 | 1.3e-2 | 471 | 1.7e-6* | 461 | 1.3e-5* |
| lateralorbitofrontal (lh) | 265 | 1.00 | 414 | 1.1e-2 | 184 | 1.00 | 464 | 7.5e-6* | 294 | 0.98 | 440 | 4.5e-4* |
| lateralorbitofrontal (rh) | 271 | 1.00 | 478 | 3.4e-7* | 422 | 4.5e-3* | 429 | 2.0e-3* | 258 | 1.00 | 468 | 3.3e-6* |
| lingual (lh) | 250 | 1.00 | 463 | 9.1e-6* | 494 | 3.7e-9* | 468 | 3.3e-6* | 453 | 5.8e-5* | 497 | 1.4e-9* |
| lingual (rh) | 435 | 9.0e-4* | 460 | 1.6e-5* | 480 | 2.0e-7* | 467 | 4.1e-6* | 477 | 4.3e-7* | 459 | 2.0e-5* |
| medialorbitofrontal (lh) | 459 | 2.0e-5* | 467 | 4.1e-6* | 479 | 2.6e-7* | 410 | 1.6e-2 | 396 | 5.2e-2 | 469 | 2.7e-6* |
| medialorbitofrontal (rh) | 395 | 5.6e-2 | 484 | 7.2e-8* | 300 | 0.97 | 430 | 1.7e-3* | 405 | 2.5e-2 | 459 | 2.0e-5* |
| middletemporal (lh) | 488 | 2.3e-8* | 487 | 3.1e-8* | 252 | 1.00 | 419 | 6.3e-3* | 457 | 2.8e-5* | 464 | 7.5e-6* |
| middletemporal (rh) | 352 | 0.49 | 463 | 9.1e-6* | 396 | 5.2e-2 | 428 | 2.2e-3* | 427 | 2.5e-3* | 450 | 9.5e-5* |
| parahippocampal (lh) | 502 | 2.1e-10* | 476 | 5.5e-7* | 462 | 1.1e-5* | 437 | 6.8e-4* | 425 | 3.2e-3* | 463 | 9.1e-6* |
| parahippocampal (rh) | 473 | 1.1e-6* | 468 | 3.3e-6* | 349 | 0.54 | 458 | 2.4e-5* | 415 | 9.6e-3* | 459 | 2.0e-5* |
| paracentral (lh) | 314 | 0.91 | 424 | 3.6e-3* | 204 | 1.00 | 425 | 3.2e-3* | 491 | 9.6e-9* | 438 | 6.0e-4* |
| paracentral (rh) | 331 | 0.77 | 484 | 7.2e-8* | 280 | 1.00 | 441 | 3.9e-4* | 330 | 0.78 | 485 | 5.5e-8* |
| parsopercularis (lh) | 362 | 0.35 | 461 | 1.3e-5* | 494 | 3.7e-9* | 418 | 7.0e-3* | 459 | 2.0e-5* | 472 | 1.4e-6* |
| parsopercularis (rh) | 388 | 9.1e-2 | 482 | 1.2e-7* | 305 | 0.96 | 477 | 4.3e-7* | 447 | 1.6e-4* | 466 | 5.0e-6* |
| parsorbitalis (lh) | 293 | 0.98 | 490 | 1.3e-8* | 230 | 1.00 | 462 | 1.1e-5* | 419 | 6.3e-3* | 462 | 1.1e-5* |
| parsorbitalis (rh) | 295 | 0.98 | 455 | 4.1e-5* | 336 | 0.71 | 427 | 2.5e-3* | 269 | 1.00 | 471 | 1.7e-6* |
| parstriangularis (lh) | 339 | 0.68 | 507 | 2.6e-11* | 401 | 3.5e-2 | 419 | 6.3e-3* | 419 | 6.3e-3* | 504 | 9.5e-11* |
| parstriangularis (rh) | 404 | 2.7e-2 | 455 | 4.1e-5* | 288 | 0.99 | 394 | 6.0e-2 | 362 | 0.35 | 463 | 9.1e-6* |
| pericalcarine (lh) | 442 | 3.3e-4* | 465 | 6.1e-6* | 441 | 3.9e-4* | 470 | 2.1e-6* | 460 | 1.6e-5* | 509 | 9.9e-12* |
| pericalcarine (rh) | 390 | 8.0e-2 | 449 | 1.1e-4* | 480 | 2.0e-7* | 451 | 8.1e-5* | 443 | 2.9e-4* | 492 | 7.0e-9* |
| postcentral (lh) | 329 | 0.79 | 460 | 1.6e-5* | 331 | 0.77 | 423 | 4.0e-3* | 372 | 0.23 | 477 | 4.3e-7* |
| postcentral (rh) | 350 | 0.52 | 449 | 1.1e-4* | 269 | 1.00 | 426 | 2.8e-3* | 514 | 6.1e-13* | 387 | 9.8e-2 |
| posteriorcingulate (lh) | 317 | 0.90 | 488 | 2.3e-8* | 487 | 3.1e-8* | 436 | 7.9e-4* | 328 | 0.80 | 465 | 6.1e-6* |
| posteriorcingulate (rh) | 315 | 0.91 | 478 | 3.4e-7* | 480 | 2.0e-7* | 450 | 9.5e-5* | 356 | 0.44 | 485 | 5.5e-8* |
| precentral (lh) | 284 | 0.99 | 457 | 2.8e-5* | 239 | 1.00 | 372 | 0.23 | 444 | 2.5e-4* | 424 | 3.6e-3* |
| precentral (rh) | 325 | 0.83 | 420 | 5.6e-3* | 312 | 0.93 | 387 | 9.8e-2 | 470 | 2.1e-6* | 488 | 2.3e-8* |
| precuneus (lh) | 240 | 1.00 | 408 | 1.9e-2 | 251 | 1.00 | 483 | 9.4e-8* | 410 | 1.6e-2 | 430 | 1.7e-3* |
| precuneus (rh) | 288 | 0.99 | 414 | 1.1e-2 | 244 | 1.00 | 465 | 6.1e-6* | 363 | 0.34 | 457 | 2.8e-5* |
| rostralACC (lh) | 269 | 1.00 | 436 | 7.9e-4* | 505 | 6.2e-11* | 467 | 4.1e-6* | 353 | 0.48 | 460 | 1.6e-5* |
| rostralACC (rh) | 275 | 1.00 | 501 | 3.1e-10* | 481 | 1.6e-7* | 448 | 1.3e-4* | 275 | 1.00 | 448 | 1.3e-4* |
| rostralmiddlefrontal (lh) | 370 | 0.25 | 482 | 1.2e-7* | 276 | 1.00 | 430 | 1.7e-3* | 283 | 0.99 | 489 | 1.7e-8* |
| rostralmiddlefrontal (rh) | 318 | 0.89 | 418 | 7.0e-3* | 319 | 0.88 | 431 | 1.5e-3* | 219 | 1.00 | 442 | 3.3e-4* |
|  | ANCOVA |  | Partial Corr |  | ANCOVA |  | Partial Corr |  | ANCOVA |  | Partial Corr |  |
|  | W | p | W | p | W | p | W | p | W | p | W | p |
| superiorfrontal (lh) | 370 | 0.25 | 443 | 2.9e-4* | 456 | 3.4e-5* | 464 | 7.5e-6* | 481 | 1.6e-7* | 435 | 9.0e-4* |
| superiorfrontal (rh) | 392 | 6.9e-2 | 432 | 1.3e-3* | 271 | 1.00 | 467 | 4.1e-6* | 376 | 0.19 | 480 | 2.0e-7* |
| superiorparietal (lh) | 369 | 0.26 | 333 | 0.75 | 369 | 0.26 | 440 | 4.5e-4* | 489 | 1.7e-8* | 386 | 0.10 |
| superiorparietal (rh) | 506 | 4.0e-11* | 430 | 1.7e-3* | 437 | 6.8e-4* | 463 | 9.1e-6* | 456 | 3.4e-5* | 430 | 1.7e-3* |
| superiortemporal (lh) | 413 | 1.2e-2 | 484 | 7.2e-8* | 427 | 2.5e-3* | 441 | 3.9e-4* | 446 | 1.8e-4* | 481 | 1.6e-7* |
| superiortemporal (rh) | 401 | 3.5e-2 | 452 | 6.8e-5* | 364 | 0.32 | 469 | 2.7e-6* | 425 | 3.2e-3* | 470 | 2.1e-6* |
| supramarginal (lh) | 466 | 5.0e-6* | 466 | 5.0e-6* | 274 | 1.00 | 488 | 2.3e-8* | 509 | 9.9e-12* | 481 | 1.6e-7* |
| supramarginal (rh) | 394 | 6.0e-2 | 422 | 4.5e-3* | 483 | 9.4e-8* | 412 | 1.3e-2 | 376 | 0.19 | 432 | 1.3e-3* |
| frontalpole (lh) | 385 | 0.11 | 445 | 2.1e-4* | 319 | 0.88 | 453 | 5.8e-5* | 347 | 0.56 | 428 | 2.2e-3* |
| frontalpole (rh) | 284 | 0.99 | 410 | 1.6e-2 | 391 | 7.5e-2 | 451 | 8.1e-5* | 218 | 1.00 | 399 | 4.1e-2 |
| temporalpole (lh) | 356 | 0.44 | 379 | 0.16 | 308 | 0.94 | 384 | 0.12 | 385 | 0.11 | 436 | 7.9e-4* |
| temporalpole (rh) | 252 | 1.00 | 344 | 0.61 | 421 | 5.0e-3* | 403 | 2.9e-2 | 225 | 1.00 | 396 | 5.2e-2 |
| transversetemporal (lh) | 406 | 2.3e-2 | 467 | 4.1e-6* | 367 | 0.29 | 421 | 5.0e-3* | 445 | 2.1e-4* | 482 | 1.2e-7* |
| transversetemporal (rh) | 324 | 0.84 | 417 | 7.8e-3* | 477 | 4.3e-7* | 442 | 3.3e-4* | 347 | 0.56 | 435 | 9.0e-4* |
| insula (lh) | 249 | 1.00 | 470 | 2.1e-6* | 220 | 1.00 | 457 | 2.8e-5* | 263 | 1.00 | 438 | 6.0e-4* |
| insula (rh) | 267 | 1.00 | 444 | 2.5e-4* | 401 | 3.5e-2 | 439 | 5.2e-4* | 235 | 1.00 | 410 | 1.6e-2 |

For both group comparisons and correlation analyses, statistical outcomes varied substantially across the 26 Monte Carlo Arithmetic (MCA) repetitions (Figures S4 and S5). For cortical area (68 regions; Figure S4), significance flipped in 31% of all (regions, analysis) pairs, indicating frequent inconsistencies across repetitions. For cortical volume (68 regions; Figure S5), 29% of pairs were similarly unstable. These fluctuations demonstrate that numerical noise alone can alter downstream statistical inference in structural MRI analyses of PD.

### S4 Distribution of statistical test coefficients

This section provides the per-metric coefficient distributions underlying the substantial run-to-run variability of test statistics reported in Results, Section 2.1. Figures S8, S9, S10 and S11 show the distribution of partial correlation coefficients and F-statistics for subcortical volume, cortical thickness, cortical surface area and cortical volume measures, across all subjects and regions. Red triangles indicate the unperturbed (IEEE-754) run for reference. For all analyses, the unperturbed (IEEE-754) result is included in the range of numerically perturbed results, which supports the validity of the numerical perturbation.

### S5 Numerical validation of numerical variability propagation

To numerically validate the proposed variability-propagation framework, we compared analytical standard-deviation estimates derived from the closed-form expressions reported in Table 1 with empirical estimates obtained through repeated Monte Carlo Arithmetic (MCA) perturbations of the full processing pipelines. For each statistical test (Cohen’s *d*, two-sample *t*-statistics, partial correlation coefficients, and ANCOVA *F* -statistics), as well as their associated *p*-values (except for Cohen’s *d*), we computed the standard deviation across MCA repetitions and quantified the relative error between sampled and analytical estimates. This comparison was performed across all cortical and subcortical regions, imaging metrics, and for both cross-sectional and longitudinal study designs.

Overall, the analytical estimates captured the order of magnitude and regional structure of the sampled variability, although the agreement is not uniformly centered at zero across all configurations. In cross-sectional analyses, the average relative error was -0.014 (± 0.160) for the two-sample *t*-test, - 0.014 (±0.160) for Cohen’s *d*, 0.015 (±0.200) for ANCOVA *F* -statistics, and 0.079 (±0.260) for partial correlations. These distributions remain visually concentrated around zero with limited skewness. Longitudinal analyses yielded similarly small mean discrepancies for the *t*-test (-0.010 ±0.145) and Cohen’s *d* (-0.003 ±0.146), whereas partial correlations (-0.002 ±0.184) and especially ANCOVA (-0.184 ±0.207) display a broader spread and a clearer negative displacement. Inspection of the violin plots (Fig. S12) shows that this shift is not uniform across metrics: cortical thickness and area tend to remain closer to zero, while cortical and subcortical volumes contribute more strongly to the negative tail.

A similar but more dispersed pattern was observed for *p*-values. In cross-sectional analyses, mean relative errors were -0.071 (±0.180) for the two-sample *t*-test, 0.106 (±0.323) for ANCOVA, and 0.032 (±0.277) for partial correlations, with most densities overlapping zero but exhibiting heavier tails than their statistic counterparts. Longitudinal analyses showed negative mean relative errors of -0.290 (±0.156) for the *t*-test, -0.286 (±0.159) for ANCOVA, and -0.269 (±0.174) for partial correlations. The violin plots (Fig. S13) indicate that these negative tendencies arise from broadened and asymmetric distributions with several metrics maintaining substantial overlap with zero.

These results indicate that the analytical variability model provides a faithful approximation of numerically induced variability in downstream statistical measures, with accuracy depending on both the statistical test and study design. Cross-sectional configurations show near-unbiased and tightly bounded relative errors, whereas longitudinal settings exhibit increased dispersion and negative shifts, especially for partial correlation and *p*-values. Importantly, the error distributions remain largely confined within ±0.5 for the majority of regions and metrics, and Cohen’s *d* consistently demonstrates the highest stability. The longitudinal negative bias reflects a tendency of the analytical model to overestimate variability.

### S6 Numerical validation of flip significance simulations

To assess the validity of the analytical variability framework used to model numerically induced significance flips, we compared simulated estimates derived from closed-form variability expressions with empirical estimates obtained from Monte Carlo Arithmetic (MCA) repetitions. Formally, the proportion of significant tests (positive prediction rate, PPR) is defined as:

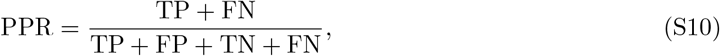

where TP, FP, TN, and FN denote true positives, false positives, true negatives, and false negatives, respectively.

Figure S15 compares standard deviations of *p*-values estimated directly from 26 MCA runs with those predicted by the analytical variability formulas (Table 1). Each point corresponds to a cortical or subcortical brain region. Results show close agreement between sampled and simulated estimates, particularly in the cross-sectional setting. In the longitudinal analyses, simulated standard deviations tend to be slightly larger than sampled values, indicating a conservative bias of the analytical model.

Figure S16 evaluates how this analytical variability propagates to downstream inference by comparing the simulated positive prediction rate (PPR; Eq. S10) with the empirical proportion of significant tests observed across MCA repetitions. Cross-sectional results closely follow the identity line, while longitudinal results show a moderate overestimation of PPR, consistent with the conservative overestimation of *p*-value variability observed in Fig. S15.

Finally, Figure S17 assesses the Beta distribution model used to translate sampled *p*-value variability into PPR estimates. Here, PPR is computed using MCA-derived standard deviations propagated through the Beta model (Eq. 4.4). Both cross-sectional and longitudinal results align closely with the identity line, supporting the suitability of the Beta modeling framework for capturing numerically induced variability in statistical significance.

Figure S14 extends the analysis of Figure 4 by providing a breakdown of significance flip probabilities by statistical test type: Student’s *t*-test (T), ANCOVA (*F* -values), and partial correlation (R). The same overall pattern holds across all three test types: significance flip probability is highest for results near the significance threshold (negative distances for reported significant findings, positive distances for reported non-significant findings) and decreases as results move farther from the threshold.

Partial correlations tend to exhibit slightly higher flip probabilities, consistent with their generally larger numerical variability as quantified by the NPVR framework (Table 1). These results confirm that the widespread numerical fragility observed across the 13 reviewed Parkinson’s disease studies is not specific to any single statistical test type.

**Figure S2:**
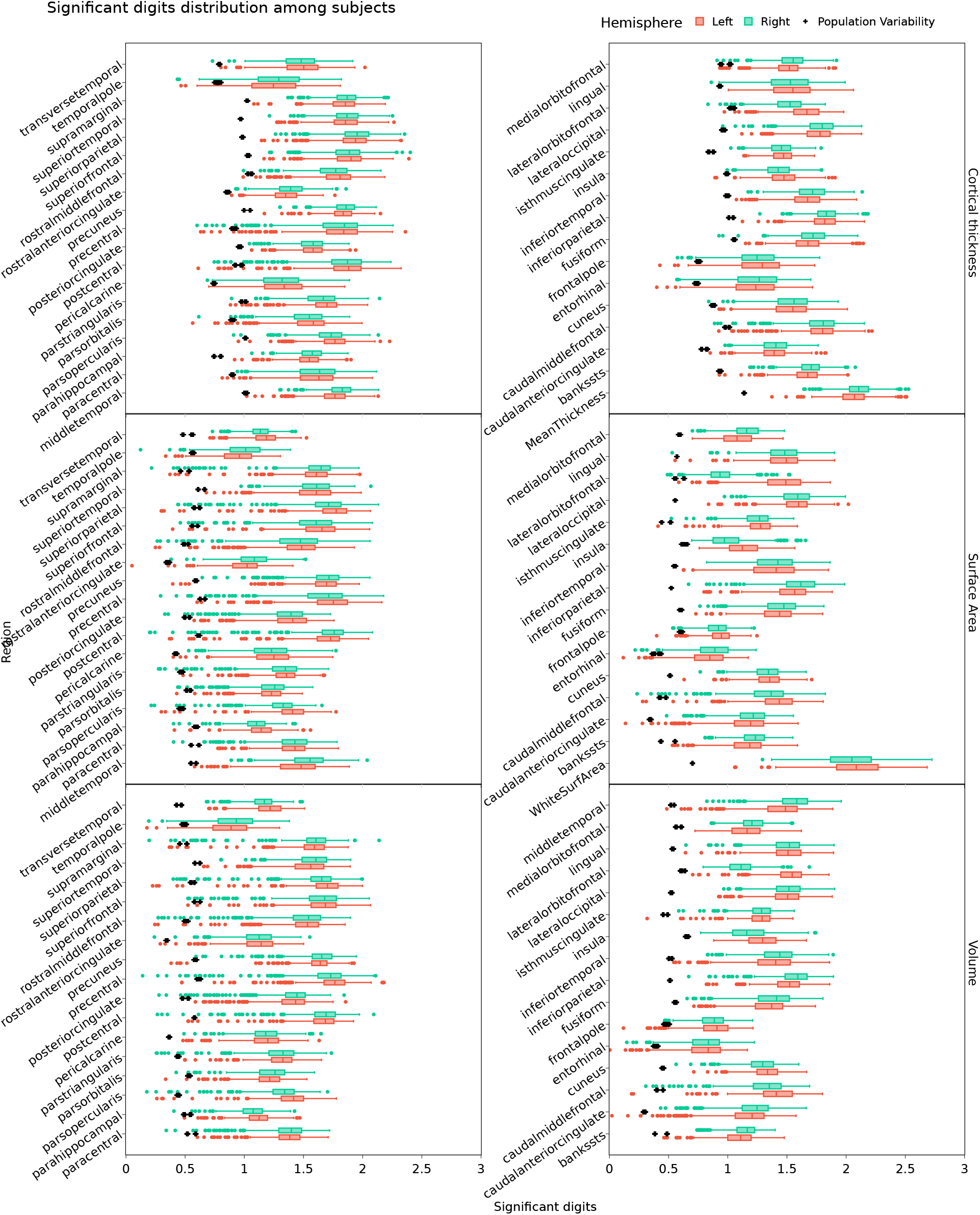
Numerical precision of cortical measures. The number of significant digits for cortical thickness, surface area, and volume across regions. Lower values indicate higher variability. Significant digits for population variability (black cross) remain consistently lower than numerical variability on average, indicating that numerical variability does not dominate population variability.

**Figure S3:**
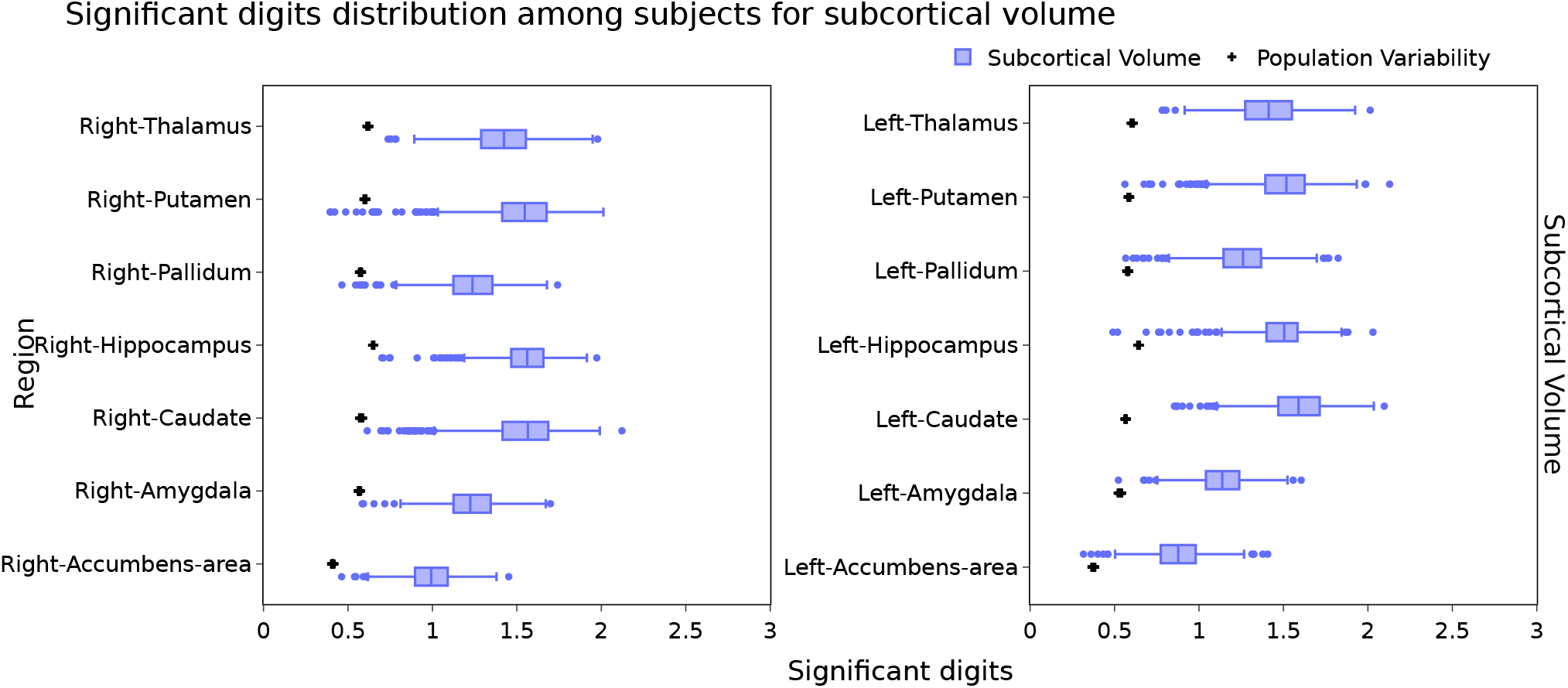
Numerical precision of subcortical volumes. The number of significant digits estimated for each subcortical region, showing comparable variability to cortical volume measures. Significant digits for population variability (black cross) remain consistently lower than numerical variability on average, indicating that numerical variability does not dominate population variability.

**Figure S4:**
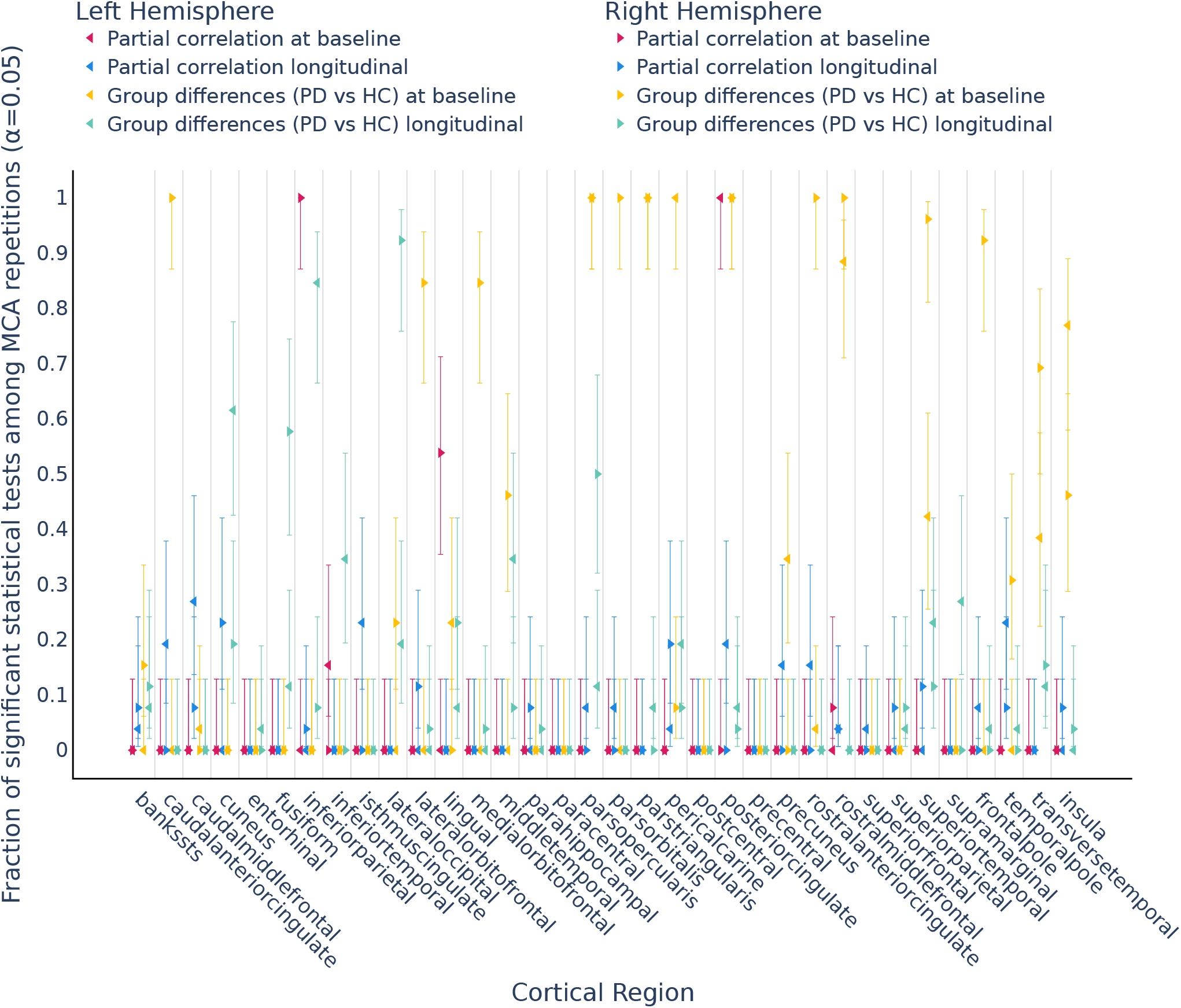
Proportion of significant tests (*p <* 0.05, uncorrected) for cortical surface area across 26 numerical perturbations.

**Figure S5:**
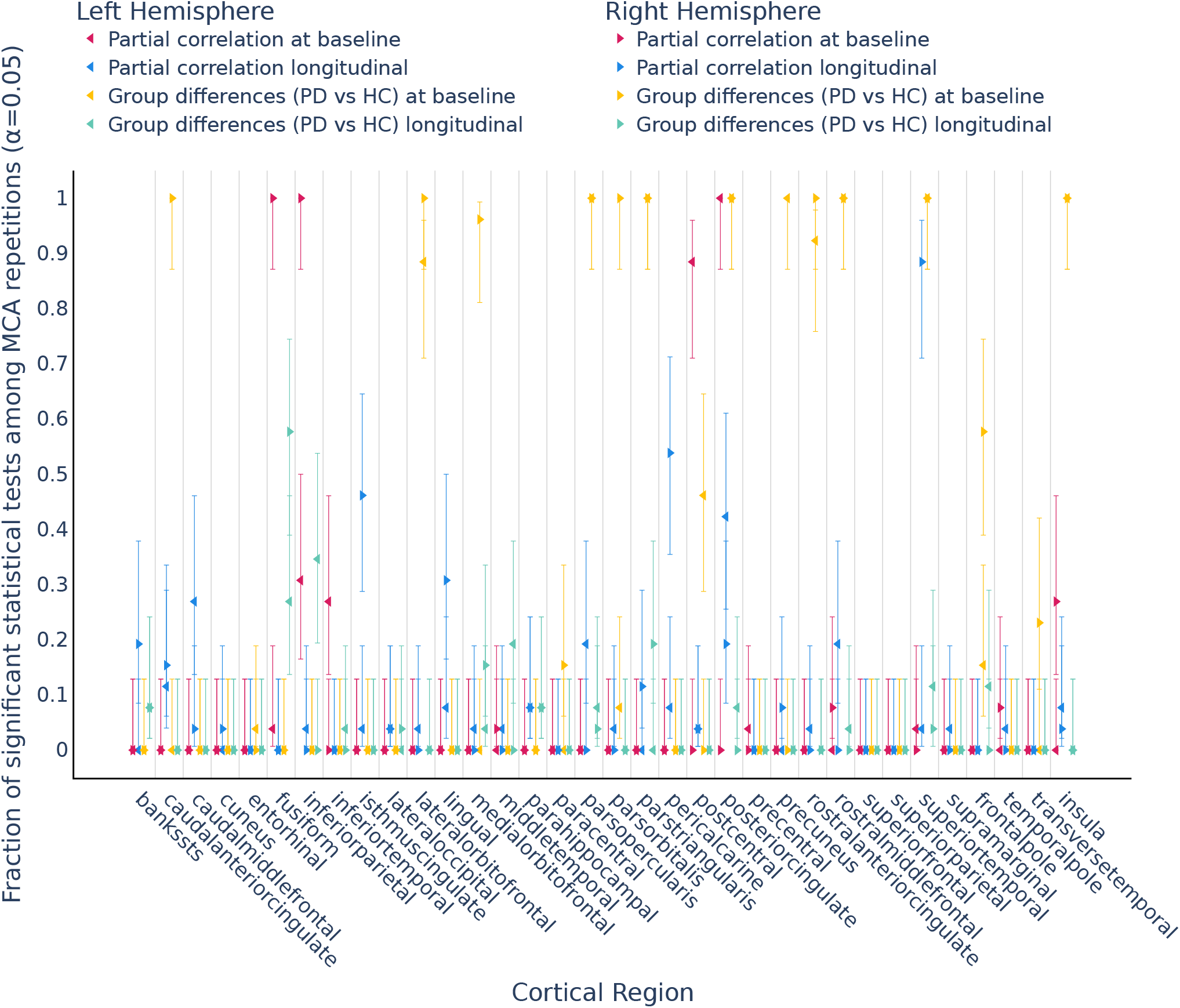
Proportion of significant tests (*p <* 0.05, uncorrected) for cortical volume across 26 numerical perturbations.

**Figure S6:**
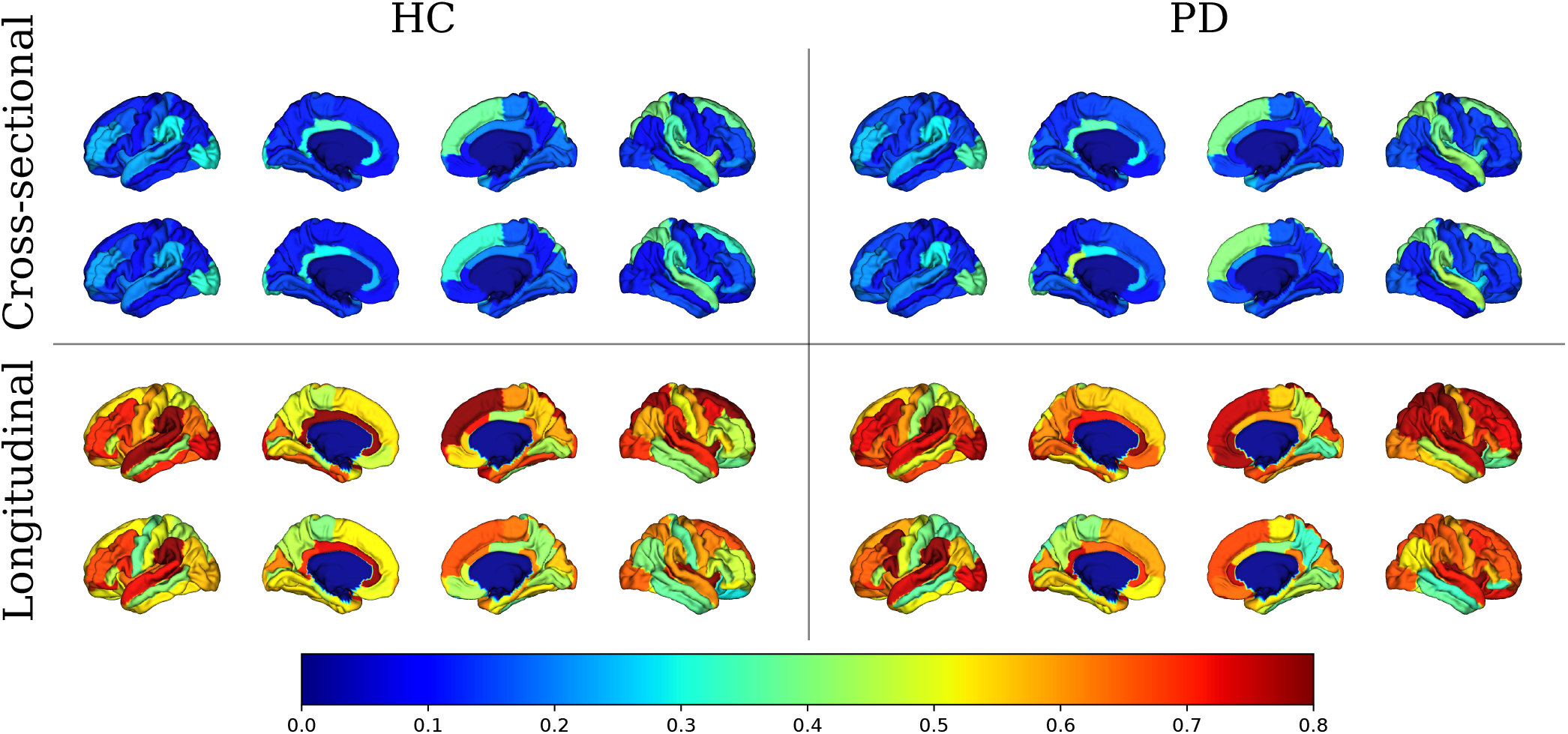
Numerical-Population Variability Ratio (*ν*_npv_) for cortical surface area (top row in each panel) and cortical volume (bottom row in each panel) in healthy controls (HC) and Parkinson’s disease (PD). Higher *ν*_npv_ values indicate higher numerical variability relative to inter-subject variability.

**Figure S7:**
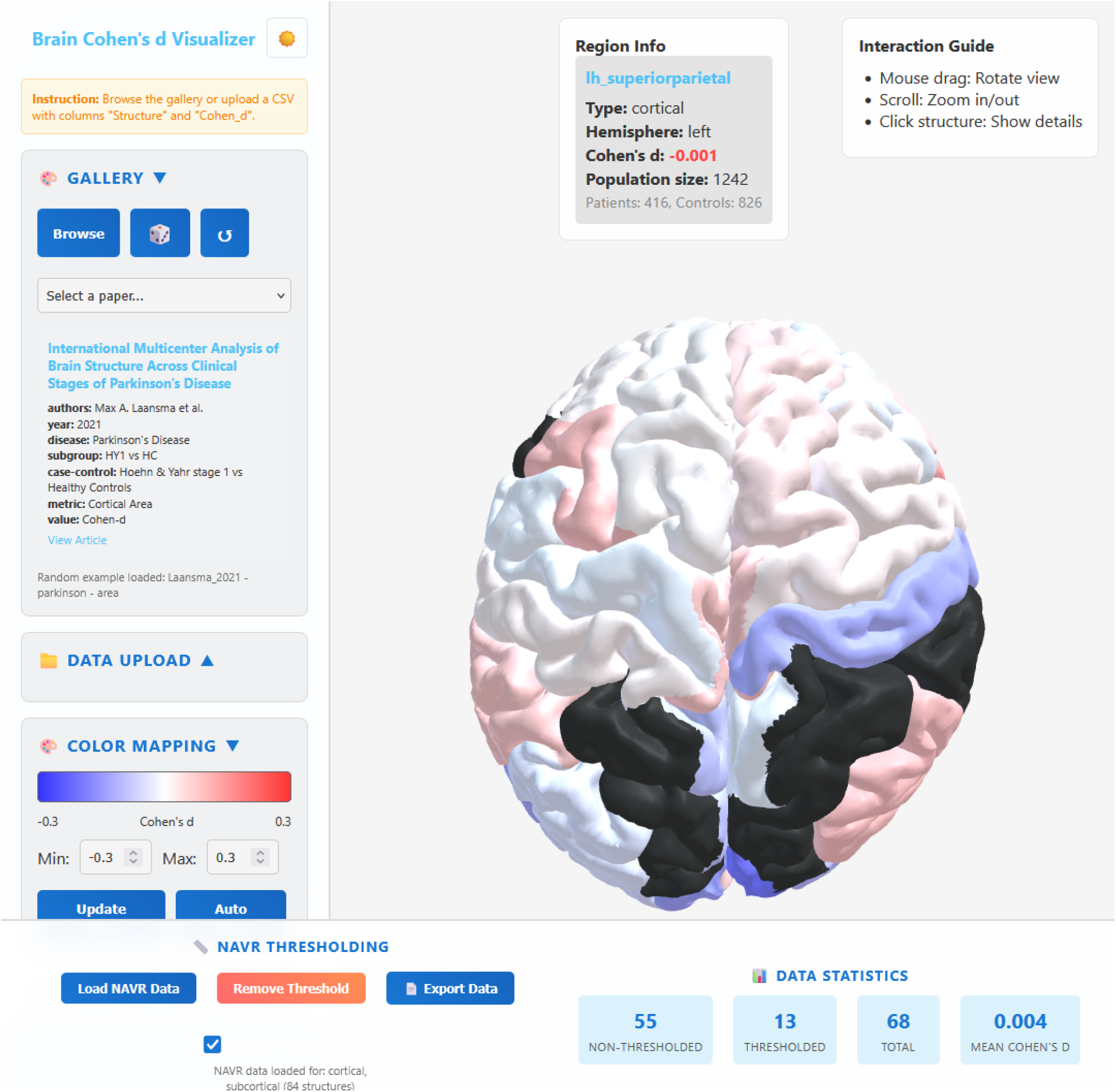
Interactive web tool for estimating NPVR and assessing numerical variability in neuroimaging studies. Users can input summary statistics to obtain NPVR values and visualize the impact of numerical variability on effect size estimates. The tool is available at yohanchatelain.github.io/brain render.

**Figure S8:**
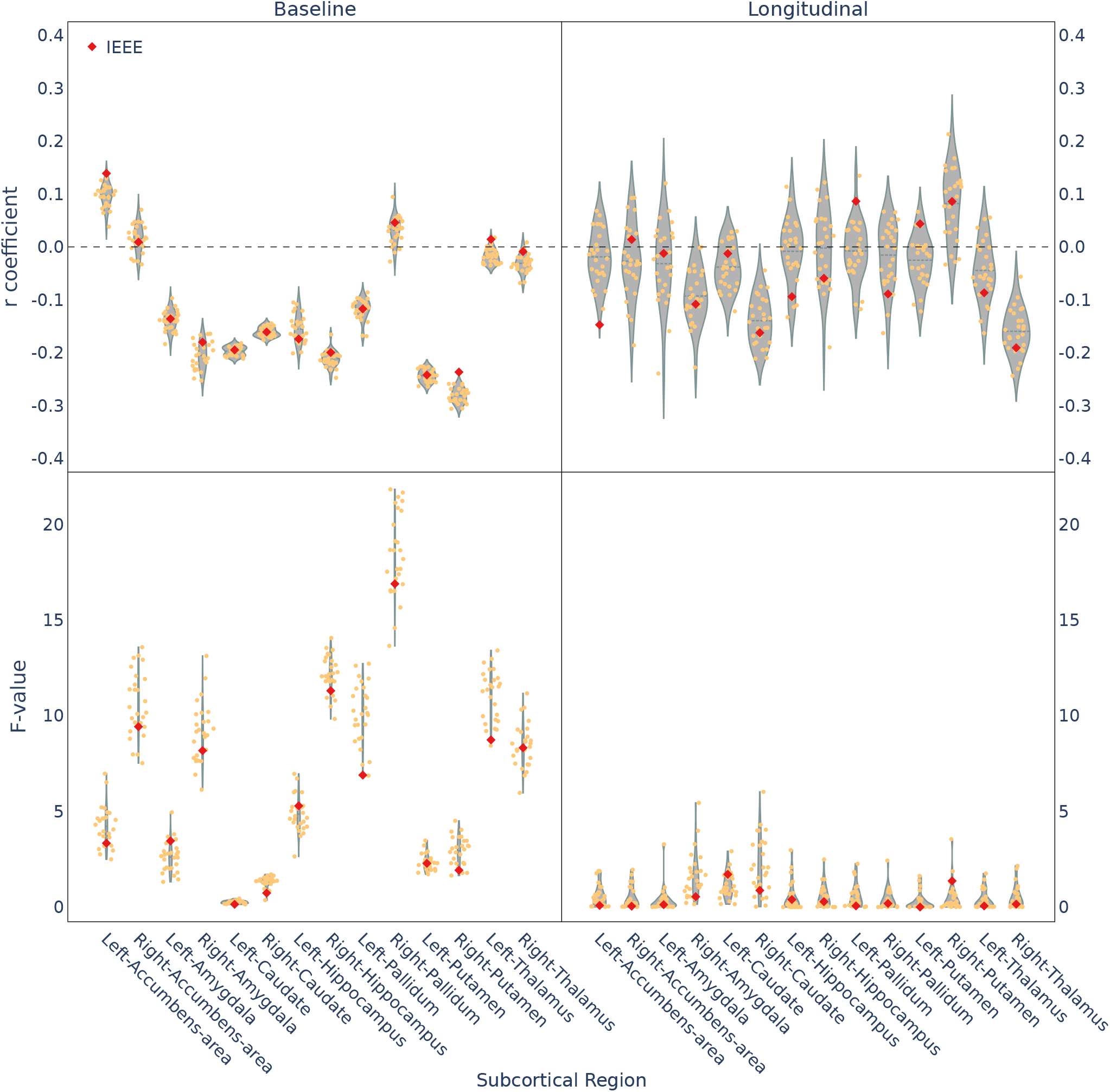
Distribution of partial correlation coefficients (r-values) and F-statistics from ANCOVA across MCA repetitions for subcortical volume measures. Red dots represent the IEEE-754 unperturbed results.

**Figure S9:**
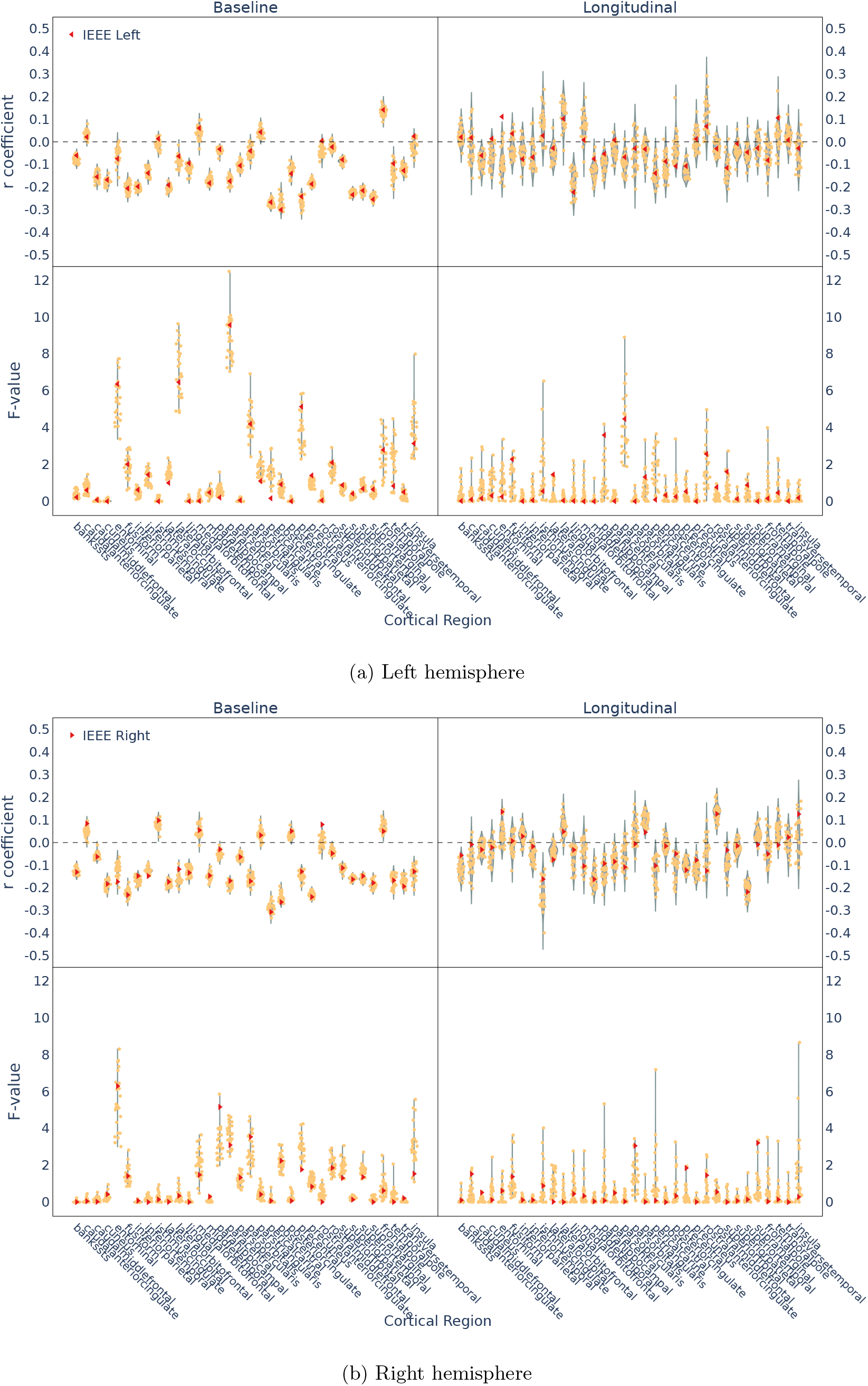
Distribution of partial correlation coefficients (r-values) and F-statistics from ANCOVA across MCA repetitions for cortical thickness measures. Red dots represent the IEEE-754 unperturbed results.

**Figure S10:**
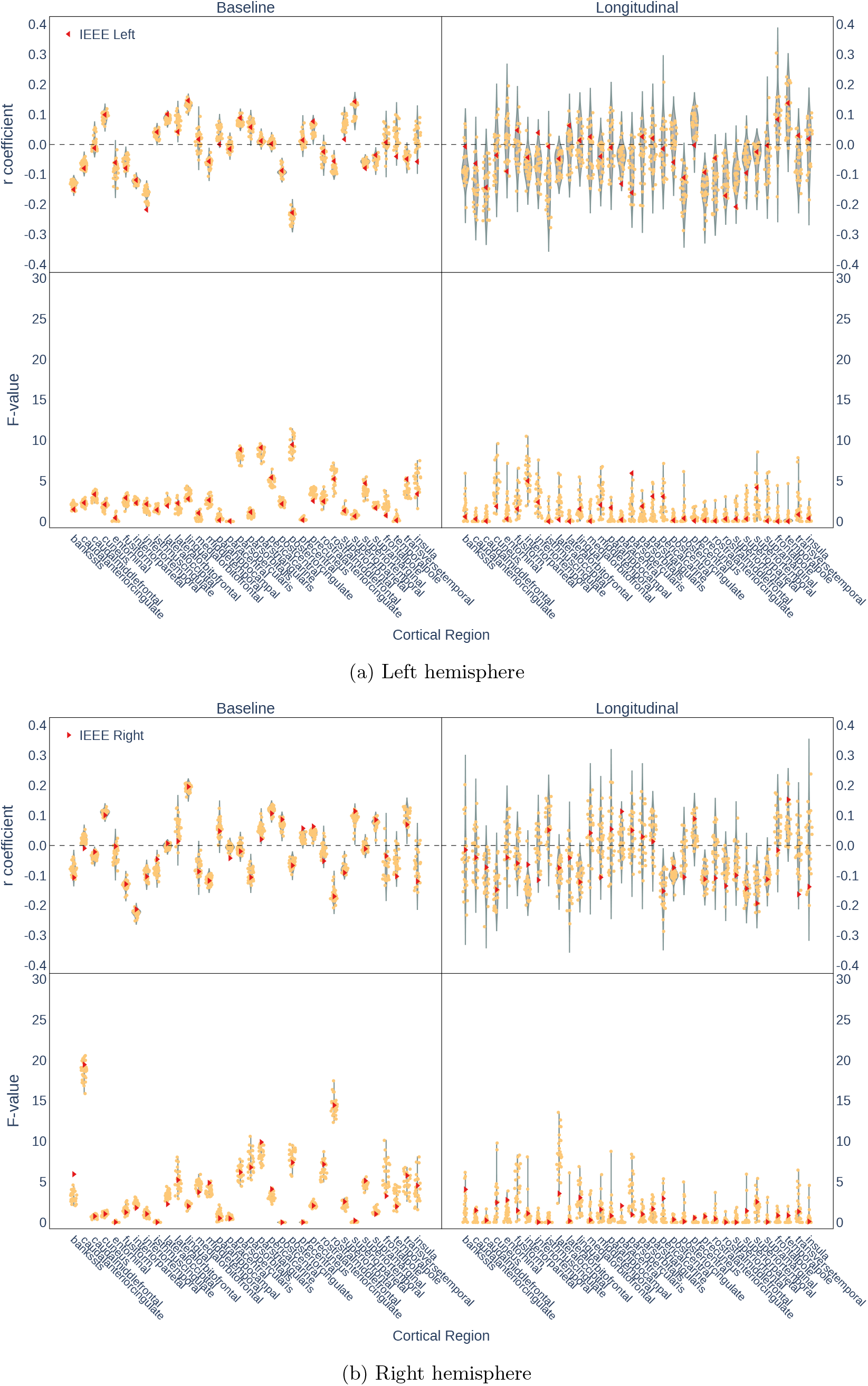
Distribution of partial correlation coefficients (r-values) and F-statistics from ANCOVA across MCA repetitions for cortical surface area measures. Red dots represent the IEEE-754 unperturbed results.

**Figure S11:**
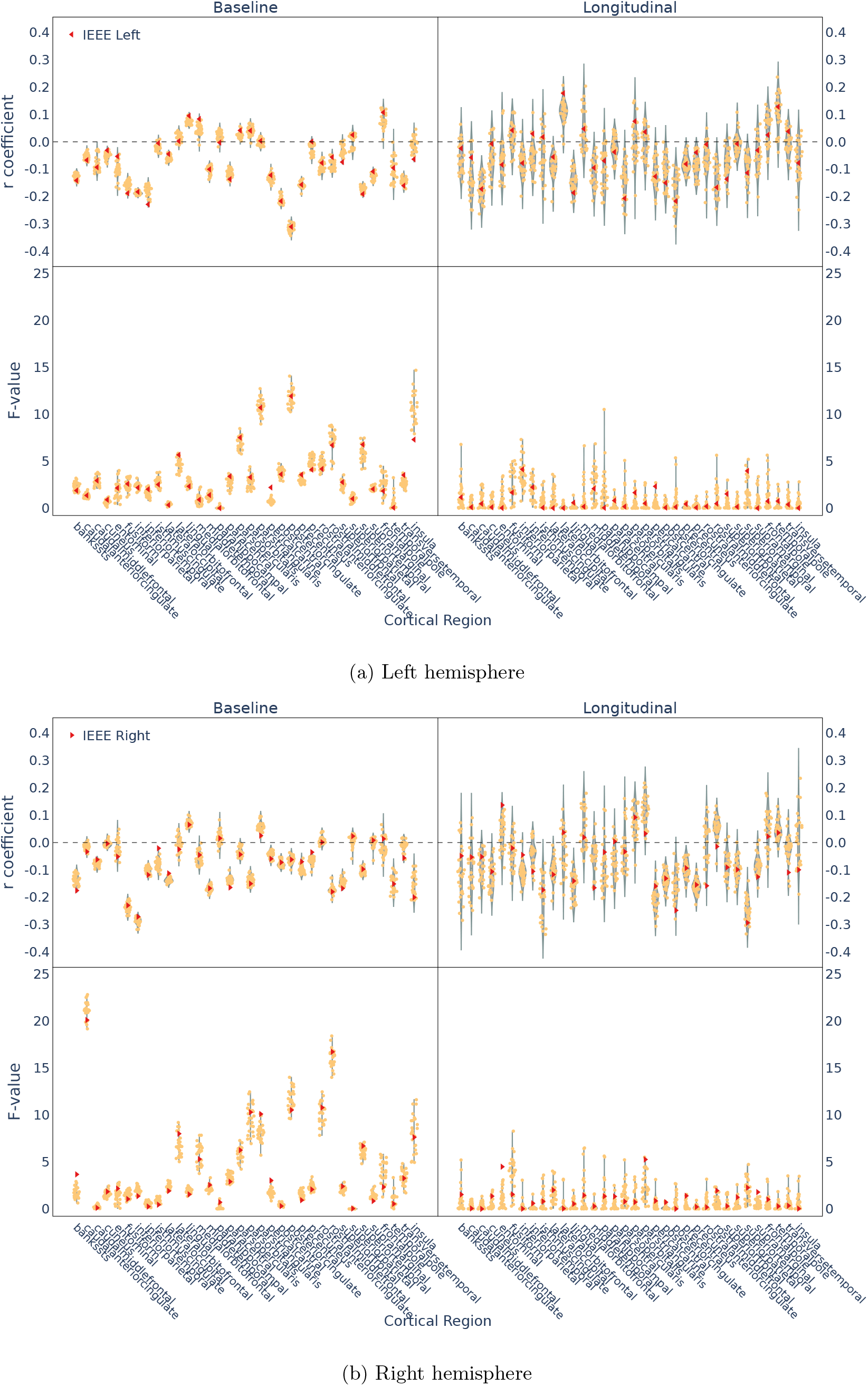
Distribution of partial correlation coefficients (r-values) and F-statistics from ANCOVA across MCA repetitions for cortical volume measures. Red dots represent the IEEE-754 unperturbed results.

**Figure S12:**
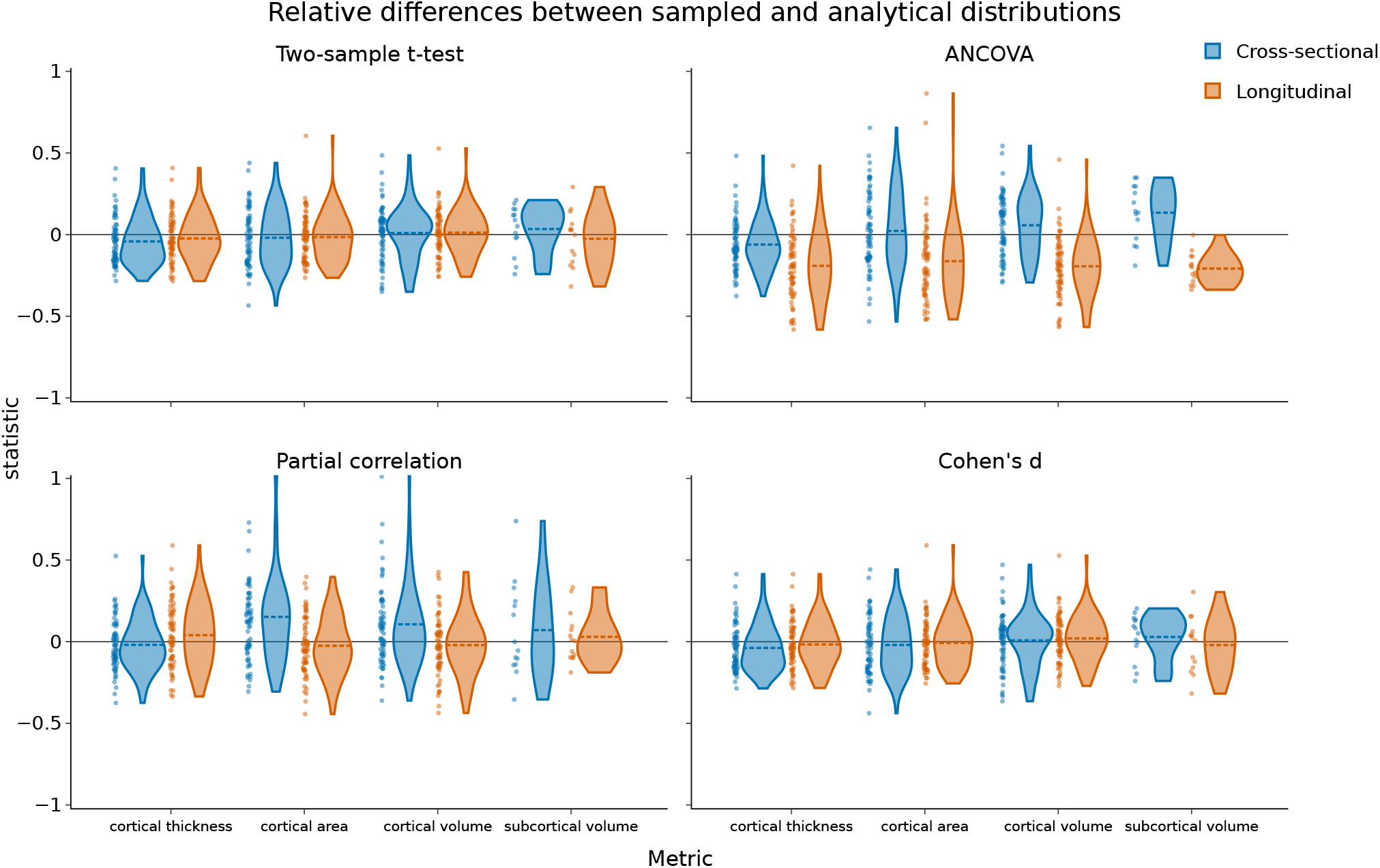
Relative error between analytical and sampled standard-deviation estimates of test statistics across all brain regions and metrics. Each violin plot corresponds to a specific statistic (Cohen’s *d*, two-sample *t*-test, partial correlation coefficient, ANCOVA *F* -statistic) and analysis type (cross-sectional or longitudinal). Distributions are generally concentrated within the interval [− 0.5, 0.5], with cross-sectional analyses remaining close to zero and longitudinal analyses exhibiting broader dispersion and a moderate negative shift for some statistics, most prominently ANCOVA.

**Figure S13:**
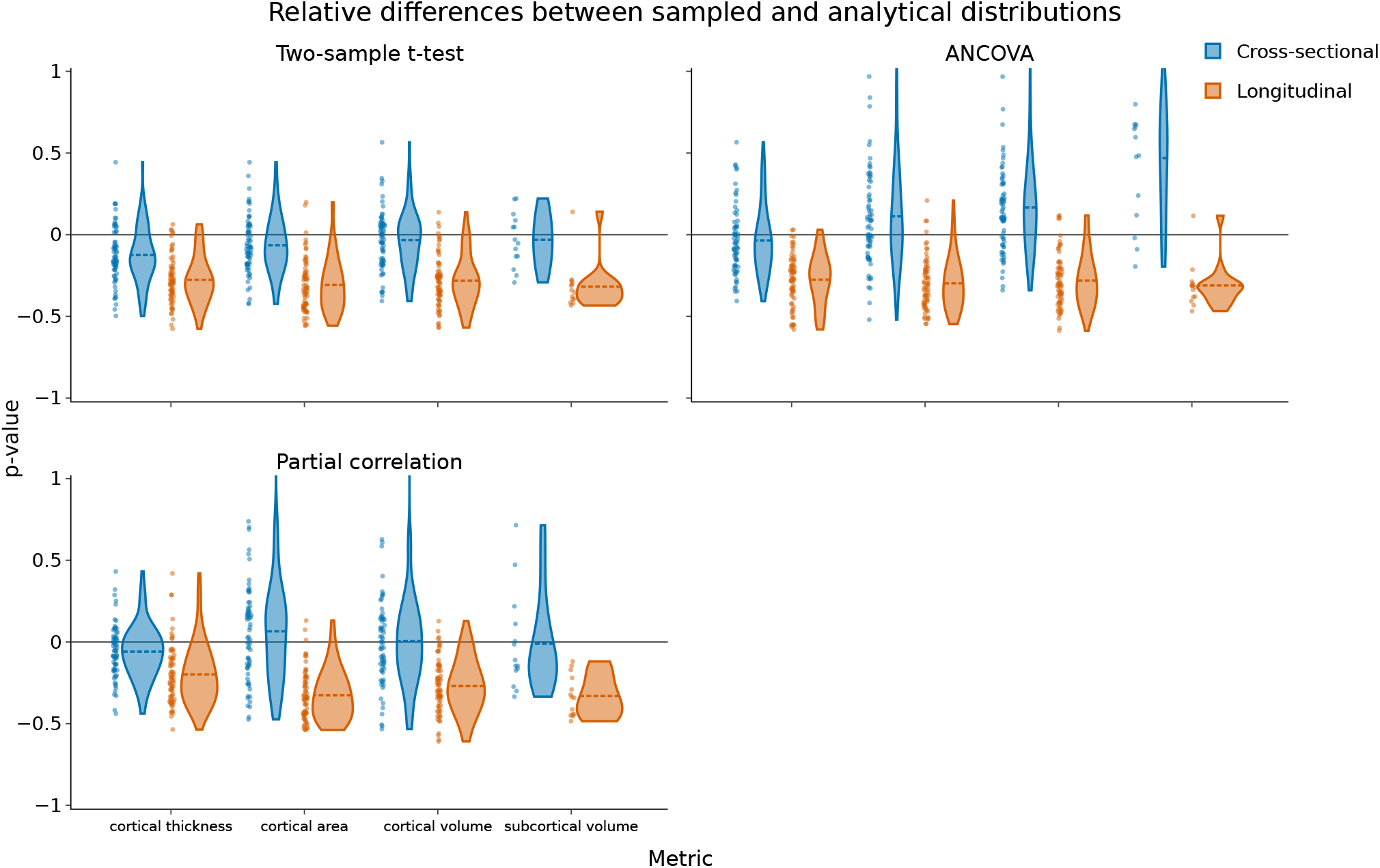
Relative error between analytical and sampled standard-deviation estimates of *p*-values across all brain regions and metrics. Each violin plot corresponds to a specific statistical test (two-sample *t*-test, partial correlation coefficient, ANCOVA *F* -statistic) and analysis type. Cross-sectional distributions remain centered near zero, whereas longitudinal analyses display wider spreads and systematic negative tendencies, particularly for cortical area and volume metrics.

**Figure S14:**
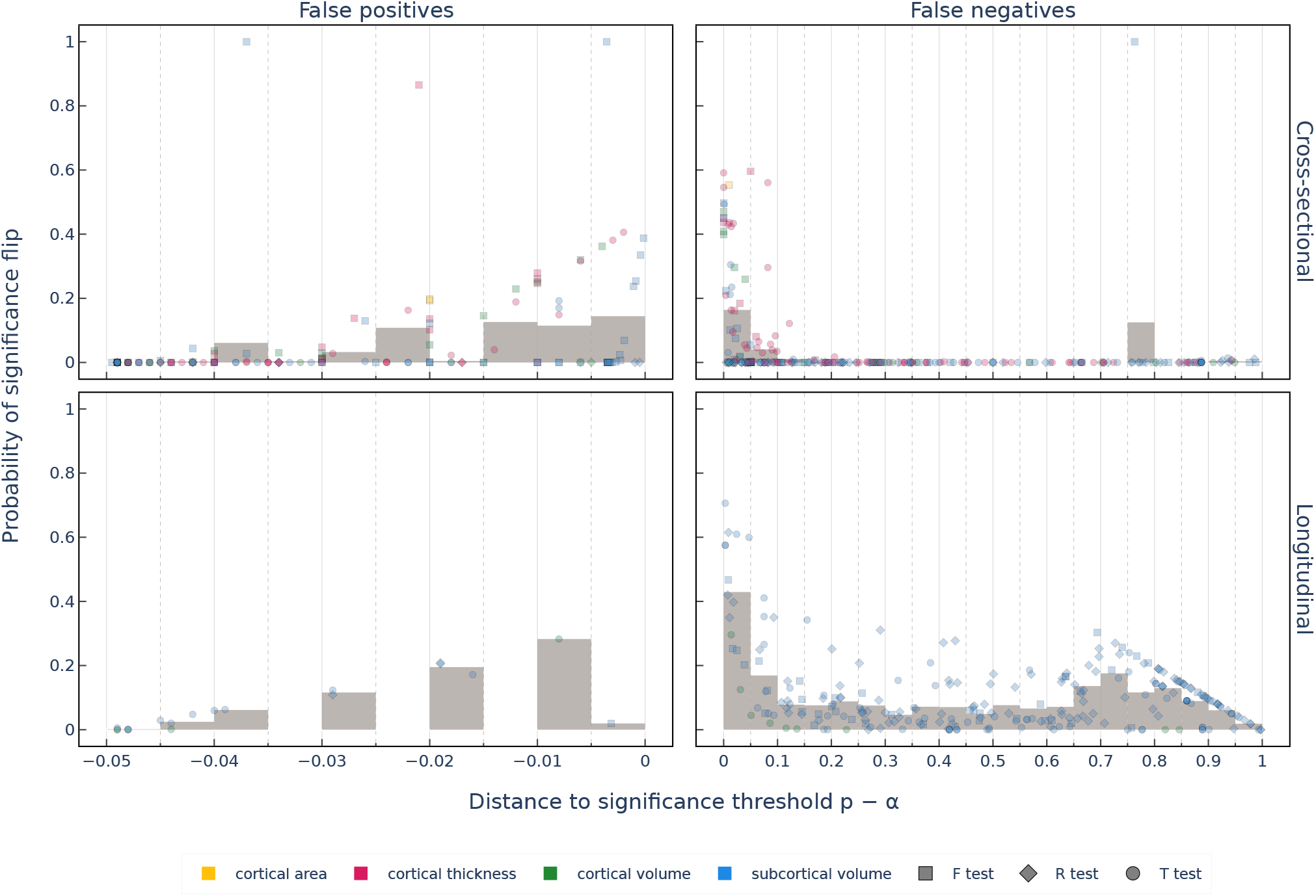
Probability of numerically-induced significance flips as a function of the *p*-value distance to significance threshold, broken down by statistical test type: Student’s *t*-test (T), ANCOVA (*F* -values), and partial correlation (R). Each point represents a result (significant or non-significant) reported in the 13 reviewed studies. Negative distances correspond to results reported as significant (*p < α*), for which the y-axis gives the probability of flipping to non-significant (false positive risk); positive distances correspond to results reported as non-significant (*p > α*), for which the y-axis gives the probability of flipping to significant (false negative risk). Bars indicate the mean significance flip probability within bins of distance to the threshold (e.g., 0.005 for false positive risk and 0.05 for false negative risk).

**Figure S15:**
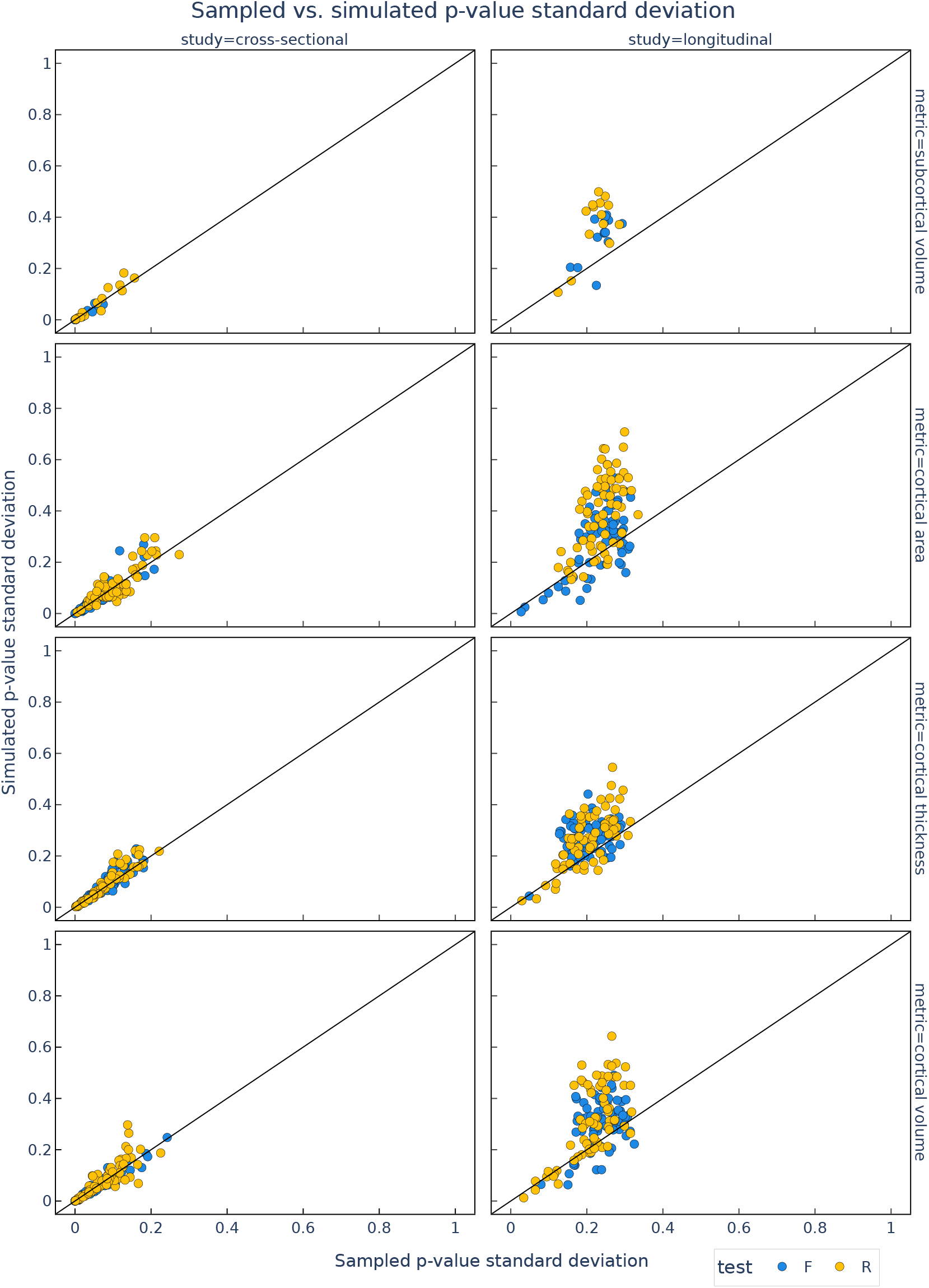
Comparison of standard deviation estimates of *p*-values obtained from MCA sampling and analytical simulations. Sampled estimates are computed from 26 MCA repetitions, while simulated estimates are derived from the variability propagation formulas reported in Table 1. Each point corresponds to a cortical or subcortical brain region. The solid line denotes the identity line. Results show close agreement between sampled and simulated estimates, with cross-sectional analyses exhibiting tighter clustering than longitudinal analyses. In the longitudinal setting, simulated standard deviations tend to slightly overestimate sampled variability, indicating conservative behavior of the analytical variability model.

**Figure S16:**
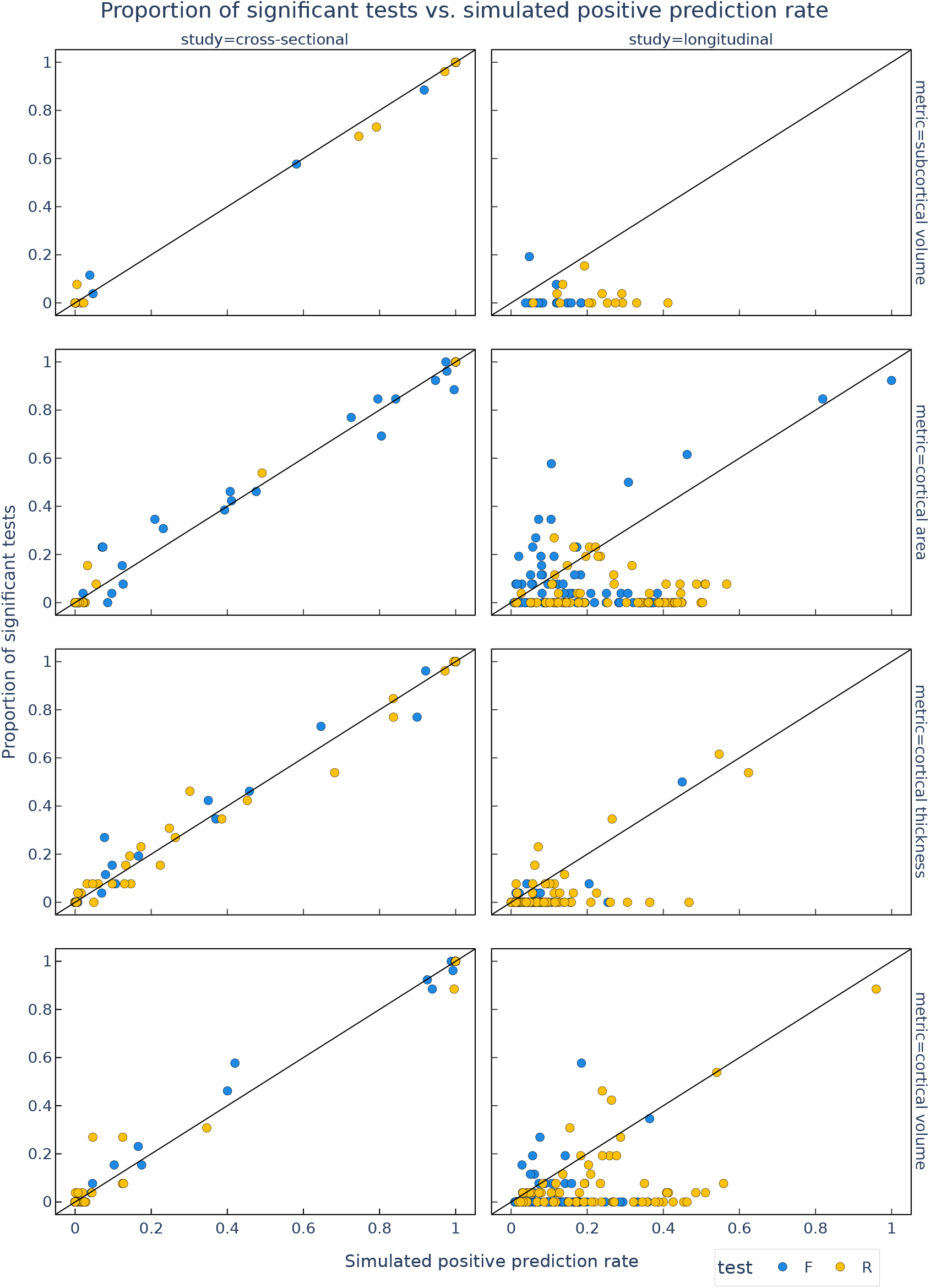
Comparison between simulated positive prediction rate (PPR; Eq. S10) and the empirical proportion of significant tests (Section 2.1). Simulated PPR values are computed using analytical variability estimates derived from Table 1. Each point represents a cortical or subcortical brain region, and the solid line indicates the identity line. Simulated PPR closely matches empirical significance rates in the cross-sectional setting, while longitudinal analyses show a tendency toward overestimation, consistent with the corresponding overestimation of *p*-value variability.

**Figure S17:**
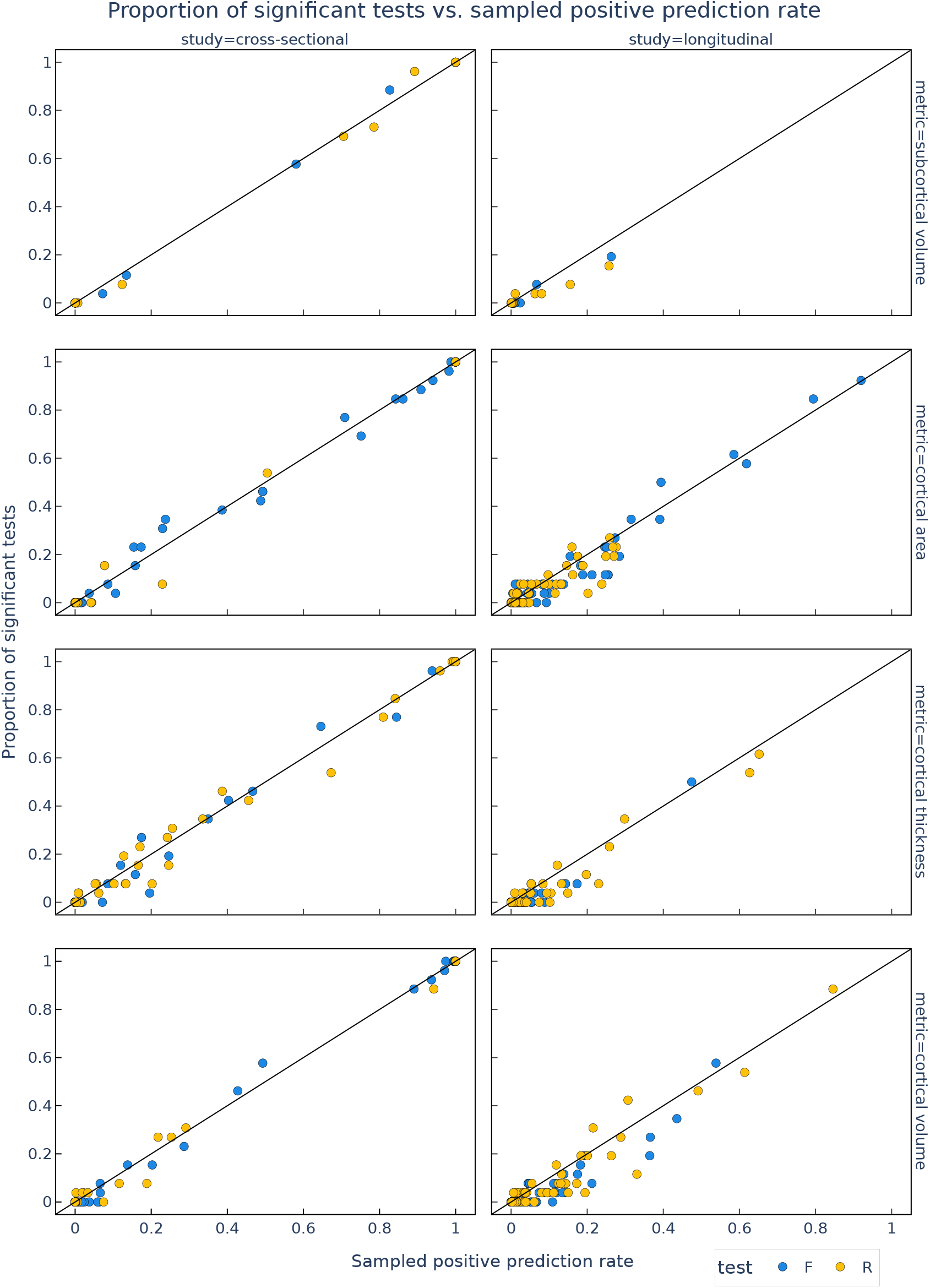
Comparison between sampled positive prediction rate (PPR; Eq. S10) and the empirical proportion of significant tests (Section 2.1). Sampled PPR values are computed from MCA-derived *p*-value standard deviations and propagated through the Beta distribution model (Eq. 4.4). Each point corresponds to a cortical or subcortical brain region, and the solid line denotes the identity line. Both cross-sectional and longitudinal results align closely with the identity line, supporting the suitability of the Beta modeling framework for capturing numerically induced variability in statistical significance.

## Footnotes

1 mean, variance, standard-deviation, covariance, Pearson correlation coefficient and correlation identities.

2 github.com/verificarlo/significantdigits

## Notes

https://github.com/yohanchatelain/livingpark-numerical-variability

https://yohanchatelain.github.io/brain_render/

### Competing Interest Statement

The authors have declared no competing interest.

### Summary of Updates

This version incorporates two rounds of peer review at Scientific Reports. Figure 4 was simplified and re-captioned, with the per-test-type breakdown moved to Supplementary Note S6. We added a metric-specific breakdown of numerical instability and confirmed that significance-flip risk is only weakly associated with sample size overall (r = -0.118), driven specifically by correlation-based tests (r = -0.417) rather than T/F-based tests. A “Practical recommendations” paragraph was added to the Discussion, and Figure 1 now shows 95% confidence intervals. In response to a second review round, we standardized terminology to a single consistent term ("numerical variability") throughout, replaced all inequality-only significance statements (p<0.05/p>0.05) with exact statistics, moved method-descriptive text out of the Results into Methods, added a new Methods subsection and table characterizing the design, sample size, and focus of the 13 retrospective studies, and substantially reorganized the Supplementary Information with a table of contents, a section-by-section roadmap linking each supplementary note to the main text, and corrected table captions.

## References

[1] Bhagwat, N. et al. Understanding the impact of preprocessing pipelines on neuroimaging cortical surface analyses. GigaScience 10, giaa155 (2021).

[2] Botvinik-Nezer, R. et al. Variability in the analysis of a single neuroimaging dataset by many teams. Nature 582, 84–88 (2020).

[3] Schilling, K. G. et al. Tractography dissection variability: What happens when 42 groups dissect 14 white matter bundles on the same dataset? Neuroimage 243, 118502 (2021).

[4] Kennedy, D. N. et al. Everything matters: the repronim perspective on reproducible neuroimaging. Frontiers in neuroinformatics 13, 1 (2019).

[5] Kiar, G. et al. Why experimental variation in neuroimaging should be embraced. Nature communications 15, 9411 (2024).

[6] Gronenschild, E. H. et al. The effects of freesurfer version, workstation type, and macintosh operating system version on anatomical volume and cortical thickness measurements. PloS one 7, e38234 (2012).

[7] Glatard, T. et al. Reproducibility of neuroimaging analyses across operating systems. Frontiers in neuroinformatics 9, 12 (2015).

[8] Markstein, P. The new ieee-754 standard for floating point arithmetic. Technical Report, Schloss Dagstuhl–Leibniz-Zentrum für Informatik (2008). Lecture /tutorial on the IEEE-754 revision.

[9] Salari, A., Chatelain, Y., Kiar, G. & Glatard, T. Accurate simulation of operating system updates in neuroimaging using monte-carlo arithmetic. In Uncertainty for Safe Utilization of Machine Learning in Medical Imaging, and Perinatal Imaging, Placental and Preterm Image Analysis: 3rd International Workshop, UNSURE 2021, and 6th International Workshop, PIPPI 2021, Held in Conjunction with MICCAI 2021, Strasbourg, France, October 1, 2021, Proceedings 3, 14–23 (Springer, 2021).

[10] Kiar, G. et al. Numerical uncertainty in analytical pipelines lead to impactful variability in brain networks. PloS one 16, e0250755 (2021).

[11] Des Ligneris, M. et al. Reproducibility of tumor segmentation outcomes with a deep learning model. In 2023 IEEE 20th International Symposium on Biomedical Imaging (ISBI), 1–5 (IEEE, 2023).

[12] Vila, G. et al. The impact of hardware variability on applications packaged with docker and guix: A case study in neuroimaging. In Proceedings of the 2nd ACM Conference on Reproducibility and Replicability, 75–84 (2024).

[13] Chatelain, Y. et al. A numerical variability approach to results stability tests and its application to neuroimaging. IEEE Transactions on Computers (2024).

[14] Mirhakimi, N., Chatelain, Y.Poline, J.-B. & Glatard, T. Numerical uncertainty in linear registration: An experimental study. arXiv preprint arXiv:2508.00781 (2025).

[15] Gonzalez-Pepe, I., Sivakolunthu, V., Chatelain, Y. & Glatard, T. Uncertain but useful: Leveraging cnn variability into data augmentation. arXiv preprint arXiv:2509.05238 (2025).

[16] Marek, K. et al. The parkinson progression marker initiative (ppmi). Progress in neurobiology 95, 629–635 (2011).

[17] Parker, D. S. Monte Carlo arithmetic: exploiting randomness in floating-point arithmetic (Cite-seer, 1997).

[18] Chagas, M. H. N. et al. Neuroimaging of major depression in parkinson’s disease: Cortical thickness, cortical and subcortical volume, and spectroscopy findings. Journal of psychiatric research 90, 40–45 (2017).

[19] Garcia-Diaz, A. I. et al. Structural mri correlates of the mmse and pentagon copying test in parkinson’s disease. Parkinsonism & related disorders 20, 1405–1410 (2014).

[20] Gerrits, N. J. et al. Cortical thickness, surface area and subcortical volume differentially contribute to cognitive heterogeneity in parkinson’s disease. PloS one 11, e0148852 (2016).

[21] Hanganu, A. et al. Mild cognitive impairment is linked with faster rate of cortical thinning in patients with parkinson’s disease longitudinally. Brain 137, 1120–1129 (2014).

[22] Lewis, M. M. et al. The pattern of gray matter atrophy in parkinson’s disease differs in cortical and subcortical regions. Journal of neurology 263, 68–75 (2016).

[23] Li, J. et al. Cortical and subcortical morphological alterations in motor subtypes of parkinson’s disease. npj Parkinson’s Disease 8, 167 (2022).

[24] Mak, E., Bergsland, N., Dwyer, M., Zivadinov, R. & Kandiah, N. Subcortical atrophy is associated with cognitive impairment in mild parkinson disease: a combined investigation of volumetric changes, cortical thickness, and vertex-based shape analysis. American Journal of Neuroradiology 35, 2257–2264 (2014).

[25] Pellicano, C. et al. Morphometric changes in the reward system of parkinson’s disease patients with impulse control disorders. Journal of neurology 262, 2653–2661 (2015).

[26] Radziunas, A. et al. Brain mri morphometric analysis in parkinson’s disease patients with sleep disturbances. BMC neurology 18, 88 (2018).

[27] Sokolowski, A. et al. The impact of freesurfer versions on structural neuroimaging analyses of parkinson’s disease. bioRxiv 2024–11 (2024).

[28] Sokolowski, A. et al. Longitudinal brain structure changes in parkinson’s disease: A replication study. Plos one 19, e0295069 (2024).

[29] Wilson, H., Niccolini, F., Pellicano, C. & Politis, M. Cortical thinning across parkinson’s disease stages and clinical correlates. Journal of the neurological sciences 398, 31–38 (2019).

[30] Yang, W. et al. The longitudinal volumetric and shape changes of subcortical nuclei in parkinson’s disease. Scientific Reports 14, 7494 (2024).

[31] Jenkinson, M., Beckmann, C. F., Behrens, T. E., Woolrich, M. W. & Smith, S. M. Fsl. Neuroimage 62, 782–790 (2012).

[32] Avants, B. B., Tustison, N., Song, G. et al. Advanced normalization tools (ants). Insight j 2, 1–35 (2009).

[33] Ashburner, J. Computational anatomy with the spm software. Magnetic resonance imaging 27, 1163–1174 (2009).

[34] Torabi, M., Mitsis, G. D. & Poline, J.-B. On the variability of dynamic functional connectivity assessment methods. GigaScience 13, giae009 (2024).

[35] Billot, B. et al. Synthseg: Segmentation of brain mri scans of any contrast and resolution without retraining. Medical Image Analysis 86, 102789 (2023).

[36] Hoopes, A., Mora, J. S., Dalca, A. V., Fischl, B. & Hoffmann, M. Synthstrip: Skull-stripping for any brain image. NeuroImage 260, 119474 (2022).

[37] Hoffmann, M. et al. Synthmorph: learning contrast-invariant registration without acquired images. IEEE transactions on medical imaging 41, 543–558 (2021).

[38] Pepe, I. G., Sivakolunthu, V., Park, H. L., Chatelain, Y. & Glatard, T. Numerical uncertainty of convolutional neural networks inference for structural brain mri analysis. In International Workshop on Uncertainty for Safe Utilization of Machine Learning in Medical Imaging, 64–73 (Springer, 2023).

[39] Amrhein, V., Greenland, S. & McShane, B. Retire statistical significance. Nature 567, 305–307 (2019).

[40] Lefort-Besnard, J., Nichols, T. E. & Maumet, C. Statistical inference for same data meta-analysis in neuroimaging multiverse analyzes. Imaging Neuroscience 3 (2025).

[41] Shearer, H. et al. BrainEffeX: A web app for exploring fMRI effect sizes. Aperture Neuro 5 (2025).

[42] Denis, C., de Oliveira Castro, P. & Petit, E. Verificarlo: checking floating point accuracy through monte carlo arithmetic. In 2016 IEEE 23nd Symposium on Computer Arithmetic (ARITH) (2016).

[43] Johnson, N. L., Kotz, S. & Balakrishnan, N. Continuous univariate distributions, volume 2, vol. 2 (John wiley & sons, 1995).

[44] Glatard, T. et al. A virtual imaging platform for multi-modality medical image simulation. IEEE transactions on medical imaging 32, 110–118 (2012).

[45] Sohier, D. et al. Confidence intervals for stochastic arithmetic. ACM Transactions on Mathematical Software (TOMS) 47, 1–33 (2021).

